# Cell segmentation-free inference of cell types from *in situ* transcriptomics data

**DOI:** 10.1101/800748

**Authors:** Jeongbin Park, Wonyl Choi, Sebastian Tiesmeyer, Brian Long, Lars E. Borm, Emma Garren, Thuc Nghi Nguyen, Bosiljka Tasic, Simone Codeluppi, Tobias Graf, Matthias Schlesner, Oliver Stegle, Roland Eils, Naveed Ishaque

**Affiliations:** Digital Health Center, Berlin Institute of Health (BIH) and Charité Universitätsmedizin, Berlin, Germany; Faculty of Biosciences, Heidelberg University, Heidelberg, Germany; Division of Computational Genomics and System Genetics, German Cancer Research Center (DKFZ), Heidelberg, Germany; Department of Computer Science, Boston University, Boston, the United States of America; Allen Institute for Brain Science, Seattle, WA, USA; Division of molecular neurobiology, Department of medical biochemistry and biophysics, Karolinska Institutet, Stockholm, Sweden; Science for life laboratory, Stockholm, Sweden; Bioinformatics and Omics Data Analytics, German Cancer Research Center (DKFZ), Heidelberg, Germany; Genome Biology Unit, European Molecular Biology Laboratory (EMBL), Heidelberg, Germany; Health Data Science Unit, Heidelberg University Hospital, Heidelberg, Germany

**Author notes:** These authors contributed equally to this work (Jeongbin Park, Wonyl Choi). These authors jointly supervised the work (Roland Eils, Naveed Ishaque). **Contact information** Correspondence: Roland Eils and Naveed Ishaque.

**Keywords:** *In situ* transcriptomics, spatial cell-type calling, cell segmentation-free, multiplexed FISH, SSAM, osmFISH, mFISH, multiplexed smFISH, MERFISH, spatially resolved RNA profiling

## Abstract

Multiplexed fluorescence *in situ* hybridization techniques have enabled cell-type identification, linking transcriptional heterogeneity with spatial heterogeneity of cells. However, inaccurate cell segmentation reduces the efficacy of cell-type identification and tissue characterization. Here, we present a novel method called Spot-based Spatial cell-type Analysis by Multidimensional mRNA density estimation (SSAM), a robust cell segmentation-free computational framework for identifying cell-types and tissue domains in 2D and 3D. SSAM is applicable to a variety of *in situ* transcriptomics techniques and capable of integrating prior knowledge of cell types. We apply SSAM to three mouse brain tissue images: the somatosensory cortex imaged by osmFISH, the hypothalamic preoptic region by MERFISH, and the visual cortex by multiplexed smFISH. We found that SSAM detects regions occupied by known cell types that were previously missed and discovers new cell types.

## Introduction

The underlying transcriptional and spatial heterogeneity of cells gives rise to the plethora of phenotypes observed in cell types, tissues, organs, and organisms. Recent technological advances^1^ have seen the profound adoption of single-cell sequencing to unravel transcriptional heterogeneity in healthy and diseased tissues, and have subsequently given rise to international consortia such as the Human Cell Atlas (HCA)^2^. Such efforts would not be possible without computational frameworks supporting the analysis of single-cell sequencing data^3^. Linking this transcriptional heterogeneity with spatial heterogeneity of cells is a critical factor in understanding cell identity in the context of the tissue, for example, revealing the transcriptional basis of invasive cancer regions^4^ and highlighting the rich diversity of neuronal subtype expression and localization^5^. Recently developed multiplexed fluorescence in-situ hybridization^6–8^ and *in situ* mRNA tissue sequencing techniques^9–14^ have enabled the simultaneous measurement of multiple mRNAs in a spatial context.

Traditionally, mRNA molecules identified by *in situ* transcriptomics are assigned to cells and subsequently used for computing gene expression profiles of those cells^15–18^. Identification of cells relies on cell segmentation, a procedure demarcating the interior and exterior of the cell membranes, which relies on additional signals or landmarks obtained by staining nuclei^19^, cell membrane^20–22^, or total poly-A RNA^5,6^. However, accurate cell segmentation is difficult to achieve with current techniques due to tightly apposed or overlapping cells, uneven cell borders, varying cell and nuclear shapes, signal intensity variation, probe fluorescence emission efficiency variation, and tiling artifacts^23^. Such obstacles can result in detecting fewer cells or incorrect cell borders. Subsequent analysis would then be spatially restricted to inaccurately segmented cells and may mean that large portions of meaningful mRNA signals are discarded. This may result in incorrect cell-type signatures, incomplete cell-type maps, or missing rare cell types. Therefore, there is a need for robust cell segmentation-independent methods for identifying cell-type signatures, cell-type organization, and tissue domains from multidimensional mRNA expression data in complex tissues. These methods could be used for datasets lacking landmarks or to validate segmentation-based approaches.

Here we introduce a novel computational framework named Spot-based Spatial cell-type Analysis by Multidimensional mRNA density estimation (SSAM). In contrast to existing methods, SSAM departs from the spatial restriction of approaches based on cell segmentation and instead identifies cell types using mRNA signals in the image, without the need for prior cell segmentation. Furthermore, instead of labelling only segmented regions, our approach assigns cell-type labels to each pixel, ensuring a more complete picture of cell-type specific spatial heterogeneity.

We apply SSAM to three mouse brain tissue images obtained by different techniques: the somatosensory cortex (SSp) by osmFISH, the hypothalamic preoptic region (POA) by MERFISH, and the visual cortex (VISp) by multiplexed smFISH. With all three datasets, we demonstrate the robustness of SSAM in identifying 1) cell types *in situ*, 2) spatial distribution of cell types, 3) spatial relationships between cell types, and 4) tissue domains (e.g., cortical layers) based on the local composition of cell types without fine-tuning of parameters. We demonstrate that SSAM 1) correctly identifies the spatial distribution of known cell types in regions missed in the SSp by cell segmentation based methods for the osmFISH data ; 2) can analyze the POA MERFISH 3D data using the same parameters as for the 2D SSp osmFISH data without any extra adjustments of the settings; 3) identifies new and rare cell types in the VISp, multiplexed smFISH data.

## Results

### The SSAM computational framework

SSAM consists of 4 major steps (**Fig. 1**), namely 1) mRNA signal estimation and downsampling; 2) computation of cell-type signatures; 3) generation of a cell-type map; and 4) identification of tissue domains.

**Figure 1.**
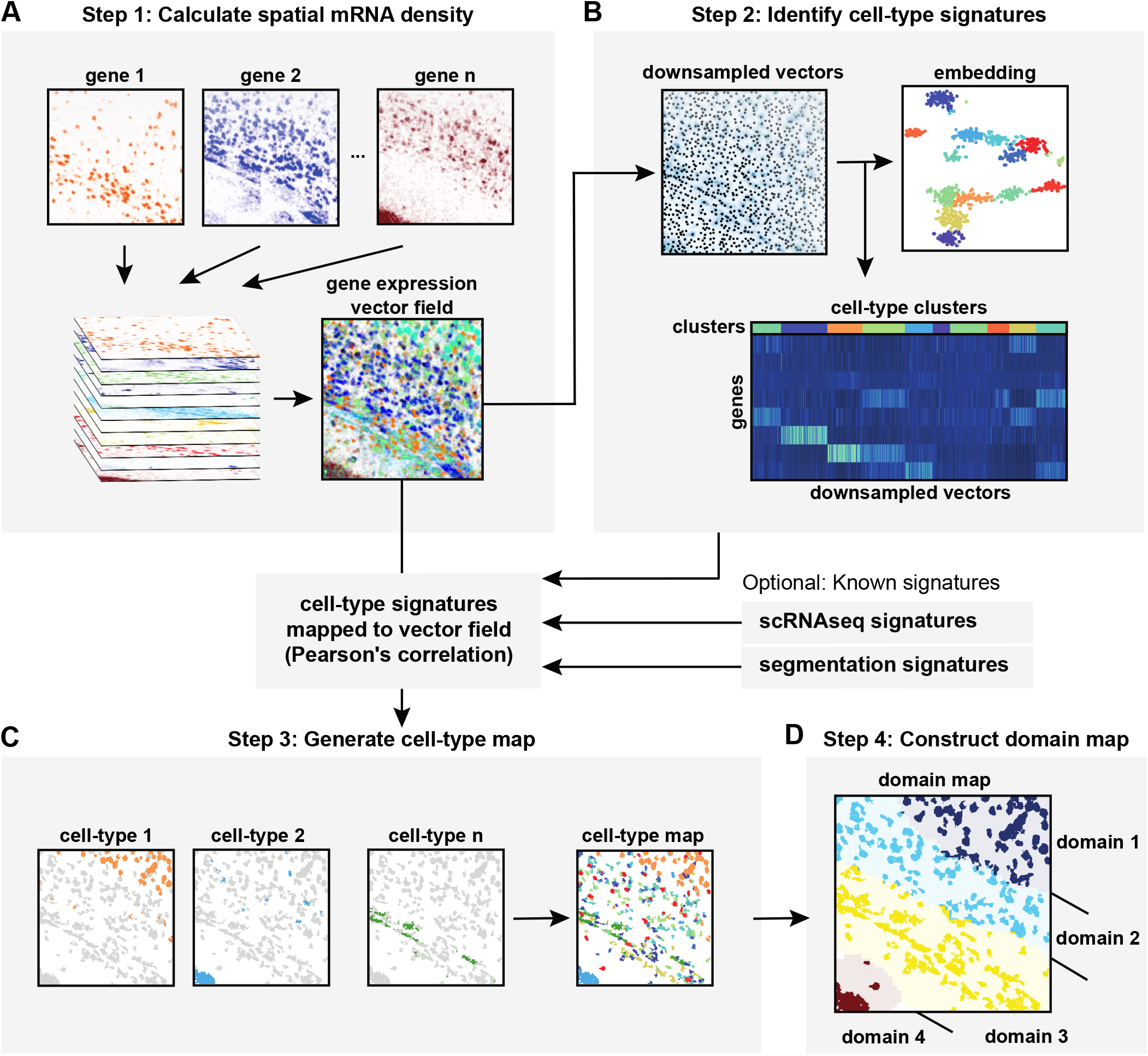
Schematic diagram of the SSAM computational workflow for cell type and tissue domain definition based on gene expression data. (A) In step 1, SSAM converts mRNA locations into a vector field of gene expression values. For this, SSAM applies a Gaussian KDE to mRNA locations for each gene and projects the resulting mRNA density values to a square lattice which represents coordinates in the tissue. The mRNA density estimated per each gene are stacked to produce a “gene expression vector field” over the lattice. The gene expression vector field is analogous to a 2D/3D image where each pixel/voxel encodes the averaged gene expression of the unit area. Further details of the application of KDE can be found in Supplementary Fig. 1A; (B) In step 2, cell-type signatures are identified *de novo*. First, the gene expression profile at probable cell locations are identified as the local regions in the gene expression vector field where the signal is highest. These downsampled gene expression signals are identified and used for *de novo* cell type identification by cluster analysis. Alternatively, previously defined cell-type signatures can be used. (C) In step 3, a cell-type map is generated. For this, the cell-type signatures are mapped onto the gene expression vector field and cell types are assigned based on Pearson’s correlation between each cell-type expression signature to the vector field to define cell-type distribution *in situ*. Further details about creating the cell-type map can be found in Supplementary Fig. 2A; (D) In step 4, the tissue domains are identified. The tissue domain signatures are identified using a sliding window to compute domain signatures based on the count of cell-type labels in the window. The tissue domains are defined by clustering these signatures. Further details on creating the tissue domain map can be found in Supplementary Fig. 2B.

In the first step, SSAM estimates mRNA signal intensity over the tissue image (**Fig. 1A**). Firstly, for each gene, mRNA signal intensity distribution is estimated by applying a Kernel Density Estimation (KDE) with a Gaussian kernel, which is then resolved to pixels in the image. The mRNA signal intensity distribution for each gene is stacked to create a gene expression vector field, which is a multichannel image where the pixels encode the expected density of mRNA count for each gene. This essentially assigns gene expression profiles to pixels in the image.

In the second step, SSAM identifies cell-type gene expression signatures by clustering (**Fig. 1B**). Before running the clustering algorithm, SSAM downsamples gene expression vectors to reduce computational processing time. As default, SSAM performs informed downsampling by selecting pixels that are local maxima in the gene vector field (Methods). After that, both the downsampled vectors and the gene expression vector field are normalized (Methods). SSAM clusters the sampled vectors using either DBSCAN^24^, HDBSCAN^25^, OPTICS^26^, or the Louvain community detection method implemented in Seurat^27^ (Methods). The Louvain methods is the default as it has been widely utilized to analyze single cell data. After the clustering step, sampled vectors with a large distance in gene expression space to their cluster medoid are removed as outliers to ensure the quality of selected vectors (**Supplementary Fig. 1B**). The gene expression cluster centroids are used to represent the gene expression signature of a cell type.

In the third step, SSAM classifies each pixel in the image to create a “cell-type map” (**Fig. 1C, Supplementary Fig. 2A**). SSAM includes a guided mode, which assigns pixels to a labeled set of given gene expression signatures (e.g. from scRNA-seq/segmentation), as well as a *de novo* mode, which assigns pixels to the cell type signatures obtained in the previous clustering step. For the classification of pixels, SSAM first creates signature prototypes by averaging the signatures per cell-type class of the given signatures, then it classifies all spots in the vector field according to the maximum correlation to any of the signature prototypes.

**Figure 2.**
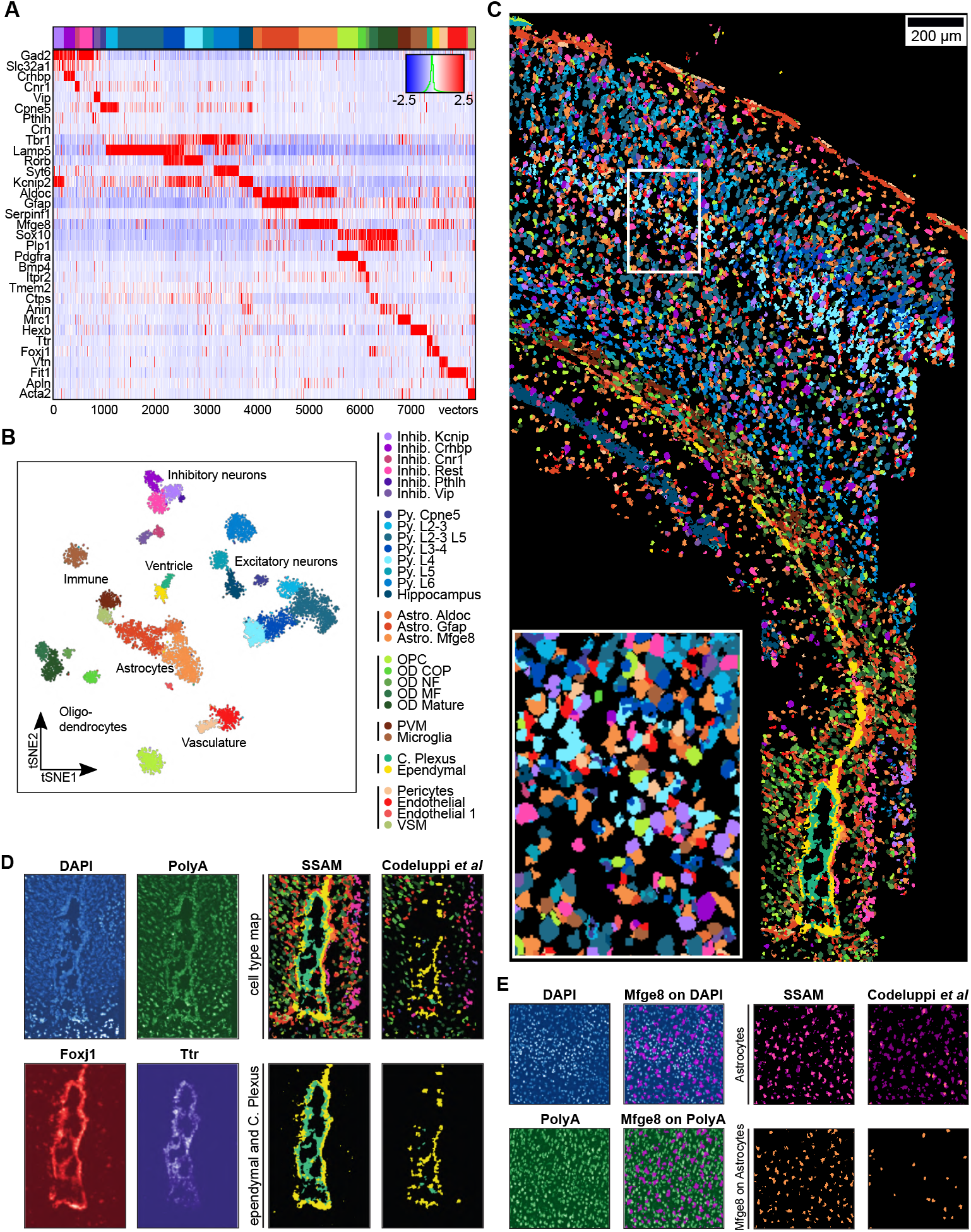
SSAM improves astrocyte and ventricle detection in the mouse SSp region. (A) Gene expression heatmap showing cell-type specific expression of marker genes (8,252 vectors). Rows show z-score normalized gene expression and columns show the gene expression patterns of filtered local maxima vectors. The top annotation shows the cell types and coloring based on the best correlating segmentation-based cell-type signature from Codeluppi *et al*. The colors of the top annotation correspond to the cell type legend in Fig. 2B; (B) A t-SNE map of cell-type signatures with distinct expression. Cell-type clusters are visualized as a 2D t-SNE embedding of filtered local maxima vectors. Cell-type annotation and coloring are based on the best correlating segmentation-based cell-type signature from Codeluppi *et al* (Supplementary Fig. 4C,D). The cell-type legend is grouped by cell-type classes labels shown in the tSNE plot, and are based on groupings by Codeluppi *et al*.; (C) The SSAM *de novo* cell-type map showing spatial organization of the cell types signatures in the gene expression vector field. Inset shows a zoom in of the highlighted tissue region. The colors of the cell types correspond to the cell-type legend in Fig. 2B; (D) SSAM improves the reconstruction of the ventricle. The upper left 2 panels show the DAPI and Poly-A signal around the ventricle area, showing tightly packed cells (occlusion) and lower signal in the ventricle structure compared to surrounding cells. The lower left 2 panels show the KDE gene expression signature for *Foxj1* (the marker for ependymal cells) and *Ttr* (the marker for choroid plexus cells). The upper right 2 panels show the cell-type maps reconstructed by SSAM, showing a more complete reconstruction, and by Codeluppi *et al*., which misses parts of the ventricle structure. The bottom right 2 panels show the reconstructions of only the ependymal (yellow) and choroid plexus (teal) cell types by SSAM and Codeluppi *et al*.; (E) SSAM has increased sensitivity of astrocyte detection. The far left upper and lower panels show DAPI and Poly-A signal for a region in the tissue. The middle left upper and lower panels show the overlap of *Mfge8* signal (a marker for one astrocyte) with DAPI and Poly-A signals, showing that *Mfge8* signal corresponds with low Poly-A signal, but with higher DAPI signal. The top right 2 panels show the cell-type signals for *Mfge8* expressing astrocytes by SSAM and Codeluppi *et al*., showing that SSAM detect much more astrocyte cell types. The bottom right 2 panels shows the overlay of *Mfge8* signal with the cell-type calls by SSAM and Codeluppi *et al*., showing the astrocyte signals detected by SSAM correspond well with *Mfge8* signal.

In the fourth step, SSAM identifies tissue domains that have distinct cell-type composition (**Fig. 1D**). SSAM computes the cell-type compositions in a circular (or spherical) sliding window over the cell-type map and clusters the cell-type composition of each window using agglomerative hierarchical clustering (**Supplementary Fig. 2B**). The resultant clusters represent putative tissue domains. Clusters with high mutual correlation are then merged into a single tissue domain signature, and the cell-type composition of each domain is calculated.

In the following sections we apply SSAM to three multiplexed FISH datasets obtained using different techniques. We reanalyze two previously published datasets, profiled by osmFISH^6^ and MERFISH^5^, to demonstrate SSAM’s strength in comparison to earlier methods. For a newly generated multiplexed smFISH dataset we demonstrate that SSAM can unravel novel biological insights into the spatial cellular organization of the brain.

### SSAM improves astrocyte and ventricle detection in the mouse brain somatosensory cortex (SSp)

To demonstrate the utility of SSAM, we analyzed published osmFISH data, where the transcripts of 33 cell-type marker genes were localized in 2D space of the mouse brain somatosensory cortex (SSp)^6^ (**Fig. 2, 3, Supplementary Fig. 3,4**). We compare results obtained from SSAM against the results obtained from Poly-A segmentation from the original study.

**Figure 3.**
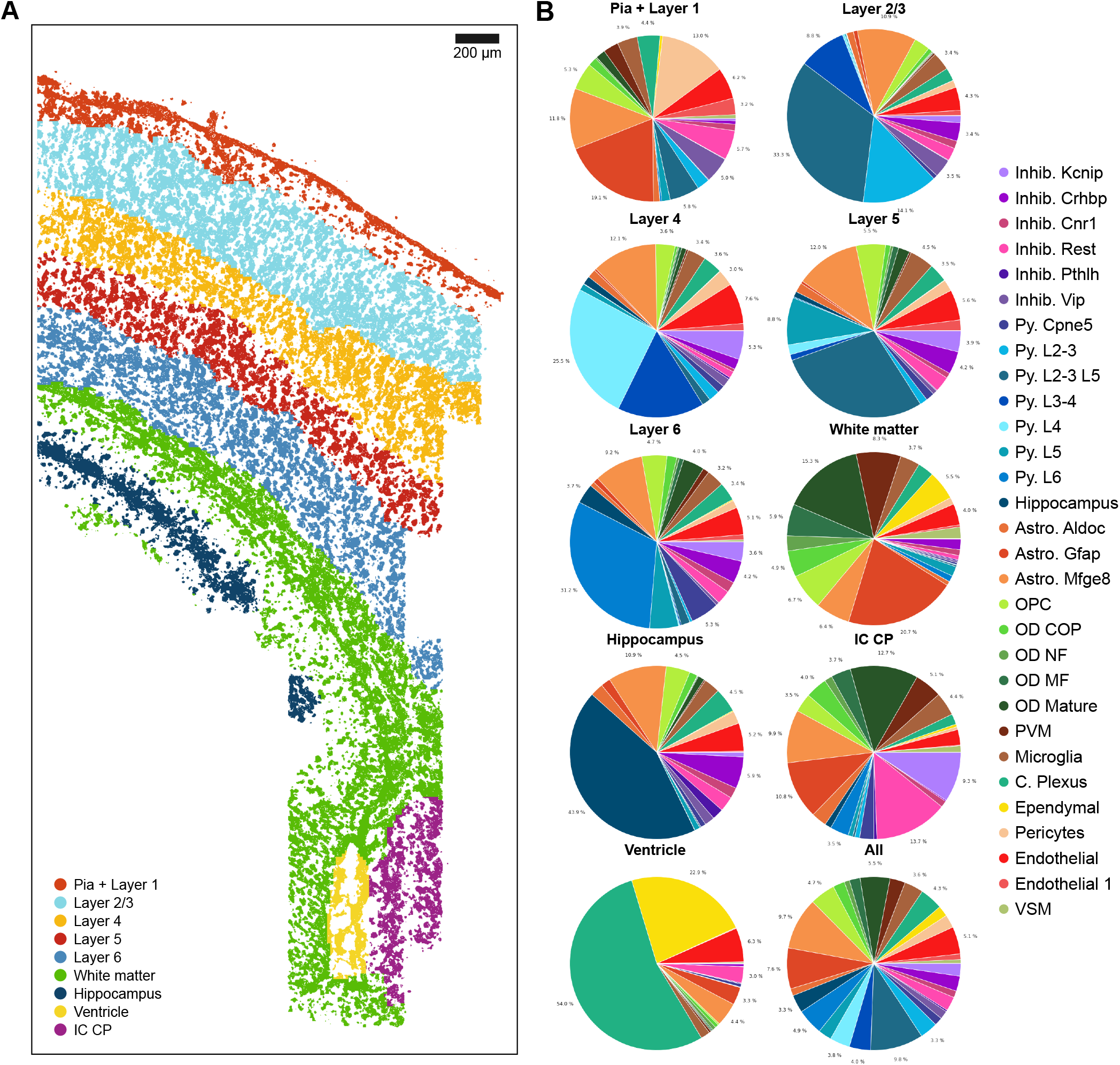
SSAM identifies cortical layer tissue domains in the mouse SSp cortex. (A) Tissue domain map generated by SSAM. Tissue domain signatures were identified from clustering local cell-type composition over sliding 100 μm circular windows, and projected back onto the cell-type map. The reconstruction shows the various cortical layers; (B) Cell-type composition within each tissue domain. The plots show that each domain consists of 7-14% Astrocyte Mfge8 cell types, apart from the ventricle, which instead shows a majority of Choroid plexus and Ependymal cell types.

**Figure 4.**
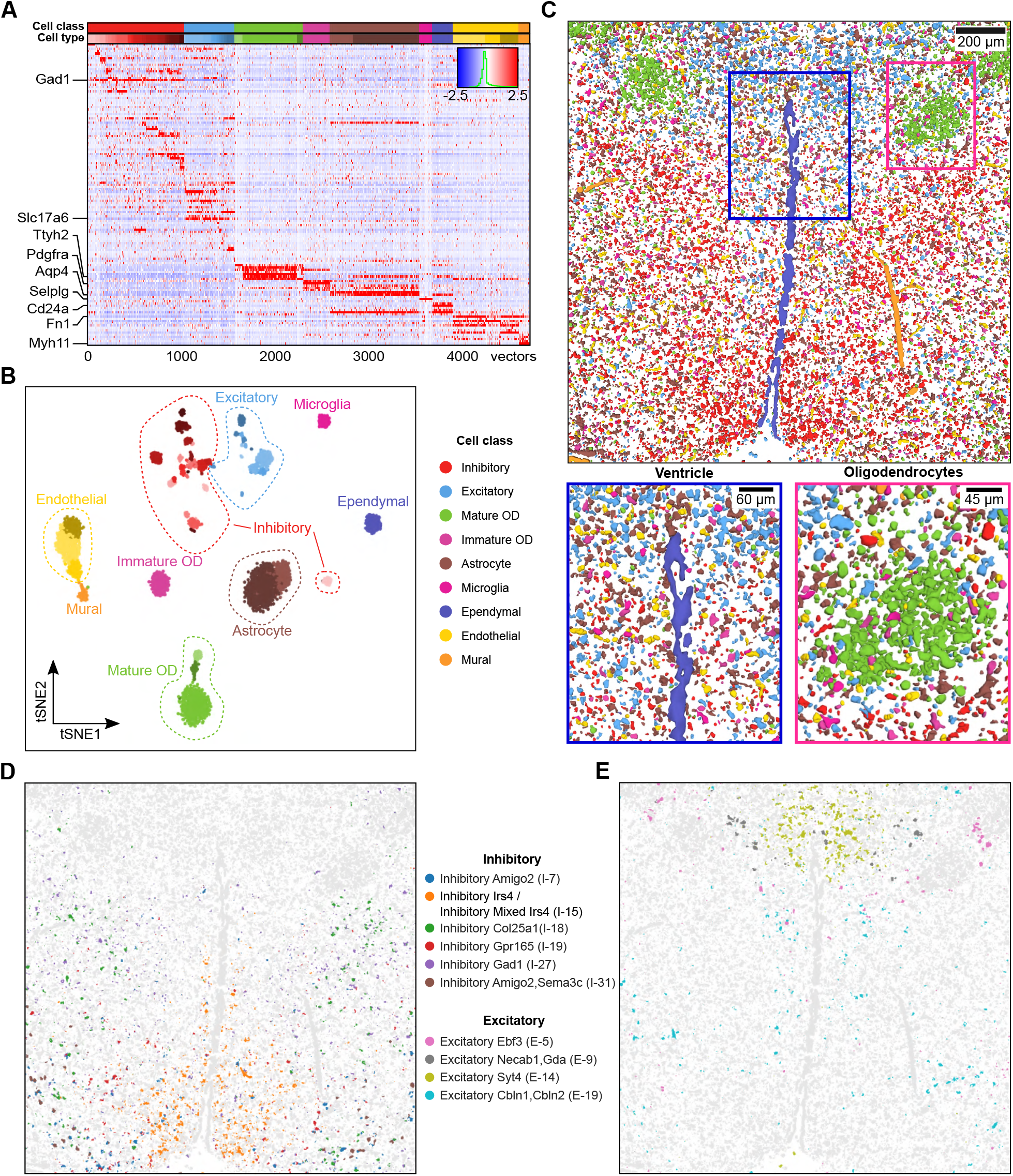
SSAM 3D cell type map confirms rich diversity of heterogeneous cells in the posterior hypothalamic POA. (A) Gene expression heatmap showing cell-type specific expression of marker genes (4,714 vectors). Rows show z-score normalized gene expression and columns show the gene expression patterns of filtered local maxima vectors (representative of gene expression within a cell). The bottom row of the top annotation shows the cell types. Due to a rich diversity of various inhibitory and excitatory neurons captured, the cell types were grouped into classes. The top row of the top annotation shows the cell classes which are named and colored based on the best cell-type signatures and cell classes from Moffitt *et al*. The colors of the cell classes top annotation correspond to the cell-type legend in Fig. 4B. The colors of the cell types are available in Supplementary Fig. 16; (B) A tSNE map of cell-type signatures with distinct expression. Cell-type clusters are visualized as a 2D t-SNE embedding of filtered local maxima vectors. Cell-type annotation and coloring are based on the best correlating segmentation-based cell-type signature from Moffitt *et al*. The tSNE map clearly shows the distinct cluster of different inhibitory and excitatory cell-type signatures. Cell types are grouped into classes based on groupings by Moffitt *et al*.; (C) The SSAM *de novo* 3D cell-type map showing spatial organization of the cell types signatures in the gene expression vector field. Below left and right a zoom in of the highlighted tissue regions of the ventricle structure and clusters of oligodendrocyte cell types. The colors of the cell types correspond to the cell-type legend in Fig. 4B; (D) Spatial localization of various inhibitory cell-type signatures. We found a number of inhibitory cell types which both matched expression signature and tissue localization described by Moffitt *et al*. See also Supplementary Fig. 22; (E) As panel D, but for excitatory cell types.

The osmFISH dataset was first analyzed using the guided mode of SSAM. Cell-type maps were generated using cell-type signatures from the prior segmentation-based approach^6^ and another from scRNA-seq^28,29^ (**Supplementary Fig. 4E**).

To quantify the similarity between the prior segmentation and the cell-type maps generated by SSAM, we calculated a “matching score” for each cell type (Methods). The matching scores between the segmentation from the previous study and SSAM guided by both segmentation-based and scRNA-seq cell-type signatures were generally high (mean and median matching score of 0.67 and 0.78 for segmentation-based, 0.60 and 0.70 for scRNA-seq-based signatures, respectively), indicating a strong agreement of the two cell-type maps as visually apparent (**Supplementary Table 1, 2, Supplementary Fig. 5,6**).

**Figure 5.**
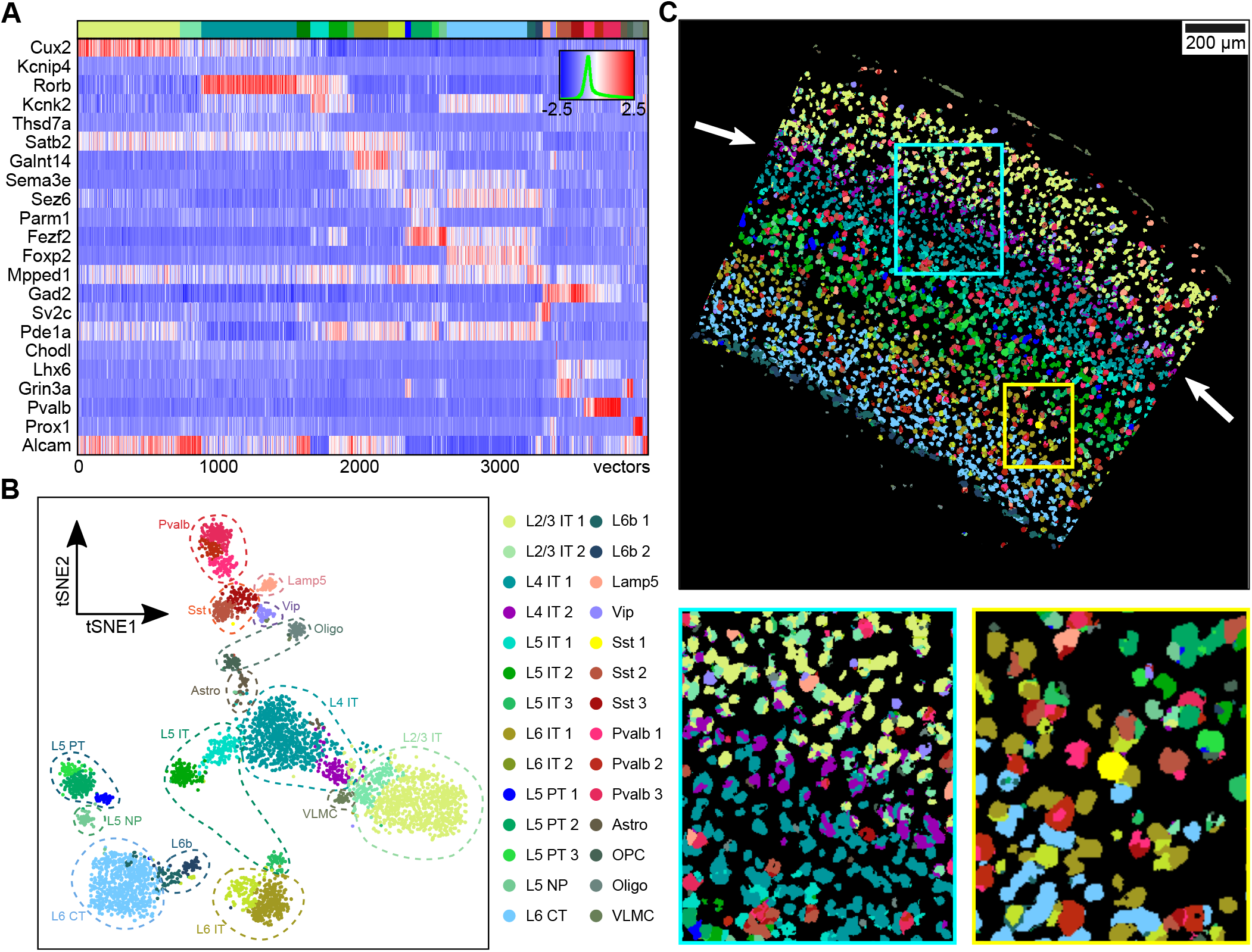
SSAM identifies layer structure in VISp and confirms rare Sst Chodl cell type in the mouse VISp region. (A) Gene expression heatmap showing cell-type specific expression of marker genes (4,113 vectors). Rows show z-score normalized gene expression and columns show the gene expression patterns of filtered local maxima vectors. The top annotation shows the cell types and coloring based on the highest correlating single cell RNA-seq based cell-type signature from previous result (Tasic *et al*., 2018). The colors of the top annotation correspond to the cell-type legend in Fig. 5B; (B) A tSNE map of cell-type signatures with distinct expression. Cell-type clusters are visualized as a 2D t-SNE embedding of filtered local maxima vectors, with groupings based on the supplementary table 9 of Tasic et al. 2018. Cell-type annotation and coloring are based on the best correlating segmentation-based cell-type signature from previous result (Tasic *et al*., 2018); (C) The SSAM *de novo* cell-type map showing spatial organization of the cell types. Highlighted are the tissue regions of the cortex including novel L4 IT cell type sub-layering (main panel, purple, white arrows, lower left panel, see also Supplementary Fig. 29B), and rare Sst Chodl cell type (lower right panel, yellow, see also Supplementary Fig. 29C). The colors of the cell types correspond to the cell-type legend in Fig. 5B.

**Figure 6.**
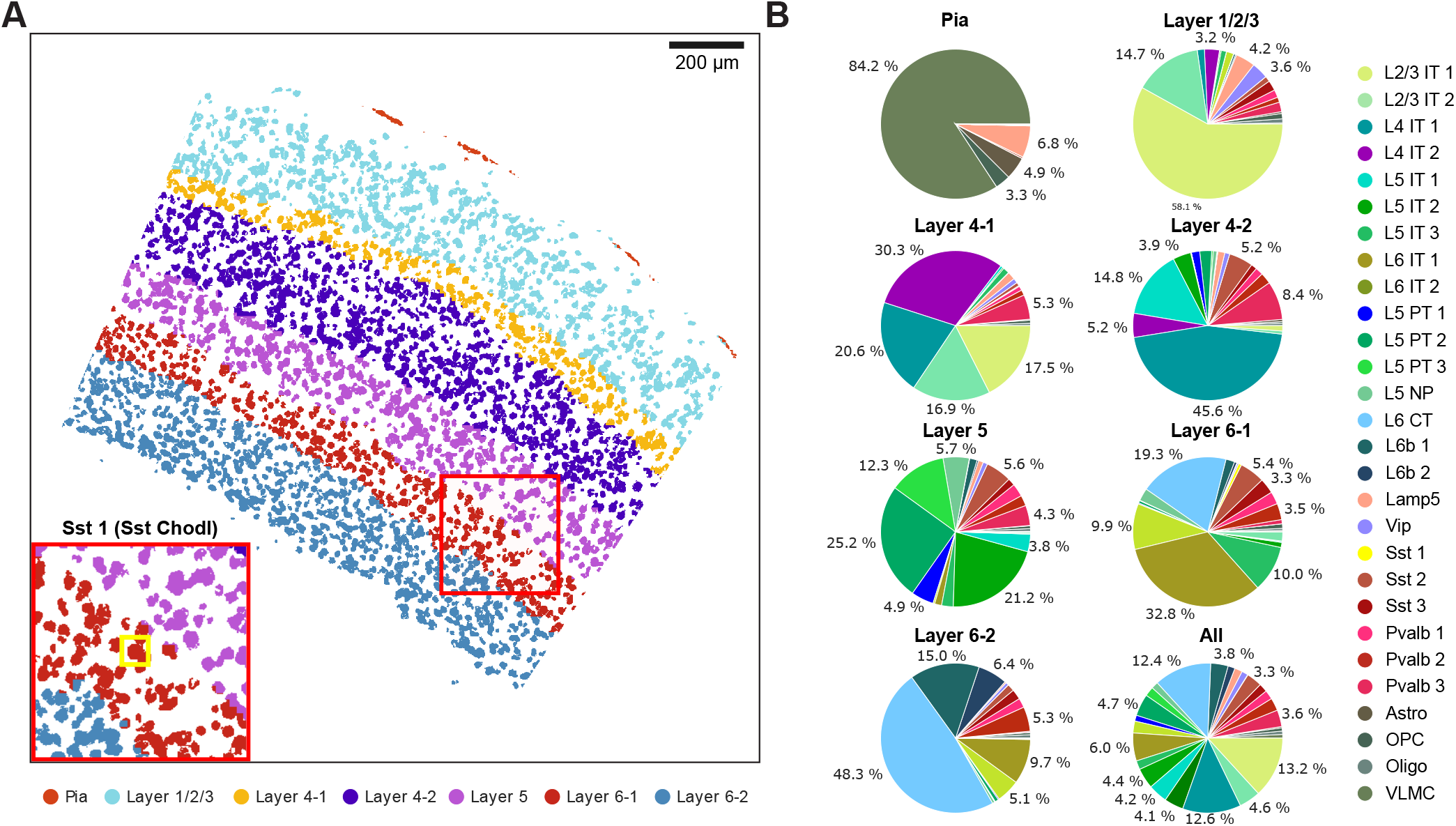
Rare Sst Chodl cell type localizes to the L6-1 layer of the mouse VISp region. (A) Tissue domain map generated by SSAM. Tissue domain signatures were identified from clustering local cell-type composition over sliding 100 μm circular windows, and projected back onto the cell-type map. The reconstruction shows the various cortical layers within the adult mouse VISp, with very clear separation of the Pia layer, and separation of layer 4 and layer 6 into 2 sub-layers. Inset zooms into the location of the rare Sst Chodl cell type found in layer 6-1; (B) Cell-type composition within each tissue domain.

Next, we continued with completely *de novo* cell-type identification. The resulting 30 cell-type signatures (**Fig. 2A, B, Supplementary Fig. 7-10**) were consistent with those identified in the segmentation-based clustering and scRNA-seq based cell-type signatures^6^ (**Supplementary Fig. 4C, D**), implicating the robustness of the *de novo* cell-type calling by SSAM. Each of the SSAM *de novo* cell-type signature clusters were assigned the label of the closest correlating segmentation-based cluster.

As with the guided mode analysis, we limit the comparison to the most comparable cell types, excluding cell types with low correlation in gene expression signatures (< 0.8) (**Supplementary Table 3, Supplementary Fig. 11**). The matching score result showed high average values (mean and median of 0.76 and 0.83, respectively) and 81% of cell types had a matching score of greater than 0.6. Comparing marker gene expression of cell types having lowest matching score (< 0.3) (**Supplementary Table 3**) confirmed that the SSAM guided cell-type map is in better agreement to their marker gene expression (**Supplementary Fig. 12-13**). Given the low correlation of C. Plexus cell type to the corresponding osmFISH cluster, which is one of the dominant cell types in the ventricle region, high-resolution investigation of Poly-A and DAPI signals confirm the existence of both cell types in the ventricle area (**Fig. 2D**). Since ependymal and choroid plexus cells were small and tightly packed and exhibit relatively lower DAPI and poly-A signal, we concluded that the performance of the watershed algorithm was insufficient to identify cells in the area. Furthermore, we statistically evaluated this for each cell type by comparing the gene expression in the unique parts of the segmentation and SSAM *de novo* cell-type map, to the overlapping parts (Methods). Gene expression of the unique part of SSAM *de novo* cell-type map showed higher correlation to the overlapping regions compared to the unique parts of the segmentation (**Supplementary Fig. 14**).

We then performed domain analysis on the SSAM *de novo* cell-type map. Identified domains correlated well with the known cerebral cortex layers, consistent with results reported in the previous study (**Fig. 3A**). Laminar distribution of cell types is established ^30^, and can be considered as a ground truth for validating the cell type map. Cell-type assignments of excitatory pyramidal cells in the cortical layers conformed closely to known localizations (**Supplementary Fig. 15**). The domains identified as: layer 2/3 primarily consists of Pyramidal L2-3/L5, L2-3, and L3-4 cell types; layer 4 consists of Pyramidal L4 and L3-4 cell types; layer 5 consists of Pyramidal L3-5 and L5 cell types; and layer 6 consists of Pyramidal L6 cell types.

In addition, cell-type composition of the domains revealed that *Mfge8* expressing astrocytes (Astrocyte Mfge8) contributed 7-14 % of each of the tissue layers (**Fig. 3B**), in contrast to the significantly fewer numbers of Astrocyte Mfge8 cells called in the previous study^6^. Comparison of high-resolution images of DAPI and poly-A signals with Mfge8 expression densities implicates that the poly-A signal was not strong enough to discriminate the presence of astrocyte Mfge8 cells from the background, while the DAPI images clearly supported the existence of *Mfge8* expressing astrocytes at positions identified by SSAM (**Fig. 2E**). The clear DAPI signal but low poly-A signal for these astrocytes Mfge8 suggested that they have a lower mRNA content compared to other cells. We compared the total counts of mRNA molecules of astrocytes and other cell types from mouse brain scRNA-seq data^31^ and found that astrocytes exhibited significantly less mRNA molecules than other cell classes (**Supplementary Fig. 4B**). Our observation reveals the inadequacy of the watershed segmentation algorithm applied to poly-A signal when not considering cells with a low total mRNA content. This implies that the original segmentation of these cell types could be less accurate than the SSAM *de novo* cell-type map, therefore also reducing the matching score for these cell types.

### SSAM confirms diversity of inhibitory and excitatory neuron cell types and their localization in the hypothalamic preoptic region (POA) in 3D

To demonstrate the performance of SSAM for three-dimensional *in situ* transcriptomics data, we applied SSAM to previously published MERFISH data, where 135 transcripts were localized in 3D space of the hypothalamic preoptic region (POA) of a mouse brain^5^ (**Fig. 4, Supplementary Fig. 16, 18**). We compare results obtained from SSAM against the results obtained from DAPI segmentation from the original study.

We applied both SSAM guided mode and *de novo* mode. For guided mode, the previously known cell-type signatures obtained by segmentation and scRNA-seq were used. For both guided and *de novo* modes, SSAM analysis was performed in 3D space, generating a 3D cell-type map (**Fig. 4B**). The resulting cell-type maps on the x-y plane at the center of slice on the z-axis (at 5μm) were visually similar to the previous study (**Supplementary Fig. 17G**). SSAM cell-type signatures showed high expression of their marker genes (**Supplementary Fig. 18-21**) and a high correlation to the cell-type signatures from both the segmentation-based clusters and scRNA-seq clusters (**Supplementary Fig. 17E, F**). Among them, 7 inhibitory and 4 excitatory neuronal cell types showed very high correlation (>0.8) to the segmentation-based neuronal signatures, and also showed distinctive tissue localization patterns (**Fig. 4D, E**), similar to those previously reported (**Supplementary Fig. 22**).

We then quantified the similarity of the SSAM cell-type maps with the cell segmentation by Moffitt et al. The SSAM guided mode cell-type map achieved high matching scores for comparable cell types (mean and median of 0.76 and 0.83 for segmentation-based, 0.88 and 0.94 for scRNA-seq-based signatures, respectively), with only 6 of 76 cell-types exhibiting a low matching score (< 0.3) for segmentation-based case (**Supplementary Table 4, 5, Supplementary Fig. 23, 24**). Comparing the SSAM *de novo* cell-type map also yielded high matching scores (mean and median of 0.83 and 0.93, respectively) (**Supplementary Table 6, Supplementary Fig. 25**), further validating the computational approach adopted by SSAM to identify *de novo* cell-type signatures and generating cell-type maps. One of the most notable differences in the SSAM cell-type map was that we found a higher density of astrocytes compared to Moffit et al. A comparative analysis revealed that some astrocyte signals identified by SSAM were not found in the segmentation by Moffit et al. Note that the existence of astrocytes is clearly shown by the corresponding marker gene expression (**Supplementary Fig. 26**).

The generated tissue domain map identifies several domains consisting of regions consisting primarily of inhibitory neurons, excitatory neurons and oligodendrocytes, as well as the ventricle structure (**Supplementary Fig. 27**).

Finally, we reconstructed a three-dimensional cell-type map (**Movie 1**). While the thickness of the tissue image is limited (10 μm), we demonstrate the shape and size difference of the whole cell-type map and the cell-type specific maps for inhibitory neurons, excitatory neurons and astrocytes (**Movies 2, 3, 4**).

Despite the difference of dimensionality between the osmFISH data (2D) and the MERFISH data (3D), SSAM was able to successfully process the data and produce meaningful results. More importantly, the analyzes in this section were performed with almost the same procedure and parameters applied to the osmFISH data. Therefore, we set these parameters as the default values to facilitate rapid and robust analysis of other multidimensional *in situ* transcriptomics dataset using SSAM.

### SSAM identifies rare cell types and novel cortical sub-layering in the adult mouse visual cortex (VISp)

To further demonstrate that SSAM can be used for rapid and robust analysis of *in situ* transcriptomics data, we applied SSAM to unpublished multiplexed smFISH data of the mouse primary visual cortex (VISp) generated as part of the SpaceTx consortium^32^ (**Fig. 5, 6, Supplementary Fig. 28, 29**). In total, the expression of 22 genes was quantified *in situ* (Methods).

Analysis of the tissue image was restricted to the manually defined VISp region (**Supplementary Fig. 28D**). SSAM was performed in both guided mode and *de novo* mode (**Supplementary Fig. 29A**). The guided mode of SSAM was performed using scRNA-seq data^30^. For the *de novo* run, the identified cell-type signature clusters were assigned the label of the cluster in the scRNA-seq data with the highest correlation (**Fig. 5A, B**). Then, the tissue domains were identified based on the *de novo* cell-type map (**Fig. 6**), with the result showing the laminar structure of the VISp region. We identified two distinct layer 4 (L4) neuronal clusters. Interestingly, both of them showed the highest correlation to the single L4 IT type identified via scRNA-seq, but their spatial locations show a clear difference (**Fig. 5C, Supplementary Fig. 29B**). We named the cluster localizing to the superficial region of layer L4 as ‘L4 IT Superficial’ (L4 IT 2). This finding adds context to the previously observed heterogeneity of the L4 IT cell type^30^, where we show that this heterogeneity determines superficial and deep localization in layer 4.

The cell-type map generated by SSAM guided mode were visually similar to that of *de novo* mode, except for the cell types found in the layer 2 (L2) (**Supplementary Fig. 29A**). We found that the majority of cell types found in L2 were assigned to the VLMC type in SSAM guided mode. We observed that this type was actually a neuronal type in L2. This cell type showed high expression of *Alcam*, a marker gene of the VLMC cell type, but low expression of other genes. Due to the limited number of genes profiled in the multiplexed smFISH experiment, lack of other neuronal marker genes led to incorrect high correlation of this type VLMC. However, SSAM properly assigned the centroid to be L2 neurons in *de novo* mode.

SSAM was also able to identify a rare cell type, Sst Chodl, which is known to be related to long-range projection and sleep-active neurons^33–35^. In addition, we mapped the Sst Chodl cell-type signal to between layer L5 and L6 (**Supplementary Fig. 29C**), consistent with previously reported localization to L5 and L6^33^. This finding was validated against its marker gene expression (**Supplementary Fig. 30-32**), and ultimately demonstrates SSAMs ability to identify cell-type signatures of lowly abundant and rare cell-types.

## Discussion

We describe a segmentation-free computational framework for processing *in situ* transcriptomics data and demonstrate its performance on three different adult mouse brain datasets: the somatosensory cortex (SSp) profiled by osmFISH, the hypothalamic preoptic region (POA) by MERFISH, and the visual sensory cortex (VISp) by multiplexed smFISH. We find that the cell-type signatures and maps generated by SSAM for both osmFISH and MERFISH datasets were similar to the previously reported ones, validating the underlying methodology of SSAM. Based on this, we successfully determined cell types and constructed cell-type and tissue domain maps in the multiplexed smFISH mouse VISp dataset.

In the osmFISH dataset our method outperforms the original segmentation-based cell-type map reconstruction due to limitations in the segmentation process. In the MERFISH dataset we show that SSAM is able to identify diverse populations of cell types and that SSAM is scalable to 3D image data. For the VISp multiplexed smFISH dataset, SSAM identified a rare cell type and elucidated a suspected spatial heterogeneity of cell types in the cortex without segmenting a single cell. Overall, the results show that SSAM is not only a robust tool to validate segmentation-based methods, but also a reasonable alternative when segmentation is difficult or DAPI or Poly-A images are lacking.

However, for some questions it is important to distinguish between cells to e.g. delineate growth arising from increasing cell size vs cell proliferation or to investigate multinucleation in cardiomyocytes or cytotrophoblast cells. In cases such as these, we recommend the use of SSAM as a complementary method to segmentation-based analysis in two ways. First, the output of SSAM can be compared to validate that the segmentation process did not introduce artifacts. Secondly, to use the SSAM output as an input for the segmentation process to refine the segmentation procedure for different domains or cell-type signals.

In terms of methodological parsimony, SSAM minimizes the number of assumptions, avoids iterative optimization and thus offers maximal transparency, interpretability and reproducibility. The lightweight nature of the algorithm typically brings a considerable runtime advantage over other available packages. SSAM is written as a Python library, with some core analysis functions wrapped up with external C functions to speed up the computation. The package is available as an easily installable Python package, and can easily be extended with existing *in situ* transcriptomics pipelines, e.g. starfish (https://github.com/spacetx/starfish) or Giotto^36^. SSAM is accompanied with a notebook outlining all the steps presented in this paper. Taken together, we present a novel, flexible and robust method for fully automated cell-type and tissue domain analysis that is readily applicable to numerous *in situ* transcriptomics methods.

## Materials and Methods

### Using Kernel Density Estimation to generate the gene expression vector field

We used the n-dimensional KDE algorithm to estimate the density of mRNAs in 2D and 3D. To compute Gaussian KDE, we used our own implementation of the KDE algorithm for rapid computation. Spatial distribution of the probability of mRNA presence 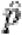 is estimated using the kernel density estimation;

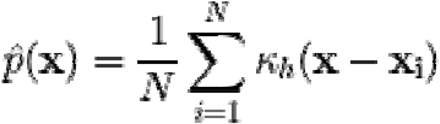

where:

- *K*_*h*_: a kernel function with a bandwidth *h*
- *N*: the number of data points
- X_1_: location vector of the data point i (i.e. location of i-th mRNA)

Here we use the Gaussian kernel:

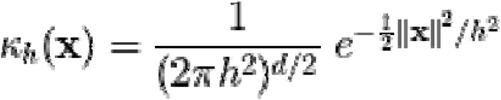

where:

- *h*: bandwidth of the Gaussian kernel
- *d*: dimension of the space where the data points reside (2 for 2D, or 3 for 3D mRNA locations)
- ‖X‖: Euclidean norm (i.e. L2 norm) of vector x

Note that the integration of 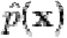 all over the space is 1. Therefore the gene expression density is calculated by multiplying the number of mRNAs per gene to .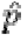

### Calculation of spatial gene expression

The continuous estimation of gene expression density is discretized over pixels of the tissue image, which in our examples is set to a size of 1□m. The expectation value of the estimated density in a unit pixel is approximated by multiplying the area of the unit pixel to the estimated gene expression density at the location of the pixel. Finally, we stack the estimated gene expression densities of genes to define the gene expression vector field over the image.

### Selection of local maxima

Local maxima were selected based on the L1-norm of the vectors in the vector field, which is the total size of each vector in the image. For the selection algorithm, we used scikit-image Python package to select local maxima. Briefly, 1) maximum filter is applied to dilate the original image, 2) the locations where the maximum filtered image equal to the original image are selected. The maximum filter with size 3 was used throughout the examples presented in this paper.

### Downsampling of the vector field

For a scalable cell-type identification analysis, the vector field is downsampled to a smaller set of vectors based on local maxima selection strategy (Supplementary discussion). SSAM applies two thresholds for local maxima selection: 1) a minimum expression threshold for a single gene defined as the height of a single Gaussian kernel to avoid regions with signal from only the Gaussian tail (see Discussion section for details), which also corresponds to the position of the observable drop in the histograms of gene expression (**Supplementary Fig. 3A, 17A, 28A**); 2) a minimum total gene expression (i.e. L1-norm) threshold (**Supplementary Fig. 3B, 17B, 28B**). Furthermore, we implemented an optional “input mask” feature to limit sampling of vectors to regions of the image containing informative data, e.g. a mask outlining the informative tissue area.

### Comparison of local maxima and random sampling strategies

The two local maxima sampling methods, 1) local maxima sampling and 2) random downsampling, were compared to justify our preference of local maxima sampling method for the downstream analysis. The osmFISH data was used for the comparison. Firstly 11,469 local maxima vectors were found in the vector field using a window size of 3, a minimal gene expression and L1 norm thresholding. For comparison, the same number of vectors were randomly sampled from the vector field, using the same thresholds used for local maxima selection. At the locations of the vectors, both the local maxima and the random sampled locations, the classified cell types on the cell-type map guided by segmentation-based signatures are called. For each case, the Pearson’s correlation coefficients between the vectors and the signature of the cell types are calculated and plotted as a distribution (**Supplementary Fig. 39**).

### Variance stabilization of local maxima vectors and the vector field

Since the gene expression profiles of local maxima vectors are representative of the transcriptomes of cells, we considered them to be analogous to the gene expression count matrix obtained from single cell RNA sequencing (scRNA-seq) using unique molecular identifiers (UMI). Therefore, we normalized the local maxima vectors of the vector field (which would be representative of single cells) using *sctransform*^*37*^, a normalization and regularization algorithm for UMI count data. After that, each vector of the vector field is normalized using *sctransform*, with the same parameters previously used to normalize the local maxima.

### Clustering of representative gene expression vectors

The SSAM framework supports clustering via DBSCAN^24^, HDBSCAN^25^, OPTICS^26^ and an implementation of the Louvain algorithm equivalent to that in the R package, Seurat^27^. DBSCAN, HDBSCAN and OPTICS are implemented via the scikit-learn Python library. The Louvain clustering algorithm is based on the R package Seurat^27^ reimplemented in Python. In short, an SNN network with correlation metric is built using a python package NetworkX^38^. The weight of the network is calculated by a Jaccard similarity coefficient. A weight smaller than 1/15 was set to zero. Clustering was done by detecting communities in the network using a Louvain community detection algorithm implemented in Python (python-louvain, https://python-louvain.readthedocs.io/). It is known that the Louvain algorithm is not sensitive in detecting small clusters^39^, optionally DBSCAN algorithm can be applied to subcluster each Louvain cluster. This sub-clustering strategy is conceptually similar to the “Polished Louvain” algorithm in Zeisel et al^31^.

### Diagnostic plots

After unsupervised clustering of gene expression vectors, some clusters may need to be manually merged or discarded. SSAM supports merging of clusters based on correlation of gene expression profile, however in many cases manual inspection is needed to rule out any non-trivial issues. To guide this process, SSAM generates a cluster-wise ‘diagnostic plot’, which consists of four panels: 1) location of the clustered vectors on the tissue image, 2) the pixels classified to belong the cluster signature (the cluster centroid), 3) the mean expression profile of the clustered vectors, and 4) the t-SNE or UMAP embedding.

In the three datasets analyzed the clusters to be merged or removed often showed a discordance between the location of sampled vectors used to determine the cluster (panel 1) and the pixels classified to belong to that cluster (panel 2). In case of overclustering, i.e. when a cell-type signature is split over 2 clusters, the map typically does not classify the full shape of the cells but instead only fragments (panel 2), and having almost the same marker gene expression of another cluster (panel 3). Such clusters can be merged. For dubious clusters that should be removed, we observed that vectors usually originate from outside the tissue region or from image artifacts (panel 1), or that the gene expression does not show any clear expression of marker genes or similarity to expected gene expression profiles (panel 3). The remaining clusters are then annotated by comparing cluster marker genes to known cell-type markers. Note that in many cases, the identity of clusters can be easily assigned by comparing the centroids of the clusters to the known cell-type signatures, e.g., from single cell RNA sequencing. To support rapid annotation of cell types to clusters, SSAM additionally shows the highest correlating known cell-type signature should this data be available in panel 3. The diagnostic plots for osmFISH, MERFISH, and multiplexed smFISH data are available online in the Jupyter notebook uploaded to zenodo (http://doi.org/10.5281/zenodo.3478502).

### Statistical evaluation of cell-type mapping

The accuracy of the SSAM cell-type map was validated by comparing the published osmFISH segmentation and the SSAM *de novo* cell-type map by two different methods.

Firstly, to quantitatively compare concordance of cell-type we implemented a matching score. The matching score for any given cell type is defined as the number of segmented cells with at least 10% of matched with the SSAM guided or *de novo* mode cell type map of the corresponding cell type of the segment, divided by the total number of segments of the cell type which represents the ratio of segments identified by SSAM. The threshold of 10% was empirically selected to account for differences in cell location in the tissue, especially for very small cells where subtle changes in cell-type labeling can drastically reduce the overlap within the segmented area.

Secondly, for evaluation of discrepancies in cell-type locations compared to the original studies, we compare the unique part of each segmentation and SSAM *de novo* cell-type map to the parts that are overlapping in both maps in the osmFISH dataset. The gene expression vectors originating from overlapping parts of the same cell types (**Supplementary Table 3**), were regarded as the ground truth set. Then, two sets of unique vectors were defined: 1) the segmentation-only set, the vectors from the regions occupied by segments excluding the overlap, and 2) the SSAM-only set, the vectors from SSAM cell-type map only regions. The distribution of the gene expression vectors in the overlapping set was then compared to the two unique parts (**Supplementary Fig. 14A**). To compare the accuracy of cell-type mapping of the two unique parts, Pearson’s correlation coefficient is calculated between the mean expression of the ground truth set and the vectors in each set (**Supplementary Fig. 14B**).

### Quantification of doublets

The doublet rates were evaluated by two Python packages, DoubletDetection^40^ and Scrublet^41^ (**Supplementary Table 8**). As the two algorithms require raw counts as input, the unnormalized raw vectors at local maxima used for clustering analysis were used as input of the two algorithms, as an analogy of the raw counts. For DoubletDetection, the doublet rate was calculated by dividing the number of doublets reported by the number of total local maxima. The doublet rate quantification by both methods was consistent, and negligible in the osmFISH and multiplexed smFISH datasets (average doublet rate of <0.5% for both), and marginal for MERFISH (average doublet rate of 3%).

### SSAM analysis of osmFISH data

KDE was performed with a bandwidth of 2.5 μm. The individual gene expression threshold and total gene expression threshold for selection of local maxima were 0.027 (the height of a single Gaussian) and 0.04, respectively (**Supplementary Fig. 3A, 3B**). Since the selected local maxima includes many locations outside of the tissue area, we further filtered local maxima based on their local density approximated using the k-nearest neighbor algorithm. More specifically, local maxima with a density lower than 0.002 over the closest 100 local maxima, corresponding to fewer than 100 local maxima in a 126.2 μm radius, were filtered out (**Supplementary Fig. 3C**). The selected local maxima vectors were passed to *sctransform* to determine normalization parameters, after which the whole vector field was normalized.

In SSAM guided mode, the mRNA count matrix of both the previously segmented cells and the scRNA-seq data were normalized by *sctransform*. The centroid of each of the annotated clusters was used to classify cell types in the vector field, generating a cell-type map guided by prior knowledge.

In SSAM *de novo* mode, the selected local maxima vectors were clustered using the Louvain algorithm with a resolution of 0.15, resulting in 66 clusters (**Supplementary Fig. 4A**). Distinct clusters representing the same cell types were identified and then manually merged, and spurious clusters were removed, resulting in a total of 30 clusters (**Fig. 2A, 2B**). For each cluster, the vectors with insufficient correlation to its cluster medoid were excluded from the centroid calculation (**Supplementary Fig. 1B**). The cluster centroids were compared to that of the segmentation-based (**Supplementary Fig. 4B**) and scRNA-seq cell-type signatures (**Supplementary Fig. 4C**) using Pearson’s correlation coefficient. The *de novo* clusters were named after the highest correlating segmentation-based cluster. Note that clusters closest mapped to Inhibitory IC and Inhibitory CP cell types do not only appear in the internal capsule and caudoputamen, but also in the cortex. Therefore, we renamed these clusters to Inhibitory Kcnip2 (since Kcnip2 was the third most expressed gene for this cluster) and Inhibitory Rest, respectively. After classification of the local maxima, we quantified the doublet rates (Methods, **Supplementary Table 8**).

Tissue domain analysis was performed using a sliding circular window with radius 100 μm with a step of 10 μm. The cell-type proportions from each window were clustered using agglomerative hierarchical clustering with 15 clusters as an initial estimate, subsequently merging the clusters with correlation coefficients higher than 0.8. Spatially connected clusters with a correlation coefficient higher than 0.6 were merged. The resulting domain map was resized to match the size of the cell-type map, after which the cells in different domains were colored.

### Quantification of mRNA abundance in astrocytes and other brain cell types for osmFISH data interpretation

The “L5_All.loom” loom object containing scRNA-seq expression data of half a million cells from the mouse nervous system^31^ was downloaded (http://mousebrain.org/downloads.html). The total number of mRNA molecules per cell were extracted and aggregated by their level 2 class labels (astrocytes, immune, vascular, ependymal, neuronal, peripheral glia and oligodendrocyte cells) using Python. The counts were log normalized and subsequently followed a normal distribution (tested using the Shapiro-Wilk test for normality, all *p-values* < 1 x 10e-4 for each class), therefore a Student’s t-test was applicable. For each of the two classes of interest (‘Astrocytes’, ‘Immune’), we performed independent log-space t-tests for unequal sample sizes and unequal variance against each of the other classes. Both astrocyte and immune cell classes have significantly lower mRNA molecule counts compared to other cell types (all *p-values* < 1 x 10e-12). While the distribution of mRNA counts in log space followed a normal distribution, the use of a Student’s t-test for large numbers may be not appropriate. Hence, we also describe the difference in their distributions. For both astrocyte and immune cell classes, more than half of the cells of each class exhibited a lower UMI count than the lowest quartile of any other cell class.

### SSAM analysis of MERFISH data

KDE was performed with bandwidth 2.5 μm. Local maxima were filtered using a gene expression threshold of 0.0055, and then filtered with total gene expression threshold of 0.0035 (**Supplementary Fig. 17A, B**). The selected local maxima vectors were passed to *sctransform* to determine normalization parameters, after which the whole vector field was normalized.

In SSAM guided mode, the mRNA count matrix of both the previously segmented cells and the scRNA-seq data were normalized by *sctransform*. The centroid of each of the annotated clusters was used to classify cell types in the vector field, generating a cell-type map guided by prior knowledge.

For SSAM *de novo* mode, the selected vectors were clustered using the Louvain algorithm with a resolution of 0.15, resulting in 68 clusters (**Supplementary Fig. 17C**). By manual inspection of gene expression and localization, overclustering was merged, and spurious clusters were removed, resulting in a total of 50 clusters (**Fig. 2A, 2B**). For each cluster, the vectors that did not have high correlation to its cluster medoid were excluded from the centroid calculation (**Supplementary Fig. 1B**). The centroids of the clusters are compared with that of the segmentation-based clustering result and scRNA-seq result using Pearson’s correlation coefficient (**Supplementary Fig. 17E, F**). The SSAM *de novo* clusters correlating best to inhibitory and excitatory neurons were named based on the most highly expressed gene of each cluster, and the non-neuronal clusters were named based on the previous study^5^. After classification of the local maxima, we quantified the doublet rates (Methods, **Supplementary Table 8**). We noticed a number of small blobs on the cell type map, which are resultant from cells on a different plane in the 3D image (**Movie 2**). After classification of the local maxima, we quantified the doublet rates (Methods, **Supplementary Table 8**).

Tissue domain analysis based on the cell-type map was performed using a sliding spherical window with radius 100 μm with a step of 10 μm. The cell-type proportions from each window were clustered using agglomerative hierarchical clustering with 20 clusters as an initial estimate, subsequently merging the clusters with correlation coefficient higher than 0.8. The resulting domain map was resized to match the size of the cell-type map, after which the cells in different domains were colored.

### Comparison of localization of inhibitory and excitatory neurons

For a number of inhibitory and excitatory neuronal subtypes identified in the posterior POA tissue image using SSAM *de novo* mode, we identified the best matching cell types based on Pearson correlation of their gene expression signatures (**Supplementary Fig. 17F)**. We matched the following cell types: SSAM cluster 39 (C39) called Inhibitory Coch to Moffitt cluster I-12, C16 Inhibitory Arhgap36 to I-13, C45 Inhibitory Isr4 to I-15, C34 Inhibitory Calcr to I-14, C14 Inhibitory Gda to I-23, C19 Excitatory Cbln1-Cbln2 to E-19, C42 Excitatory Omp to E-16, C25 Excitatory Necab1-Gda to E-9, C8 Excitatory Necab1 to E-14, and C36 Excitatory Col25a1 to E-24. For these cell types we checked the tissue localizations reported in the previous studies figures 5a, 5c, 5e, 6b, 6d, and S17^5^. Side-by-side comparison of the localization of these neuronal cell types revealed very similar patterns of localization computed by SSAM and the original publication (**Supplementary Fig. 22**).

### 3D modelling of MERFISH cell-type maps

Firstly, the connected components in 3D were determined using the python package connected-components-3d (https://github.com/seung-lab/connected-components-3d). Components comprising fewer than 100 voxels were removed. After this, the voxels filling connected components were removed, and only the contours were used for the vertex of the 3D models. For each vertex, the vertex normal was calculated by simple physics simulation, assuming that the direction of a vertex normal vector is the same as the force vector when there are pulling forces between all of the contour voxels. The surface of the objects was reconstructed using screened Poisson reconstruction algorithm^42,43^ using default parameters. The number of vertices was reduced to 5% of the total number of vertices using the ‘vtkQuadricDecimation’ function^44,45^ of VTK library^46^. Finally, the objects were merged into a single file. Each scene of the rotating movie was created using Meshlab^47^.

### VISP multiplexed smFISH data generation

Multiplexed smFISH data of the mouse primary visual cortex (VISp) was generated as part of the SpaceTx consortium. Tissue processing was carried out as previously described^48^, with some modifications.

Silanization of coverslips (#1.5, Thorlabs CG15KH) was performed by plasma cleaning for 30 min in a Plasma-Prep III (SPI 11050-AB), followed by vapor deposition of 3-aminopropyltriethoxysilane (APES, Sigma A3648) in a vacuum for 10 minutes. Coverslips were then washed in 100% methanol for 2 x 5 minutes, allowed to dry, and stored in a dust-free environment until use.

Fresh-frozen mouse brain tissue was sectioned at 10 μm onto silanized coverslips, let dry for 20 min at -20°C, then fixed for 15 min at 4 °C in 4% PFA in PBS. Sections were washed 3 × 10 min in PBS, then permeabilized and dehydrated with chilled 100% methanol at -20°C for 10 min and allowed to dry. Sections were stored at -80 °C until use. Frozen sections were rehydrated in 2X SSC (Sigma 20XSSC, 15557036) for 5 min, then treated 10 min with 8% SDS (Sigma 724255) in PBS at room temperature. Sections were washed 5 times in 2X SSC. Sections were then incubated in hybridization buffer (10% Formamide (v/v, Sigma 4650), 10% dextran sulfate (w/v, Sigma D8906), 200 µg/mL BSA (ThermoFisher AM2616), 2 mM ribonucleoside vanadyl complex (New England Biolabs S1402S), 1 mg/ml tRNA (Sigma 10109541001) in 2X SSC) for 5 min at 37°C. Probes were diluted in hybridization buffer at a concentration of 250 nM and hybridized at 37°C for 2 h. Following hybridization, sections were washed 2 × 10 min at 37°C in wash buffer (2X SSC, 20% Formamide), and 1 × 10 min in wash buffer with 5 μg/ml DAPI (Sigma 32670), then washed 3 times with 2X SSC. Sections were then imaged in Imaging buffer (20 mM Tris-HCl pH 8, 50 mM NaCl, 0.8% glucose (Sigma G8270), 30 U/ml pyranose oxidase (Sigma P4234), 50 µg/ml catalase (Abcam ab219092). Following imaging, sections were incubated 3 × 10 min in stripping buffer (65% formamide, 2X SSC) at 30°C to remove hybridization probes from the first round. Sections were then washed in 2X SSC for 3 × 5 min at room temperature before repeating the hybridization procedure.

The multiplexed smFISH image data was collected and processed using methods previously described^48^, except that images from different rounds of hybridization were registered in (x, y) based on the DAPI signal. The raw images are available on request.

### SSAM analysis of VISp multiplexed smFISH data

KDE was performed with bandwidth 2.5 μm. Local maxima were filtered using a gene expression threshold of 0.027, and then filtered with total gene expression threshold of 0.2 (**Supplementary Fig. 28A, B**). The selected local maxima vectors were passed to *sctransform* to determine normalization parameters, after which the whole vector field was normalized. To identify rare cell types expected to exist in this tissue, the initial clustering result by Louvain algorithm was sub-clustered by DBSCAN (Method). Initially 49 clusters were obtained with a resolution parameter of 0.15. By manual inspection, several over-clustered cell types, including nine L2/3 IT 1, two L2/3 IT 2, six L4 IT 2, six L6 CT, and two L6 IT 2 clusters were merged, and one spurious cluster was removed, resulting in 28 clusters. The centroids of the clusters are compared with that of scRNA-seq result using Pearson’s correlation coefficient (**Supplementary Fig. 28E**). The clusters were named after the highest correlating scRNA-seq cluster, except the newly found ‘L4 IT Superficial’ (L4 IT 2) cluster. After classification of the local maxima, we quantified the doublet rates (Methods, **Supplementary Table 8**).

Tissue domains were defined using a sliding circular window with radius 100 μm with step of 10 μm over the cell-type map image. Cell type compositions of the windows were clustered using agglomerative clustering, initially with 20 clusters. Clusters with Pearson’s correlation higher than 0.7 were merged to result in nine clusters. Further, two clusters were merged since they were different parts of the Pia layer, resulting in a final set of seven clusters representing tissue domains (**Fig. 6**).

### Plotting

The python packages Matplotlib 3.1.0^49^ and Seaborn 0.9.0^50^ were used to draw 2D images, plots, and heatmaps. We include helper functions in SSAM to easily generate plots.

### Movies

Movies were generated by using Virtualdub (1.10.4-AMD64, http://www.virtualdub.org/). The H.264 codec was used to compress videos.

### Software

Python version 3.7.0 was used throughout. The following python packages were used: *numpy, scipy, pandas, matplotlib, seaborn, scikit-learn, umap-learn, python-louvain, sparse, scikit-image*. R package *sctransform* was used for normalization and variance stabilization of the data.

### Data availability

The source code of SSAM is available online at https://github.com/eilslabs/ssam. A Jupyter notebook (https://github.com/eilslabs/ssam_example) outlines the commands used to download and pre-process the data, and to reproduce the results and figures of this study. The Jupyter notebooks also contain the extensive diagnostic plots used for parameter selection, and choice of removal or merging of clusters. All large files are available online from http://doi.org/10.5281/zenodo.3478502.

The osmFISH data (Codeluppi et al., 2018) used within the study is available from http://linnarssonlab.org/osmFISH/availability/. The single cell RNA sequencing data of the mouse somatosensory cortex^28,29^ are available from http://loom.linnarssonlab.org/. The single cell RNA sequencing data^31^ used to compare total mRNA molecules between cell types are available from http://mousebrain.org/. The high resolution poly-A and DAPI images of osmFISH data (Codeluppi et al., 2018) were kindly provided by Sten Linnarsson. The MERFISH data (Moffitt et al., 2018) is available from https://datadryad.org/handle/10255/dryad.192644. Mouse VISp multiplexed smFISH data are available from http://doi.org/10.5281/zenodo.3478502.

## Acknowledgements

We thank Sten Linnarsson and Jeffrey Moffitt for providing support and access to the osmFISH and MERFISH datasets, respectively. We also thank Yue Zhuo, Ed Lein, Jeremy Miller, Ambrose Carr, Nagarajan Paramasivam, Stephen Krämer, Zuguang Gu, Daniel Hübschmann, Luca Tosti, and Christian Conrad for helpful discussions and comments on data analysis. The authors also thank Bianca Hennig for designing Figure 1, and assistance in improving figures. The preliminary analysis of multiplexed smFISH data occurred during the SpaceTx SpaceJam Hackathon at the Allen Institute for Brain Science, which was organized by Ed Lein, and generously supported by the Chan Zuckerberg Initiative. This publication is part of the Human Cell Atlas - www.humancellatlas.org/publications. This research has received funding from the European Union’s Horizon 2020 research and innovation program under grant agreement No 824110 – EASI-Genomics, and was supported by the European Commission (ESPACE, 874710, Horizon 2020).

## Author contributions

JP, WC designed the concept and idea of SSAM.

JP, WC, RE, NI conceived the study.

BT, EG, TN.N, BL acquired and interpreted the multiplexed smFISH data.

JP, WC, ST, TN.N, NI performed data analysis.

LE.B, MS, BL, BT, TG, OS provided critical comments and discussions.

RE, NI supervised the study.

All authors commented on and critically revised the manuscript.

## Competing interests

The authors declare no competing interests.

## Notes

http://doi.org/10.5281/zenodo.3478502

### Competing Interest Statement

The authors have declared no competing interest.

### Summary of Updates

Substantial updates of main and supplemental text and figures.

