## Supplementary information for "Cell segmentation-free inference of cell types from *in situ* transcriptomics data"

Jeongbin Park, et. al

### Content

- Supplementary discussion
- Supplementary tables 1 – 8
- Supplementary figures 1 – 40
- Supplementary video legends
- Supplementary references

### Supplementary discussion

#### Modelling gene expression using Kernel Density Estimation (KDE)

Established principles of mRNA molecules being subject to Brownian motion or restricted Brownian motion^1–3^ constrained within the cells (plasma and internal) membranes, justifies the use of kernel density estimation (KDE) to reconstruct gene expression patterns. However, active transport of mRNA breaks the assumption of molecules undergoing Brownian motion, and this would increase the complexity of the signals being detected by SSAM. We postulate that when profiled mRNA are subject to mechanisms such as active transport or subcellular compartmentalization, SSAM may be able to identify such compartments within a cell. We observed such a phenomenon for astrocyte cell-types in the osmFISH dataset (see *Subcellular gene expression signals*). We adopted the use of a Gaussian kernel, because of the simplicity of calculations, however other kernels may also be appropriate, e.g. the Epanechnikov kernel^4^.

#### Choice of kernel bandwidth and pixel resolution

The default bandwidth for KDE is 2.5 μm, which makes its FWTM to be 10.7 μm being close to the average size of cells in the mouse brain SSp tissue image (11.6 μm). However, the choice of kernel bandwidth based on the size of cells defines maximum limit of the bandwidth, and the minimum should rather model the distance between the profiled mRNA. In other words, the bandwidth needs only to be sufficient to smooth the mRNA distribution over most of the cell body to identify a local maxima vector which represents that cell’s gene expression profile. Such variation in mRNA density can arise from the tissue or the profiling technique. So, when the density of mRNA molecules is high, a smaller bandwidth can be used to reconstruct more detailed picture of cell type localizations, but when the density of mRNA is low a larger bandwidth should be employed. We investigated the effect of bandwidth and the pixel resolution (lattice spacing) to which the gene expression is resolved (**Supplementary Fig. 16-19**). This showed that the cell-type signatures and cell type map are not highly dependent on pixel resolution/lattice spacing, that small deviations in the bandwidth have negligible effect, and large increases in the bandwidth can result in an overly smoothed cell type map while still capturing most cell-type signatures (**Supplementary Fig. 20**).

#### Downsampling of the vector field

SSAM downsamples gene expression vectors before performing cluster analysis. This step significantly reduces computation time compared to processing the entire tissue image, especially for graph-based clustering algorithms. We show that random sampling from the tissue image is effective (**Supplementary Fig. 21**), however we achieve better representation of cell-type gene expression profiles by selecting local maxima of total gene expression (**Supplementary Fig. 22**) and as such made this this default downsampling method. The reasoning of the local maxima sampling strategy follows two presumptions: 1) Pixels with a higher density of mRNA count have a larger probability of being intracellular, and that 2) Gaussian smoothing locally propagates the signal sufficiently to average the signal over intracellular regions. From those presumptions, we reason that vectors with a peak signal intensity are probably situated centrally within a cell and hence have a high signal-to-noise ratio and capture a representative signature of its cell type by integrating the surrounding mRNA molecules.

#### Clustering and cell-type gene expression profile analysis

An important step in determining the cell-type specific gene expression profiles is the clustering step. We initially implemented the Louvain algorithm for cluster analysis, which is used routinely for cluster analysis of scRNA-seq data in packages such as Seurat^5^, however we found that this was not sensitive enough to detect all cell types, in particular the Sst Chodl type in the mouse VISp dataset. In order to address this, we employed an approach similar to a previous study that applied a second round of clustering on top of the Louvain results^6^ to split potentially merged clusters. We successfully applied HDBSCAN clustering on the Louvain clustering results and were then able to identify a distinct cluster for Sst Chodl. Despite this success, it remains unclear if the same protocol would work equally well for data from different tissues and techniques, and therefore encourage the use of also other clustering methods.

#### Cell type mapping

We have employed the use of Pearson’s correlation to classify pixels in the tissue image, generating the cell-type maps (**Supplementary Fig. 2B**). Correlation-based cell-type mapping has been proven to work effectively^7,8^, and was also shown to be among top performers for automatic cell-type annotation^9^. Although our cell-type mapping worked well for the examples described in the paper, the SSAM framework allows for use of other classification strategies. For example, we have implemented an adversarial autoencoder to classify each pixel into each cell type. This preliminary work is part of the developmental branch of SSAM on *Github*.

#### Mixing of gene expression signals

In our clustering analysis of all three datasets, we observed that some clusters showed moderate mixed expression of signature genes from different cell types. This can be problematic for developmental tissues where cells can be tightly packed and transitioning between progenitor and differentiated states. Technical causes of signal mixing include: (1) different cells can overlap at different z location if the thickness of the section is comparable to the cell size; (2) clustering of the vectors might not be perfect and can include vectors in nearby clusters; (3) the gene expression estimated by the KDE algorithm is smoothed and the gene expression of one cell type can contaminate cells with different cell types located nearby. We found that (1) and (2) are of major importance for this phenomenon, but (3) also becomes noticeable for closely packed small cells. For example, we found that a relatively higher expression of the *Foxj1* gene (a signature gene of ependymal cells) is detected in the choroid plexus signature, compared to that of segmentation-based osmFISH centroid. Such signal ‘contamination’, caused by the KDE spreading signal into adjacent cells, can be controlled by the bandwidth value of KDE - the smaller the bandwidth, the lower the contamination; however, use of very low bandwidths break one of the primary assumptions of SSAM in that the smoothed KDE signal should represent cells, and not subcellular features. Still, our work showed that this phenomenon is not so critical as to hinder detection of cell types, even with extreme cases like bandwidth of 5 μm or 10 μm (**Supplementary Fig. 20**). However, we recommend trying several bandwidths to investigate this mixing effect in different tissues. Systematic evaluation of potential doublets arising from signal mixing revealed that the real effect of this phenomena is marginal (**Supplementary Table 8**).

#### Subcellular gene expression signals

SSAM identified a distinct cell-type signature of Aldoc-expressing astrocytes that also had low expression of *Gfap* and *Mfge8*. When looking closely at the localization of *Aldoc* signal they exhibited subcellular compartmentalization patterns, consistent with astrocytes that express high levels of *Mfge8*. We believe that this could due to localization of these mRNAs to internal subcellular localization compartments in Astrocytes. Such an intracellular spatial organization of the transcriptome is often an important form of post-transcriptional regulation^10^ and imaging-based methods can reveal this organization^11^. While this work does not investigate this in detail, this indicates that SSAM has potential to be used to identify and investigate organization of mRNAs.

#### Applicability of SSAM to other spatial gene expression profiling techniques

Currently, the field of *in situ* transcriptomics is advancing rapidly and more than 10,000 genes can be simultaneously profiled using FISH-based methods^12,13^. The high number of genes detected in large volumes opens up the potential for *in situ* transcriptomics methods to at least partially replace single cell RNA sequencing, placing SSAM as the first generic and segmentation-free pipeline to rapidly and precisely reconstruct tissue structure independent of the underlying imaging technique. While the profiled genes for the datasets presented in this study were carefully curated to represent cell-type diversity, when a larger number of genes are used we would suggest applying dimensionality reductions techniques before cluster analysis. Moreover, since the only required input data for SSAM is mRNA locations, it is highly adaptable to spatially resolved transcriptomics technologies beyond FISH methods, e.g. *in situ* or intact tissue sequencing^14–16^, and composite *in situ* imaging^17^.

In addition to these single molecule profiling techniques, there are those that are resolved to single cells or larger regions^18^, and Spatial Transcriptomics^19,20^. These come with the advantages and disadvantages that mRNA is aggregated to single cells/spots, e.g. making allocation of mRNA to cells easier, but also making subcellular localization analysis impossible. These techniques have a resolution limited to the spotting profile of the technologies, compared to the native organization of mRNA within tissues. For example, the size of the spots for Visium (55um) are very large, making it less appropriate for those interested in single cell resolution, and inappropriate for tissues with very small structures, e.g. placental villi which can be 60 um in diameter, or for those interested in localization of cells that are distributed over the tissue (e.g. delta cells in the pancreas).

#### SSAM as a segmentation-free analysis framework

We define SSAM as a framework, since it consists of various different functions required for segmentation-free spatial cell-type detection. For example, one can choose 1) local maxima selection, or random sampling for downsampling, 2) *sctransform**^21^*, L1 or L2 normalization combined by log-transformation for the normalization of vectors, or 3) Louvain (Seurat) clustering^5^, DBSCAN^22^, HDBSCAN^23^, or OPTICS^24^ as the clustering strategy. SSAM provides such functions as different modules already included with the package as ‘batteries-included’, therefore various different analysis scenarios are possible and different pipelines can be easily made with SSAM.

Thanks to the modular structure of SSAM framework, it is also easy to add new functionality to SSAM to reflect further user requests. For example, since we have made our package available on *Github*, we have received frequent requests to implement segmentation based on cell-type map as prior, therefore we have worked on a preliminary watershed segmentation feature. Also there have been some concerns with scalability of SSAM, therefore we added support of *dask*, which enables lazy-computation of arrays without loading the whole data in memory, so that SSAM can be applied to extremely large datasets without memory concerns. Both experimental functionalities are available in the developmental branch of SSAM on *Github*.

#### Marker gene selection

The selection of genes to profile is important for successful cell-type detection - i.e. the genes must be carefully selected to be suitable cell-type markers prior to using the SSAM algorithm. Although this has to be carefully considered on the experimental side, we benchmarked the cell-type detection rate as a function of the number of genes imaged, using several simulated settings by removing proportion of genes from MERFISH dataset (Supplementary Fig. 40). Surprisingly, we found that even when removing 80% of the genes, we were still able to recapitulate more than 65% of the cell-type signatures obtained by *de novo* analysis using all genes. The high variance of each category shows that the number of cell types detected also highly depends on which genes were selected for analysis, which means that not only the number of genes, but also the selection of genes is important as well.

### Supplementary Table 1. The matching score between the osmFISH PolyA segmented cells vs. SSAM cell-type map guided by the segmentation-based cell-type signatures of osmFISH data.

| **Cell type** | **Matched Segments** | **Total Segments** | **Matching score** |
| --- | --- | --- | --- |
| Astrocyte Gfap | 80 | 81 | 0.99 |
| Astrocyte Mfge8 | 109 | 119 | 0.92 |
| C. Plexus | 9 | 50 | 0.18 |
| Endothelial | 155 | 235 | 0.66 |
| Endothelial 1 | 6 | 94 | 0.06 |
| Ependymal | 98 | 107 | 0.92 |
| Hippocampus | 97 | 133 | 0.73 |
| Inhibitory CP | 69 | 161 | 0.43 |
| Inhibitory Cnr1 | 29 | 40 | 0.72 |
| Inhibitory Crhbp | 102 | 120 | 0.85 |
| Inhibitory IC | 67 | 78 | 0.86 |
| Inhibitory Kcnip2 | 47 | 83 | 0.57 |
| Inhibitory Pthlh | 16 | 95 | 0.17 |
| Inhibitory Vip | 55 | 126 | 0.44 |
| Microglia | 50 | 52 | 0.96 |
| Oligodendrocyte COP | 105 | 142 | 0.74 |
| Oligodendrocyte MF | 90 | 97 | 0.93 |
| Oligodendrocyte Mature | 348 | 381 | 0.91 |
| Oligodendrocyte NF | 56 | 76 | 0.74 |
| Oligodendrocyte Precursor cells | 102 | 111 | 0.92 |
| Pericytes | 22 | 94 | 0.23 |
| Perivascular Macrophages | 4 | 66 | 0.06 |
| Pyramidal Cpne5 | 62 | 76 | 0.82 |
| Pyramidal Kcnip2 | 15 | 25 | 0.6 |
| Pyramidal L2-3 | 163 | 172 | 0.95 |
| Pyramidal L2-3 L5 | 276 | 288 | 0.96 |
| Pyramidal L3-4 | 107 | 138 | 0.78 |
| Pyramidal L5 | 121 | 153 | 0.79 |
| Pyramidal L6 | 390 | 400 | 0.98 |
| Vascular Smooth Muscle | 5 | 31 | 0.16 |
| pyramidal L4 | 385 | 483 | 0.8 |

### Supplementary Table 2. The matching score between the osmFISH PolyA segmented cells vs. SSAM cell-type map guided by the scRNA-seq cluster centroids for selected cell types based on high gene expression correlation.

| **osmFISH cell type** | **scRNA-seq cell type** | **Pearson’s r** | **Matched Segments** | **Total Segments** | **Matching score** |
| --- | --- | --- | --- | --- | --- |
| Astrocyte Mfge8 | Astro2 | 0.83 | 103 | 119 | 0.87 |
| Oligodendrocyte COP | COP | 0.8 | 109 | 142 | 0.77 |
| Inhibitory Vip | Int10 | 0.83 | 14 | 126 | 0.11 |
| Inhibitory IC | Int13 | 0.91 | 64 | 78 | 0.82 |
| Inhibitory Crhbp | Int2 | 0.88 | 84 | 120 | 0.7 |
| Inhibitory Cnr1 | Int5 | 0.94 | 25 | 40 | 0.62 |
| Microglia | Mgl1 | 0.81 | 48 | 52 | 0.92 |
| Oligodendrocyte NF | NFOL1 | 0.86 | 53 | 76 | 0.7 |
| Oligodendrocyte Precursor cells | OPC | 0.91 | 102 | 111 | 0.92 |
| Pericytes | PPR | 0.89 | 12 | 94 | 0.13 |
| Perivascular Macrophages | Pvm1 | 0.81 | 5 | 66 | 0.08 |
| Pyramidal L2-3 L5 | S1PyrL23 | 0.92 | 286 | 288 | 0.99 |
| pyramidal L4 | S1PyrL5a | 0.93 | 294 | 483 | 0.61 |
| Endothelial | Vend2 | 0.83 | 153 | 235 | 0.65 |
| Vascular Smooth Muscle | Vsmc | 0.83 | 4 | 31 | 0.13 |

### Supplementary Table 3. The matching score between the osmFISH PolyA segmented cells vs. SSAM de novo cell-type map for selected cell types based on high gene expression correlation.

| **osmFISH cell type** | **De novo cell type** | **Pearson’s r** | **Matched Segments** | **Total Segments** | **Matching score** |
| --- | --- | --- | --- | --- | --- |
| Astrocyte Gfap | Astrocyte Gfap | 0.96 | 77 | 81 | 0.95 |
| Endothelial | Endothelial | 0.96 | 144 | 235 | 0.61 |
| Oligodendrocyte COP | Oligodendrocyte COP | 0.98 | 108 | 142 | 0.76 |
| Oligodendrocyte NF | Oligodendrocyte NF | 0.97 | 56 | 76 | 0.74 |
| Microglia | Microglia | 0.93 | 49 | 52 | 0.94 |
| Pyramidal Cpne5 | Pyramidal Cpne5 | 0.95 | 55 | 76 | 0.72 |
| Vascular Smooth Muscle | Vascular Smooth Muscle | 0.93 | 5 | 31 | 0.16 |
| Astrocyte Mfge8 | Astrocyte Mfge8 | 0.94 | 107 | 119 | 0.9 |
| pyramidal L4 | pyramidal L4 | 0.97 | 400 | 483 | 0.83 |
| Pyramidal L3-4 | Pyramidal L3-4 | 0.95 | 106 | 138 | 0.77 |
| Inhibitory Kcnip2 | Inhibitory Pthlh | 0.8 | 29 | 83 | 0.35 |
| Inhibitory Crhbp | Inhibitory Crhbp | 0.96 | 102 | 120 | 0.85 |
| Pyramidal L2-3 L5 | Pyramidal L2-3 L5 | 0.98 | 282 | 288 | 0.98 |
| Oligodendrocyte Precursor cells | Oligodendrocyte Precursor cells | 0.98 | 102 | 111 | 0.92 |
| Pericytes | Pericytes | 0.84 | 21 | 94 | 0.22 |
| Inhibitory IC | Inhibitory Rest | 0.95 | 64 | 78 | 0.82 |
| Pyramidal L6 | Pyramidal L6 | 0.96 | 388 | 400 | 0.97 |
| Oligodendrocyte Mature | Oligodendrocyte Mature | 0.97 | 341 | 381 | 0.9 |
| Oligodendrocyte MF | Oligodendrocyte MF | 0.98 | 87 | 97 | 0.9 |
| Ependymal | Ependymal | 0.98 | 95 | 107 | 0.89 |
| Pyramidal L2-3 | Pyramidal L2-3 | 0.97 | 138 | 172 | 0.8 |
| Inhibitory Cnr1 | Inhibitory Cnr1 | 0.97 | 30 | 40 | 0.75 |
| Inhibitory CP | Inhibitory Kcnip2 | 0.82 | 95 | 161 | 0.59 |
| Hippocampus | Hippocampus | 0.93 | 98 | 133 | 0.74 |

### Supplementary Table 4. The matching score between the MERFISH segmented cells vs. SSAM cell-type map guided by the segmentation-based cell-type signature of MERFISH data.

| **Cell type** | **Matched segments** | **Total segments** | **Matching score** |
| --- | --- | --- | --- |
| Astrocyte | 740 | 745 | 0.99 |
| E-1 | 14 | 15 | 0.93 |
| E-10 | 7 | 14 | 0.5 |
| E-11 | 5 | 20 | 0.25 |
| E-12 | 12 | 35 | 0.34 |
| E-13 | 100 | 117 | 0.85 |
| E-14 | 240 | 298 | 0.81 |
| E-15 | 44 | 56 | 0.79 |
| E-16 | 127 | 145 | 0.88 |
| E-17 | 22 | 29 | 0.76 |
| E-18 | 69 | 123 | 0.56 |
| E-19 | 48 | 50 | 0.96 |
| E-2 | 4 | 6 | 0.67 |
| E-20 | 6 | 8 | 0.75 |
| E-21 | 5 | 6 | 0.83 |
| E-22 | 0 | 5 | 0 |
| E-23 | 25 | 27 | 0.93 |
| E-24 | 27 | 41 | 0.66 |
| E-25 | 2 | 3 | 0.67 |
| E-26 | 1 | 1 | 1 |
| E-28 | 19 | 19 | 1 |
| E-29 | 1 | 1 | 1 |
| E-3 | 13 | 18 | 0.72 |
| E-30 | 1 | 1 | 1 |
| E-31 | 2 | 2 | 1 |
| E-4 | 27 | 35 | 0.77 |
| E-5 | 1 | 1 | 1 |
| E-6 | 16 | 24 | 0.67 |
| E-7 | 47 | 79 | 0.59 |
| E-8 | 14 | 18 | 0.78 |
| E-9 | 19 | 19 | 1 |
| Endothelial 1 | 290 | 301 | 0.96 |
| Endothelial 2 | 17 | 19 | 0.89 |
| Endothelial 3 | 112 | 117 | 0.96 |
| Ependymal | 219 | 219 | 1 |
| H-1 | 1 | 1 | 1 |
| I-1 | 1 | 352 | 0 |
| I-10 | 15 | 27 | 0.56 |
| I-11 | 66 | 107 | 0.62 |
| I-12 | 110 | 139 | 0.79 |
| I-13 | 226 | 316 | 0.72 |
| I-14 | 75 | 89 | 0.84 |
| I-15 | 165 | 174 | 0.95 |
| I-16 | 77 | 99 | 0.78 |
| I-18 | 116 | 135 | 0.86 |
| I-19 | 5 | 11 | 0.45 |
| I-2 | 77 | 120 | 0.64 |
| I-20 | 10 | 10 | 1 |
| I-21 | 3 | 24 | 0.12 |
| I-22 | 6 | 10 | 0.6 |
| I-23 | 70 | 75 | 0.93 |
| I-24 | 12 | 15 | 0.8 |
| I-25 | 26 | 44 | 0.59 |
| I-26 | 1 | 1 | 1 |
| I-27 | 11 | 12 | 0.92 |
| I-29 | 6 | 10 | 0.6 |
| I-3 | 21 | 47 | 0.45 |
| I-30 | 5 | 6 | 0.83 |
| I-31 | 23 | 23 | 1 |
| I-33 | 3 | 3 | 1 |
| I-34 | 5 | 5 | 1 |
| I-37 | 1 | 1 | 1 |
| I-4 | 16 | 17 | 0.94 |
| I-5 | 5 | 5 | 1 |
| I-6 | 6 | 13 | 0.46 |
| I-7 | 142 | 245 | 0.58 |
| I-8 | 26 | 31 | 0.84 |
| I-9 | 30 | 50 | 0.6 |
| Microglia | 99 | 106 | 0.93 |
| OD Immature 1 | 222 | 223 | 1 |
| OD Immature 2 | 1 | 7 | 0.14 |
| OD Mature 1 | 26 | 49 | 0.53 |
| OD Mature 2 | 221 | 241 | 0.92 |
| OD Mature 3 | 5 | 6 | 0.83 |
| OD Mature 4 | 7 | 10 | 0.7 |
| Pericytes | 60 | 62 | 0.97 |

### Supplementary Table 5. The matching score between the MERFISH segmented cells vs. SSAM cell-type map guided by scRNA-seq cluster centroids for selected cell types based on high gene expression correlation.

| **MERFISH cell type** | **scRNA-seq cell type** | **Pearson’s r** | **Matched segments** | **Total segments** | **Matching score** |
| --- | --- | --- | --- | --- | --- |
| Astrocyte | Astrocytes 2 | 0.91 | 567 | 745 | 0.76 |
| Endothelial 3 | Endothelial 3 | 0.9 | 84 | 117 | 0.72 |
| E-5 | Excitatory (e8:Cck/Ebf3) | 0.86 | 1 | 1 | 1 |
| OD Immature 1 | Immature oligodendrocyte 1 | 0.9 | 221 | 223 | 0.99 |
| Microglia | Microglia 2 | 0.97 | 100 | 106 | 0.94 |

### Supplementary Table 6. The matching score between the MERFISH segmented cells vs. SSAM de novo cluster cell-type map for selected cell types based on high gene expression correlation.

| **MERFISH cell type** | **De novo cell type** | **Pearson’s r** | **Matched segments** | **Total segments** | **Matching score** |
| --- | --- | --- | --- | --- | --- |
| Astrocyte | Astrocyte 1 | 0.94 | 709 | 745 | 0.95 |
| OD Immature 1 | Immature OD | 0.92 | 220 | 223 | 0.99 |
| I-31 | Inhibitory Amigo2,Sema3c | 0.91 | 23 | 23 | 1 |
| I-27 | Inhibitory Gad1 | 0.93 | 10 | 12 | 0.83 |
| E-14 | Excitatory Syt4 | 0.88 | 175 | 298 | 0.59 |
| E-19 | Excitatory Cbln1,Cbln2 | 0.85 | 48 | 50 | 0.96 |
| OD Mature 2 | Mature OD 2 | 0.97 | 240 | 241 | 1 |
| Pericytes | Mural | 0.96 | 60 | 62 | 0.97 |
| E-9 | Excitatory Necab1,Gda | 0.92 | 16 | 19 | 0.84 |
| I-19 | Inhibitory Gpr165 | 0.8 | 6 | 11 | 0.55 |
| I-7 | Inhibitory Amigo2 | 0.85 | 129 | 245 | 0.53 |
| I-18 | Inhibitory Col25a1 | 0.89 | 72 | 135 | 0.53 |
| Endothelial 3 | Endothelial 3 | 0.98 | 108 | 117 | 0.92 |
| Microglia | Microglia | 0.98 | 99 | 106 | 0.93 |
| E-5 | Excitatory Ebf3 | 0.86 | 1 | 1 | 1 |
| Endothelial 1 | Endothelial 1 | 0.93 | 239 | 301 | 0.79 |
| Ependymal | Ependymal | 0.92 | 219 | 219 | 1 |
| I-15 | Inhibitory Irs4 | 0.88 | 112 | 174 | 0.64 |

### Supplementary Table 7. Full and abbreviated cell-type annotation of VISp smFISH data

| **Full name** | **Abbreviated name** |
| --- | --- |
| L2/3 IT Rrad | L2/3 IT 1 |
| L2/3 IT Agmat | L2/3 IT 2 |
| L4 IT Rspo1 | L4 IT 1 |
| L4 IT Superficial | L4 IT 2 |
| L5 IT Hsd11b1 Endou | L5 IT 1 |
| L5 IT Batf3 | L5 IT 2 |
| L5 IT Col27a1 | L5 IT 3 |
| L6 IT Penk Col27a1 | L6 IT 1 |
| L6 IT Col23a1 Adamts2 | L6 IT 2 |
| L5 PT Chrna6 | L5 PT 1 |
| L5 PT C1ql2 Cdh13 | L5 PT 2 |
| L5 PT Krt80 | L5 PT 3 |
| L5 NP Trhr Met | L5 NP |
| L6 CT Ctxn3 Brinp3 | L6 CT |
| L6b Col8a1 Rprm | L6b 1 |
| L6b Crh | L6b 2 |
| Lamp5 Lsp1 | Lamp5 |
| Vip Arhgap36 Hmcn1 / Vip Igfbp4 Map21l1 | Vip |
| Sst Chodl | Sst 1 |
| Sst Crhr2 Efemp1 | Sst 2 |
| Sst Nts / Sst Rxfp1 Eya1 | Sst 3 |
| Pvalb Th Sst / Pvalb Reln Tac1 | Pvalb 1 |
| Pvalb Calb1 Sst / Pvalb Sema3e Kank4 | Pvalb 2 |
| Pvalb Gpr149 Islr / Pvalb Reln Tac1 | Pvalb 3 |
| Astro Aqp4 | Astro |
| OPC | OPC |
| Oligo | Oligo |
| VLMC | VLMC |

### Supplementary Table 8. Doublet rates reported by double detection methods, DoubletDetection and Scrublet.

|  | SSp (osmFISH) | POA (MERFISH) | VISp (Multiplexed smFISH) |
| --- | --- | --- | --- |
| DoubletDetection | 0.91% | 3.4% | 0.47% |
| Scrublet | 0% | 2.6% | 0.5% |

### Supplementary Figure 1. Identification of intracellular regions via mRNA density estimation.

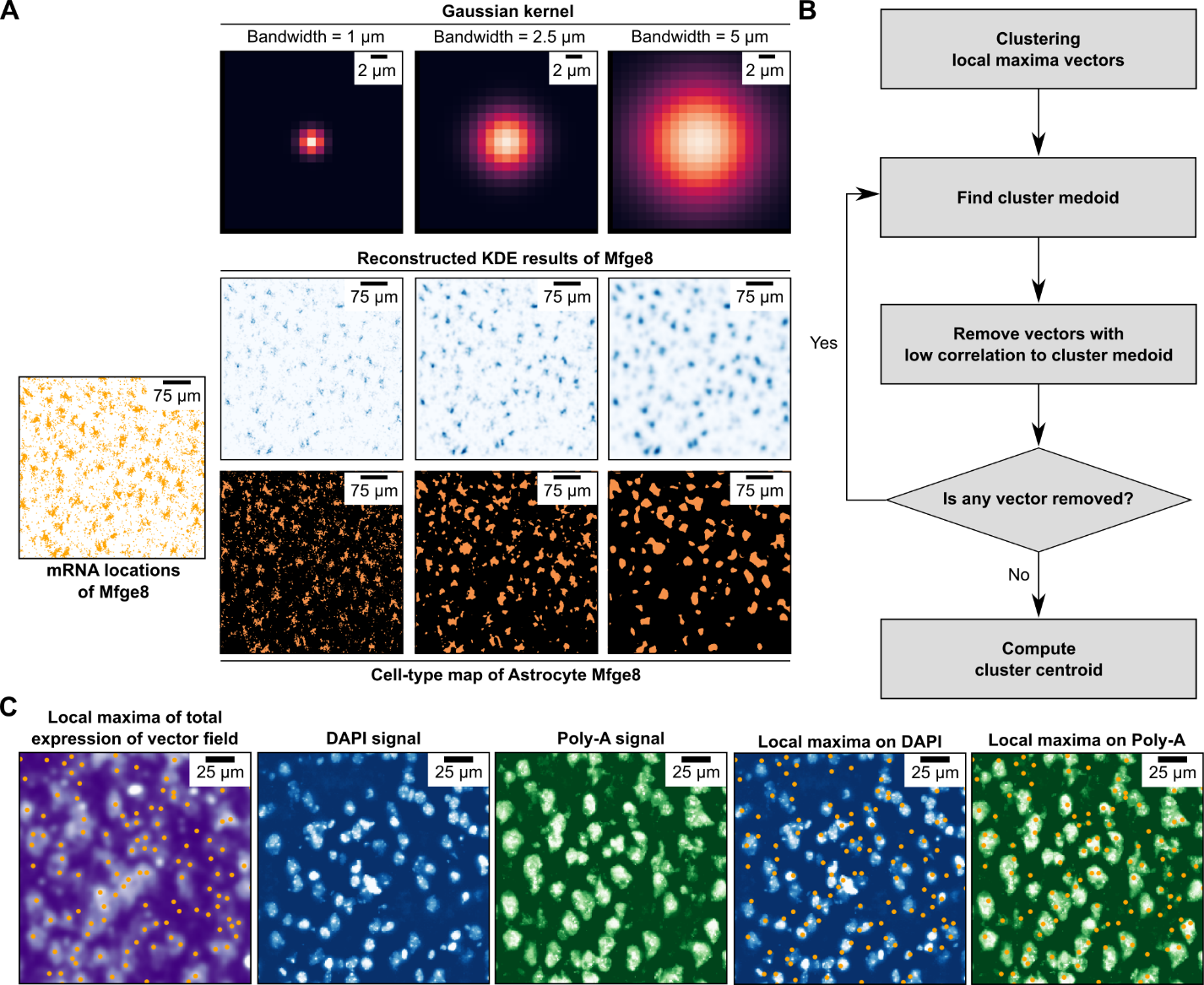

(A) Depiction of KDE reconstructed result of mRNA locations of single cell and multiple cells using Gaussian kernels with different bandwidth. With too low bandwidth (1 μm), KDE fails to reconstruct the shape of cells. With high bandwidth (5 μm), KDE loses detailed structure of cell shape (second row), but it does not have so much effect on the resolution of the corresponding cell type map (third row, see also Supplementary Figure 8).

(B) Flowchart describing selection criteria of local maxima vectors leading to cell-type signatures.

(C) Example of local maxima location in relation to cell markers (from the mouse SSp image). From left to right: local maxima (orange dots), correspond to highest local intensity of the gene expression vector field; DAPI signal over the corresponding area; Poly-A signal over the corresponding area; overlay of local maxima positions with DAPI shows many local maxima fall within or directly adjacent to strong DAPI signal; overlap of local maxima positions with Poly-A signal shows most local maxima fall within areas of high Poly-A signal. Some of the local maxima originating outside of high Poly-A regions correspond to cell-type signals originating from cells with a lower total mRNA abundance, such as astrocytes.

### Supplementary Figure 2. Identification of cell-type and tissue domain signatures and maps.

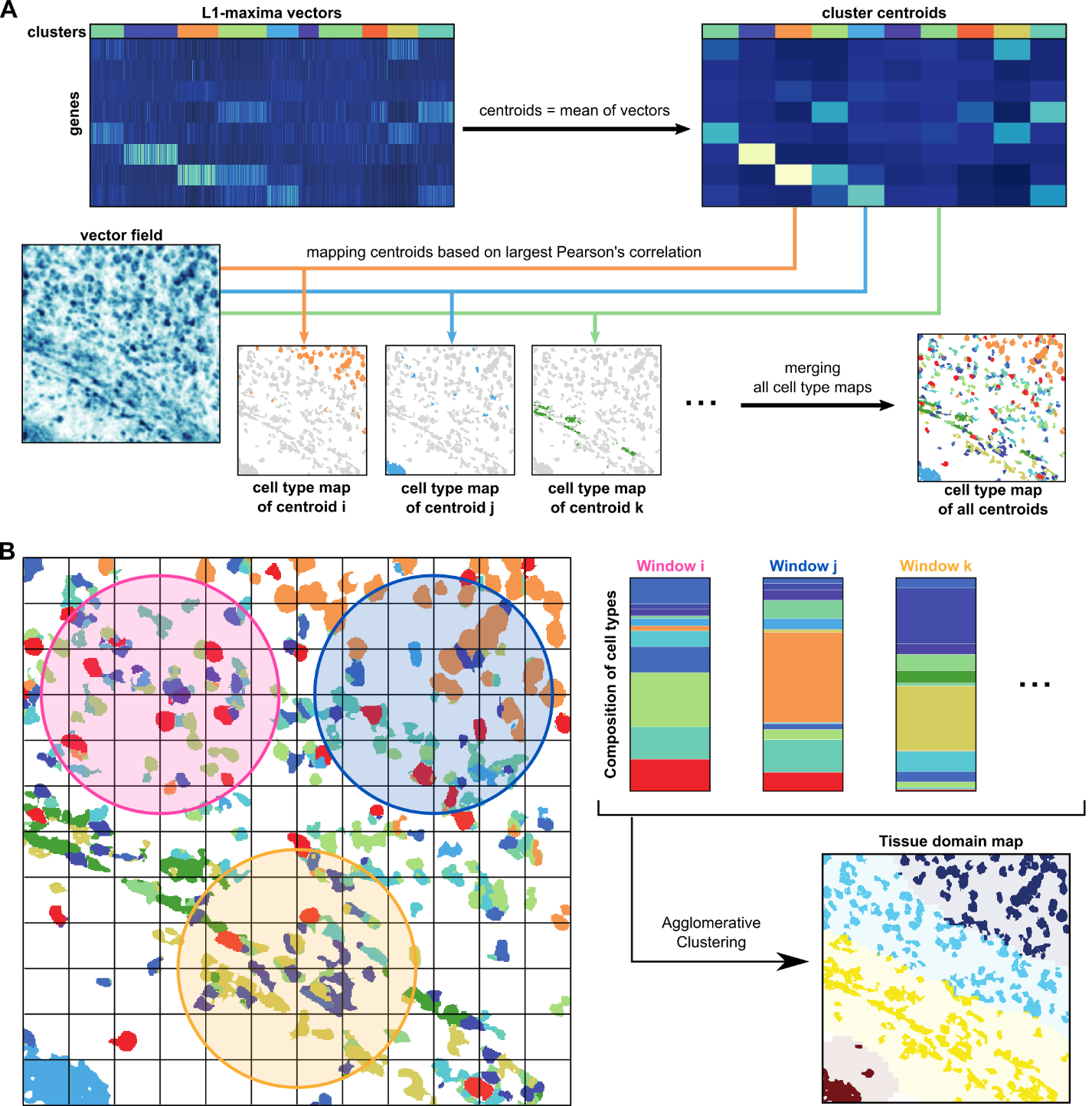

(A) Depiction of the creation of the cell-type map from the clustering of local maxima gene expression vectors. After the clustering, the mean gene expression vector (centroid) for each cluster is calculated and used as representative for the cell-type signatures. These signatures are mapped onto the vector field to construct the cell-type map, by calculating Pearson’s correlation between the centroids to the vector field and then assigning cell types based on maximum correlation.

(B) Depictions of the creation of the tissue domain map. A circular sliding window passes over all lattice points on the cell-type map, profiling the cell-type composition. These windows are clustered to create the tissue domain map.

### Supplementary Figure 3. Local maxima selection and filtering criteria in the mouse SSp osmFISH dataset.

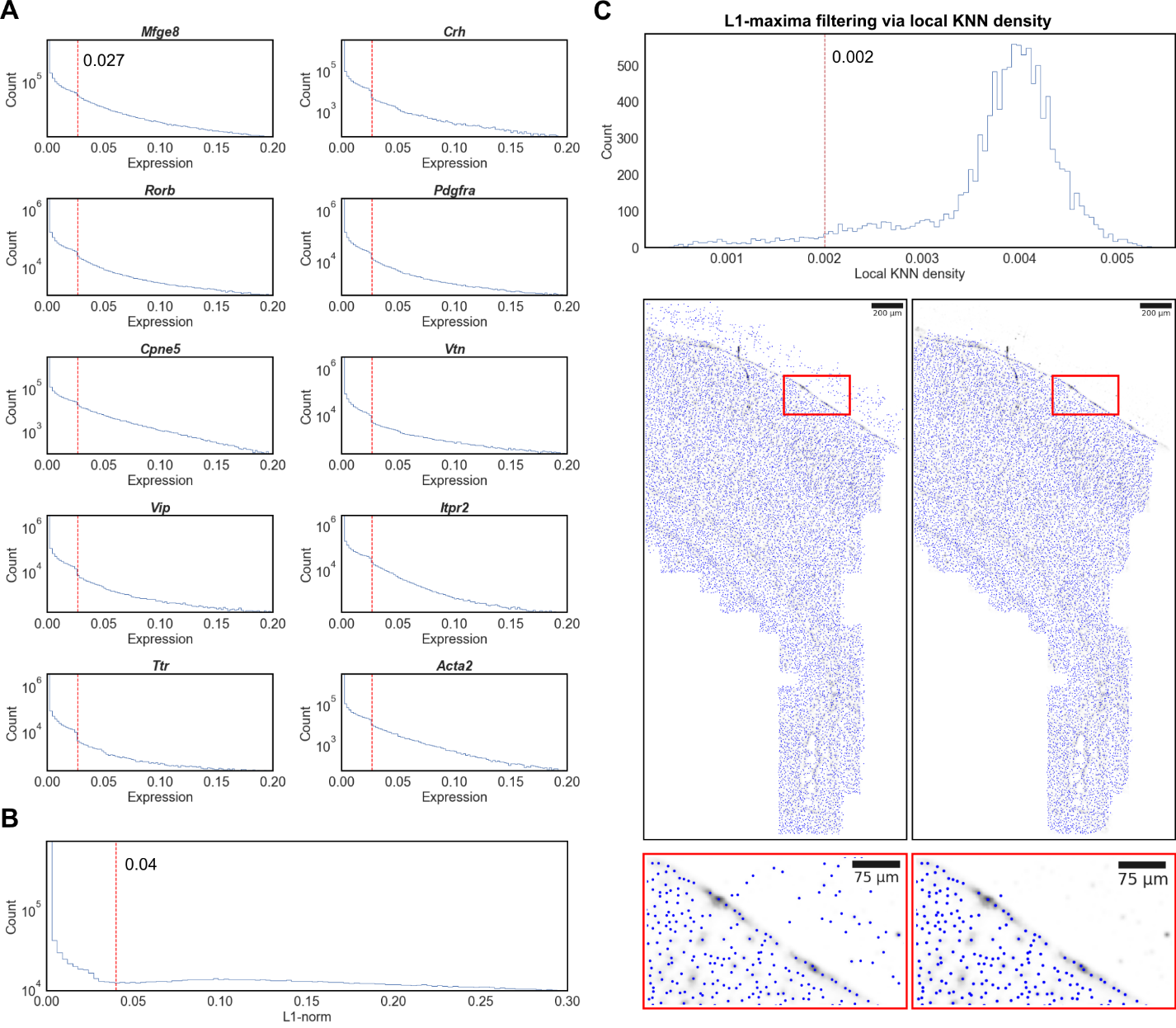

(A) The gene expression threshold corresponds to an observable drop in expression signal, shown for 10 randomly selected genes from the dataset (bottom). Local maxima are only selected if they exceed the determined gene expression threshold for at least 1 gene, so that at least 1 gene is sufficiently expressed.

(B) The total gene expression threshold (top, red vertical line). Local maxima are only selected if they exceed this threshold, preventing selection of local maxima with a low total gene expression signal.

(C) Local maxima filtering criteria. Local maxima are filtered based on local KNN density threshold local maxima (top, red vertical line) reducing spurious local maxima outside of the tissue region (bottom). Red boxed regions shows a zoomed-in region at the edge of the tissue, showing spurious local maxima originating from outside the tissue are effectively removed.

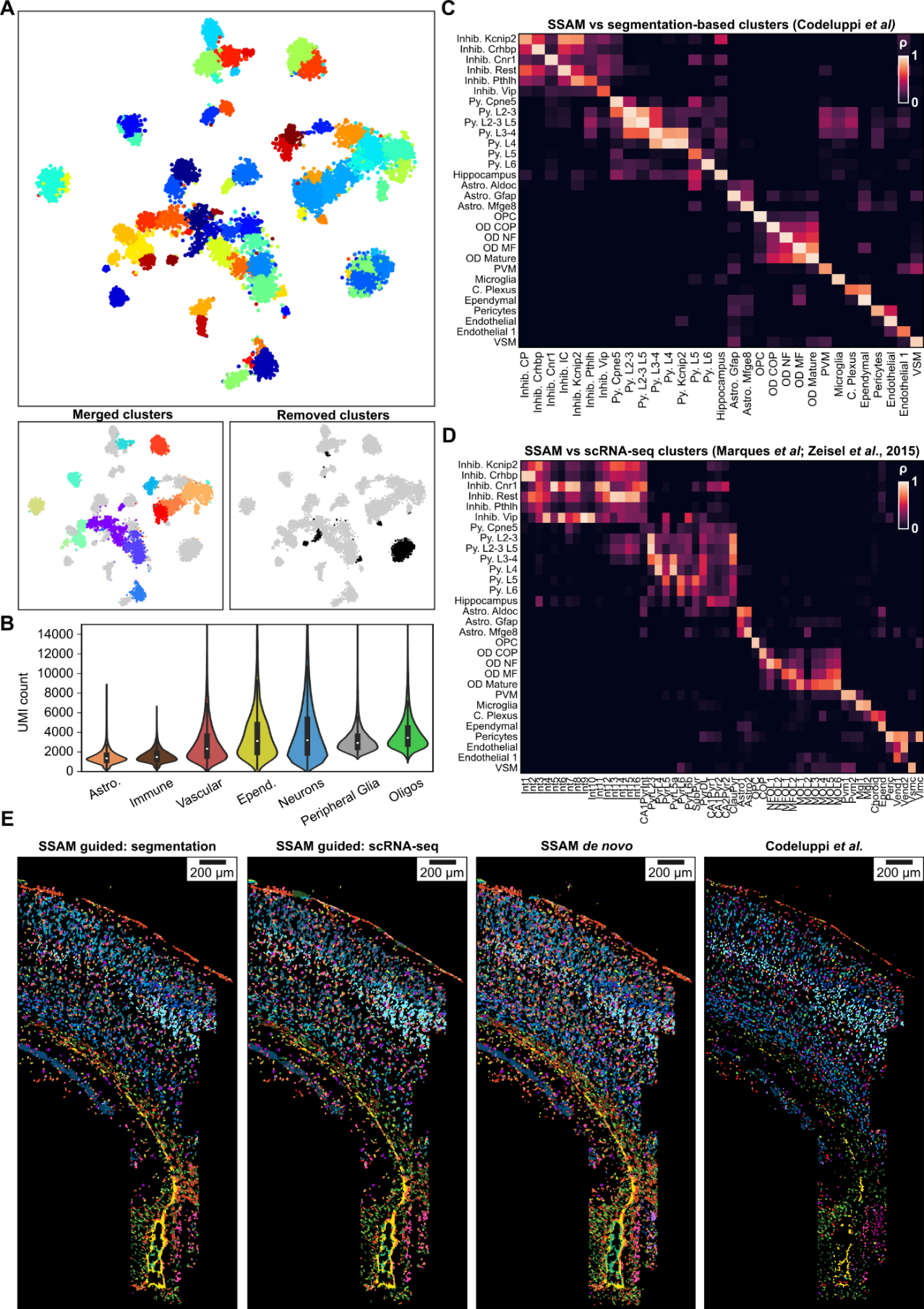

### Supplementary Figure 4. Cell-type signature identification and mapping in the mouse SSp osmFISH dataset.

(A) Clustering merging and filtering. A tSNE map of clustering all local maxima gene expression vectors (top). Clusters with similar expression profiles were merged (bottom left), and those mapping to artifact regions of the image, or containing highly variable gene expression signatures were removed (bottom right).

(B) Total cell-wise mRNA counts of different cell-type classes in the mouse brain. The violin plot shows UMI counts of ~500,000 single mouse brain cells profiled using scRNA-seq, grouped by cell class and ordered by increasing median of UMI counts. Cells from the ‘Astrocyte’ and ‘Immune’ classes show a significantly decreased UMI (all *p-values* < 1 x 10e-12), with half of the recorded cells of both classes exhibiting a lower UMI count than the lowest quartile of any other cell class. Both classes are underrepresented in the previous segmentation-based reconstruction compared to SSAM, indicating that SSAM shows higher sensitivity towards cells with low total mRNA content than segmentation-based approaches on poly-A stained images.

(C) Comparison of the SSAM *de novo* cell-type signatures to the segmentation-based cell-type signatures from Codeluppi *et al.*

(D) Comparison of the SSAM *de novo* cell-type signatures to the scRNA-seq derived cell-type signatures from Marques *et al* and Zeisel *et al.*

(E) Comparison of cell-type maps generated using SSAM guided and *de novo* mode. Cell- type maps generated using signatures derived from (left to right) segmentation from Codeluppi *et al.*, scRNAseq from Zeisel *et al.*, SSAM *de novo* signature identification, and the original cell map from Codeluppi *et al.* In all versions of the reconstruction by SSAM many more astrocytes are detected compared to original cell map. Color of scRNA-seq clusters were determined by calculating Pearson’s correlation coefficient between scRNA-seq and segmentation-based cluster centroids. The colors of the cell types correspond to the cell-type legend in Fig. 2B.

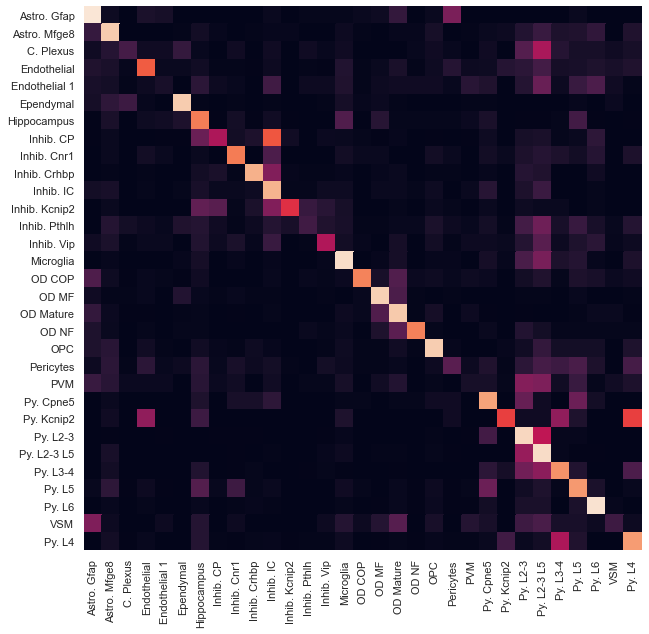

### Supplementary Figure 5. Pairwise matching scores heatmap between the guided mode cell type map (osmFISH segmentation-based) and osmFISH Poly-A segments.

### Supplementary Figure 6. Pairwise matching scores heatmap between the guided mode cell type map (osmFISH scRNA-seq based) and osmFISH Poly-A segments.

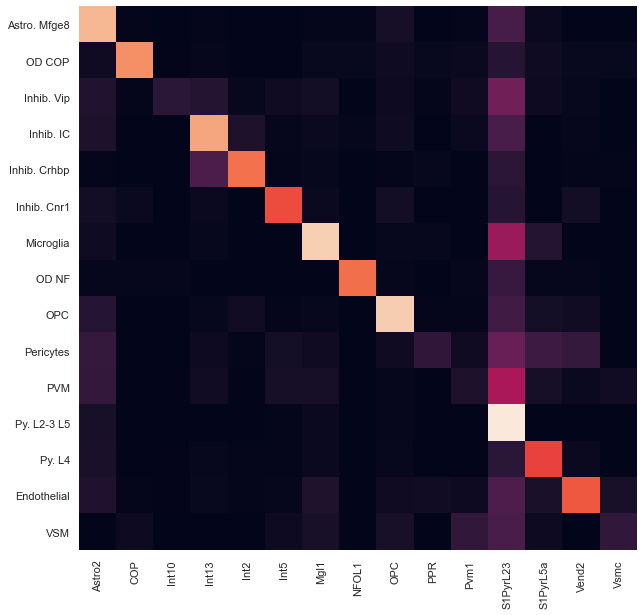

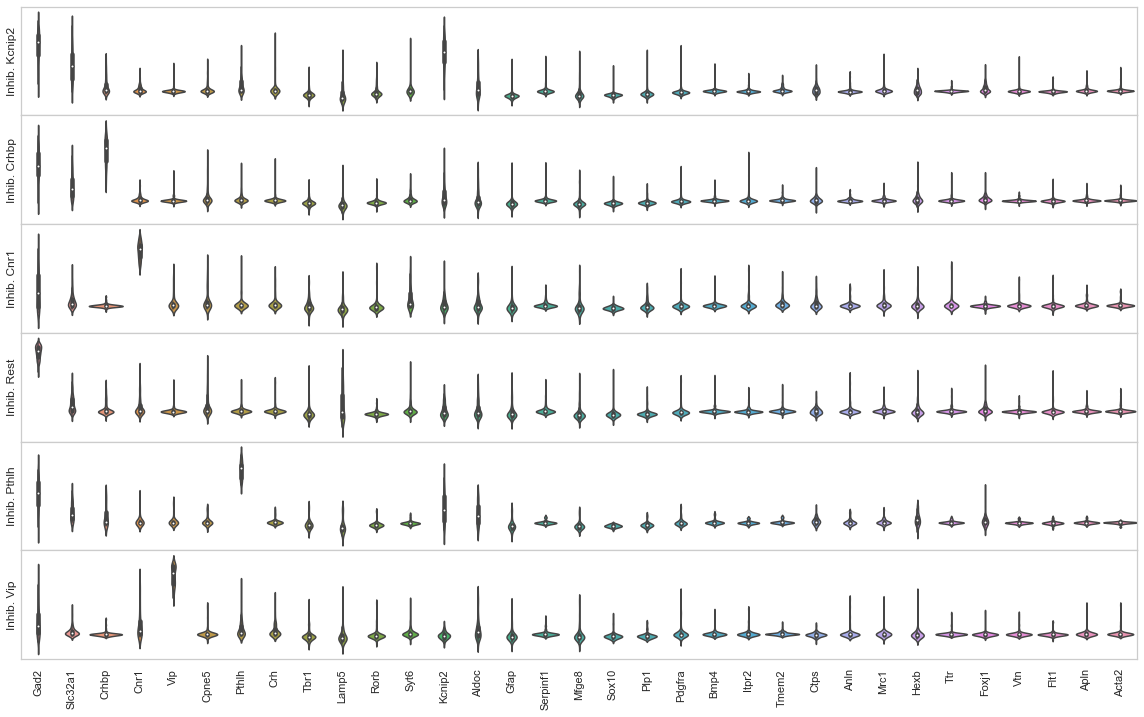

### Supplementary Figure 7. Gene expression violin plots of local maxima, osmFISH inhibitory neuronal cell types.

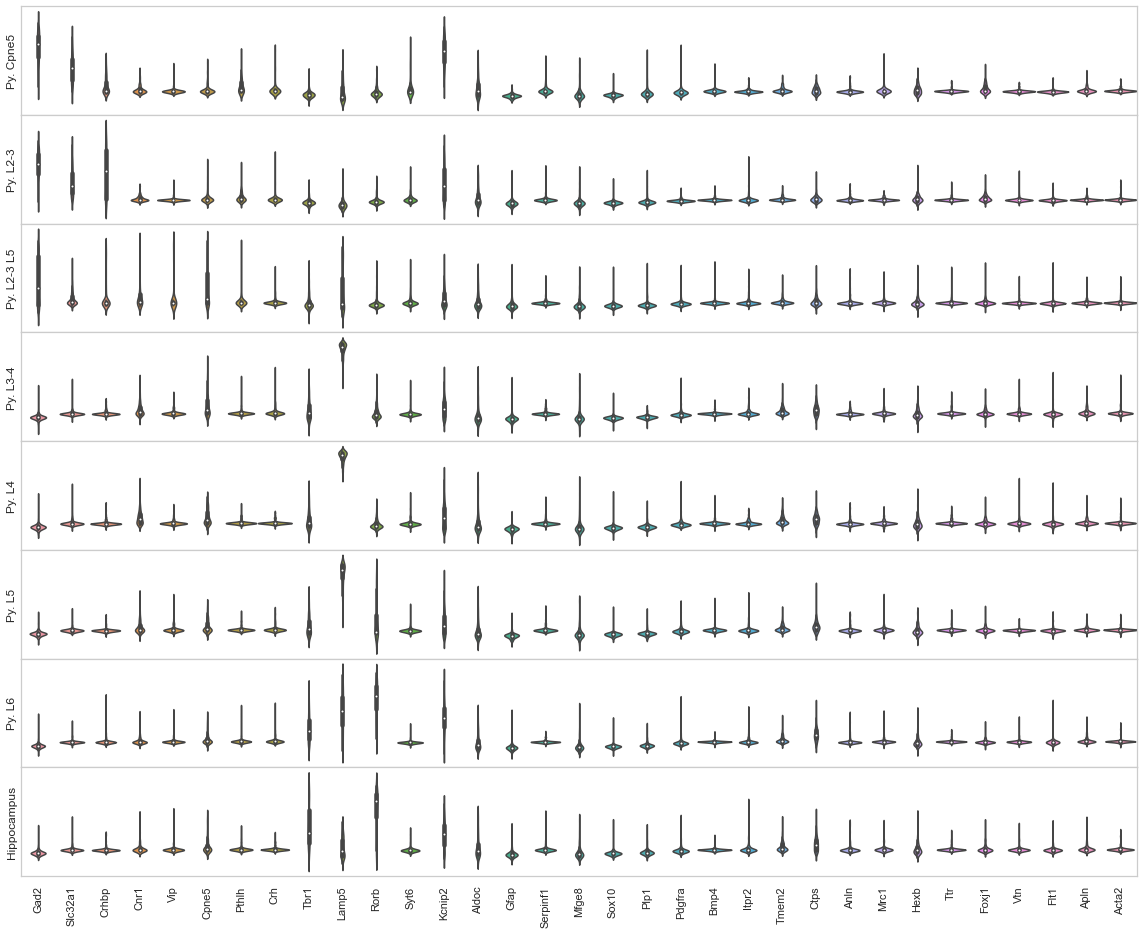

### Supplementary Figure 8. Gene expression violin plots of local maxima, osmFISH pyramidal and hippocampus cell types.

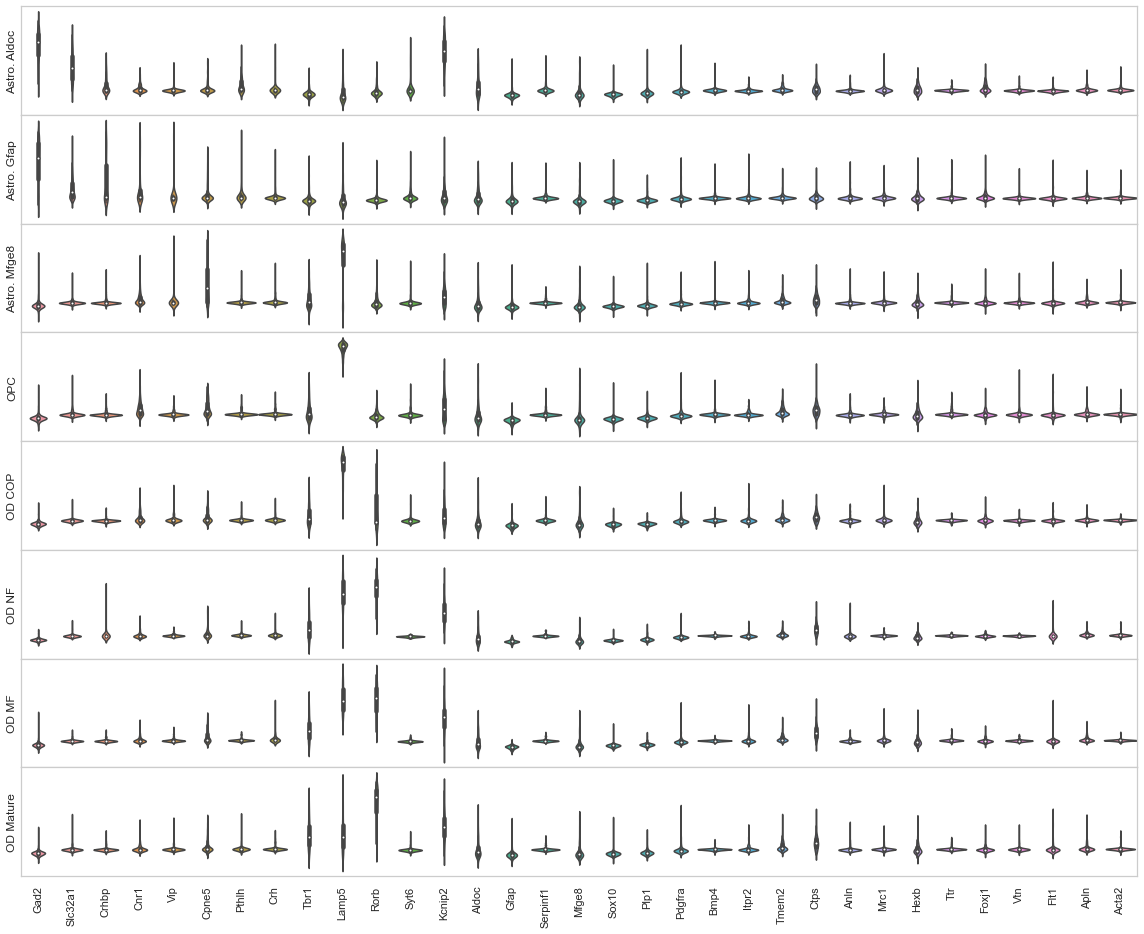

### Supplementary Figure 9. Gene expression violin plots of local maxima, osmFISH oligodendrocyte and astrocyte cell types.

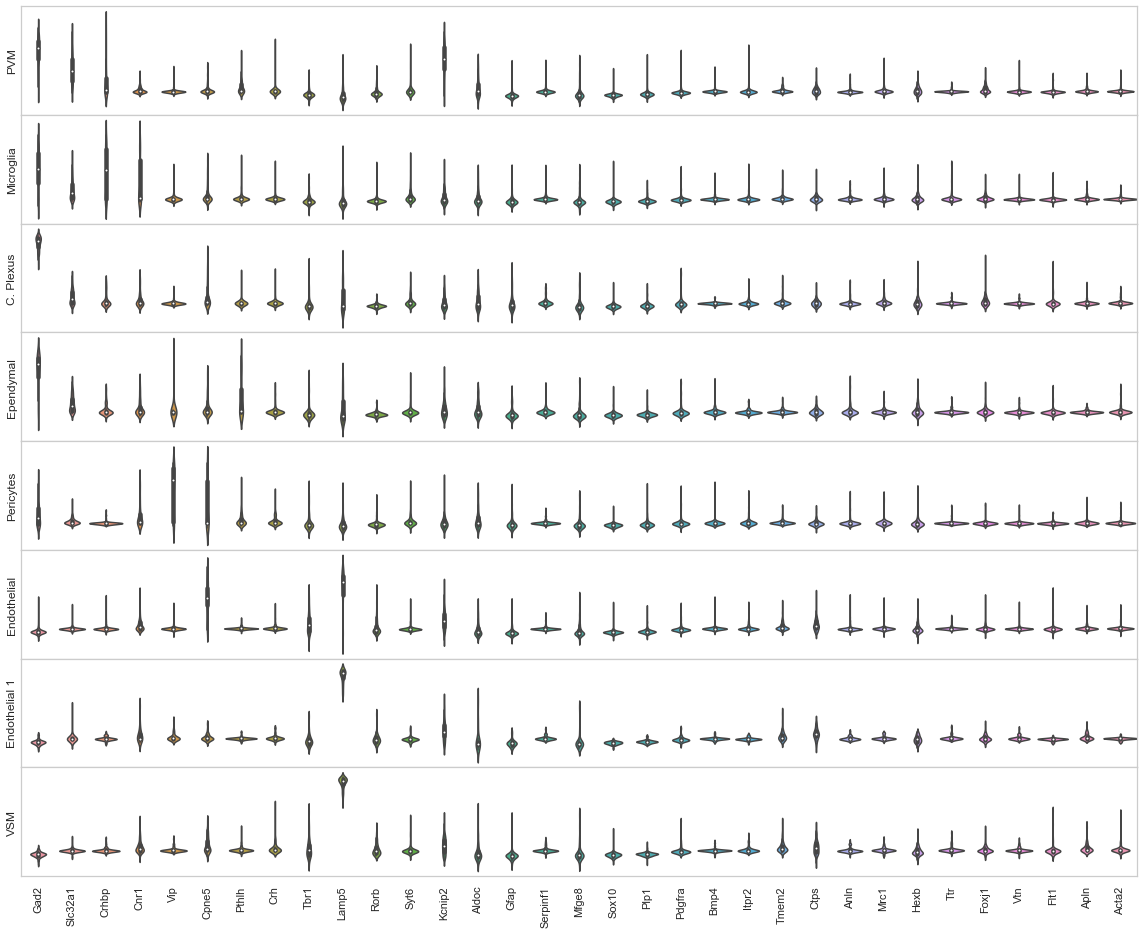

### Supplementary Figure 10. Gene expression violin plots of local maxima, osmFISH other cell types.

### Supplementary Figure 11. Pairwise matching scores heatmap between the guided mode cell type map (osmFISH scRNA-seq based) and osmFISH Poly-A segments.

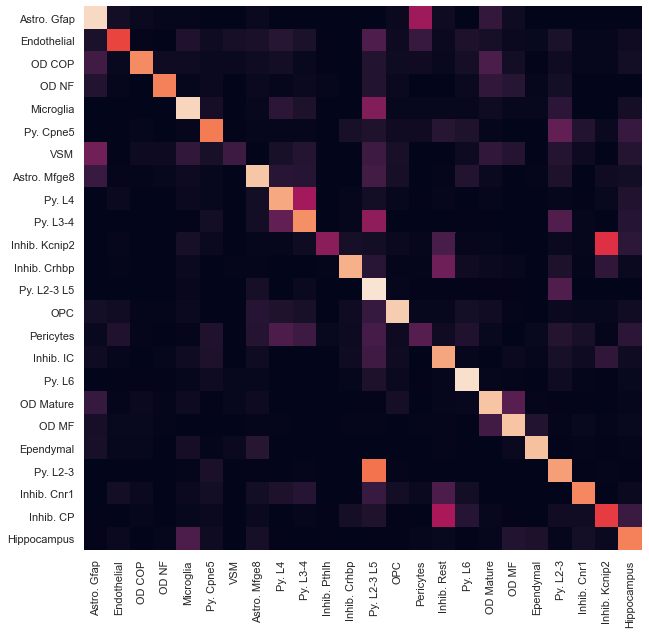

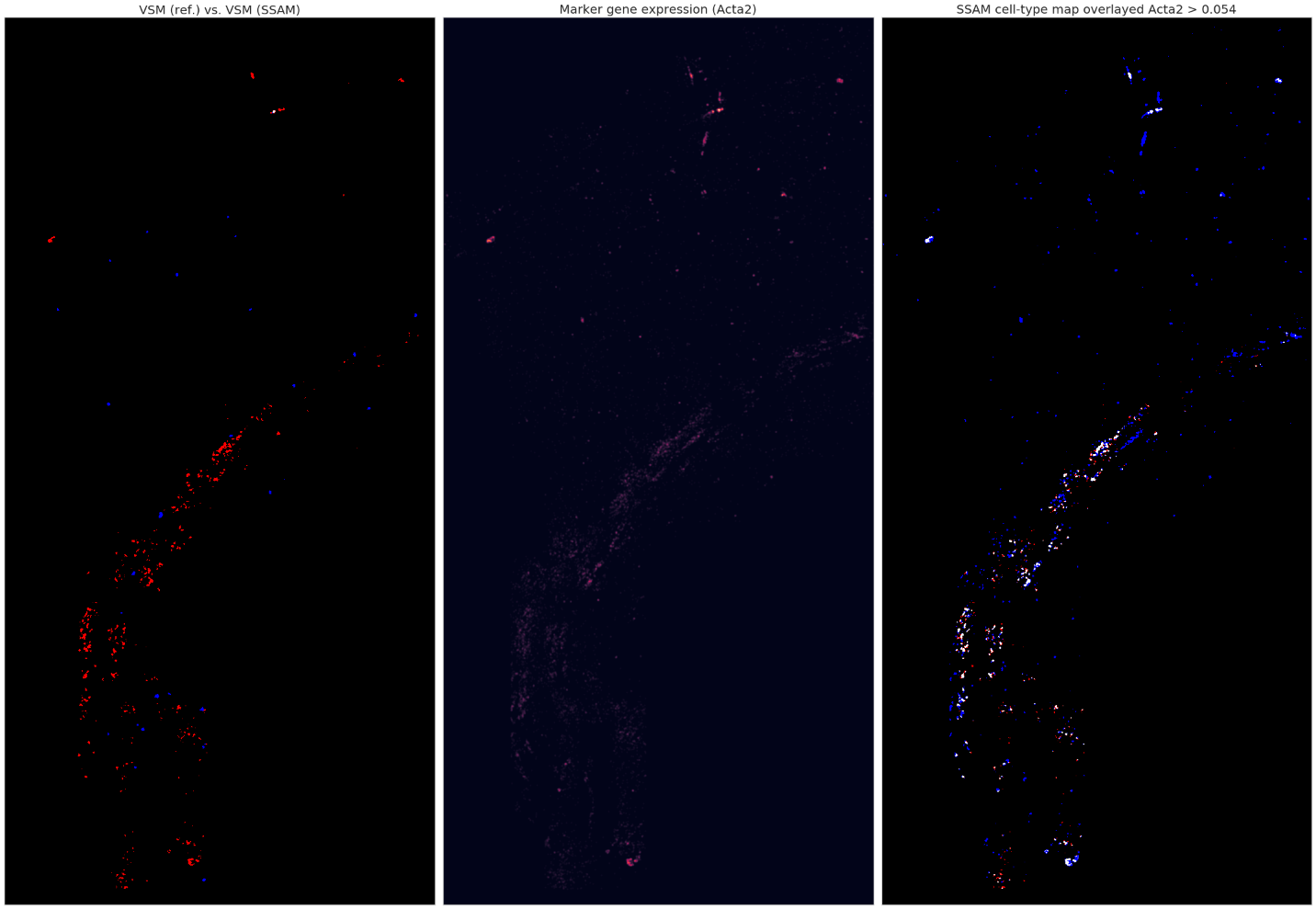

### Supplementary Figure 12. Comparison of the cell type (Vascular Smooth Muscle) and the corresponding marker gene expression (Acta2) of osmFISH data.

Vascular Smooth Muscle was found to have a low matching score between the SSAM *de* novo and the original cell-based segmentation cell-type map. (Left panel) red indicates the regions labelled with this cell type in the SSAM cell-type map, blue indicates the cells labelled with this cell type in the cell-based segmentation cell-type map, and white indicates their overlap. (Middle panel) the marker gene expression of the corresponding cell type (Acta2). (Right panel) red indicates SSAM cell-type map, blue indicates the marker gene expression above threshold 0.054, and white indicates their overlap. The marker gene expression better supports the SSAM cell-type calls compared to the segmentation-based cell-type calls.

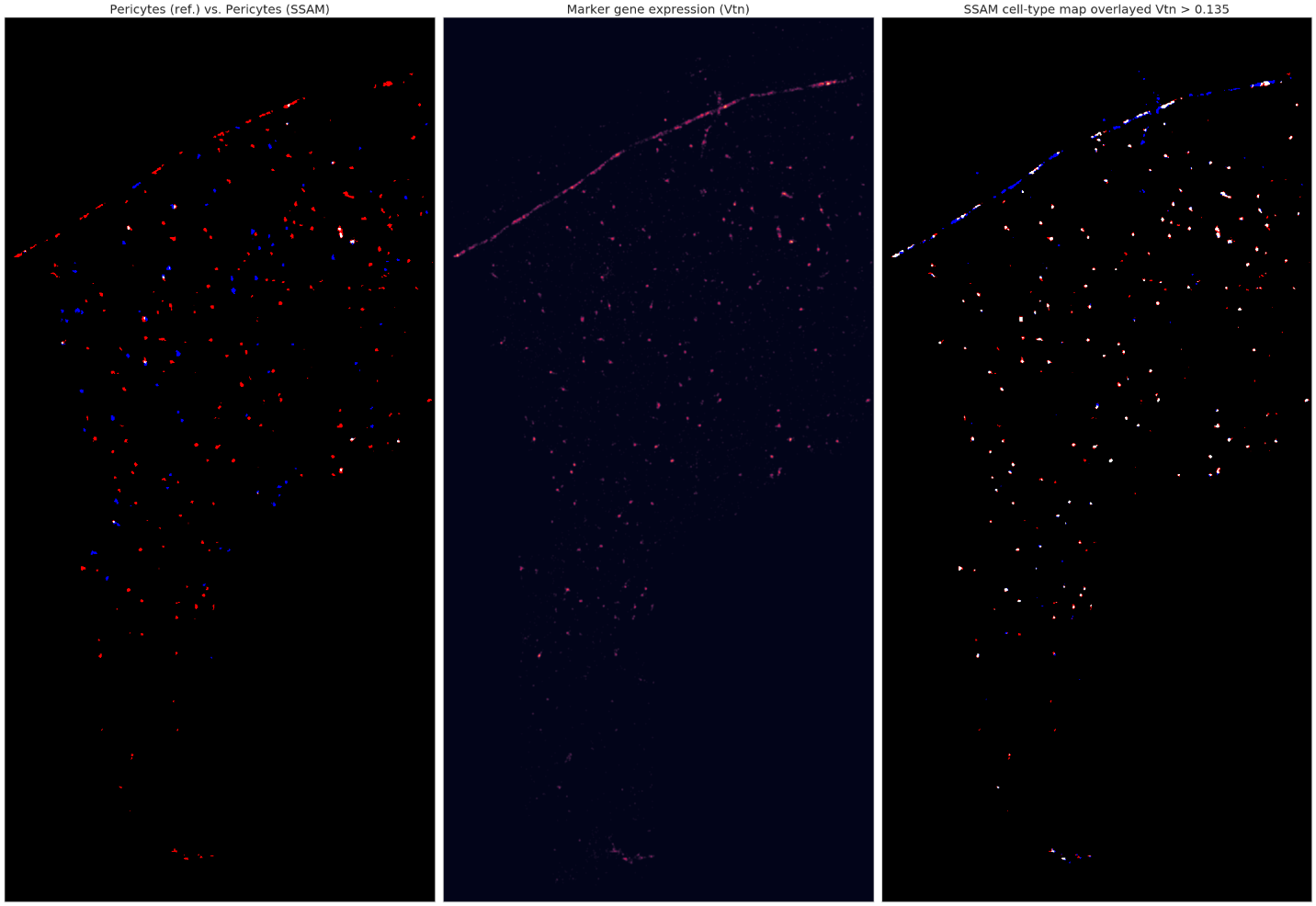

### Supplementary Figure 13. Comparison of the cell type (Pericytes) and the corresponding marker gene expression (Vtn) of osmFISH data.

Pericytes were found to have a low matching score between the SSAM *de* novo and the original cell-based segmentation cell-type map. (Left panel) red indicates the regions labelled with this cell type in the SSAM cell-type map, blue indicates the cells labelled with this cell type in the cell-based segmentation cell-type map, and white indicates their overlap. (Middle panel) the marker gene expression of the corresponding cell type (Vtn). (Right panel) red indicates SSAM cell-type map, blue indicates the marker gene expression above threshold 0.135, and white indicates their overlap. The marker gene expression better supports the SSAM cell-type calls compared to the segmentation-based cell-type calls.

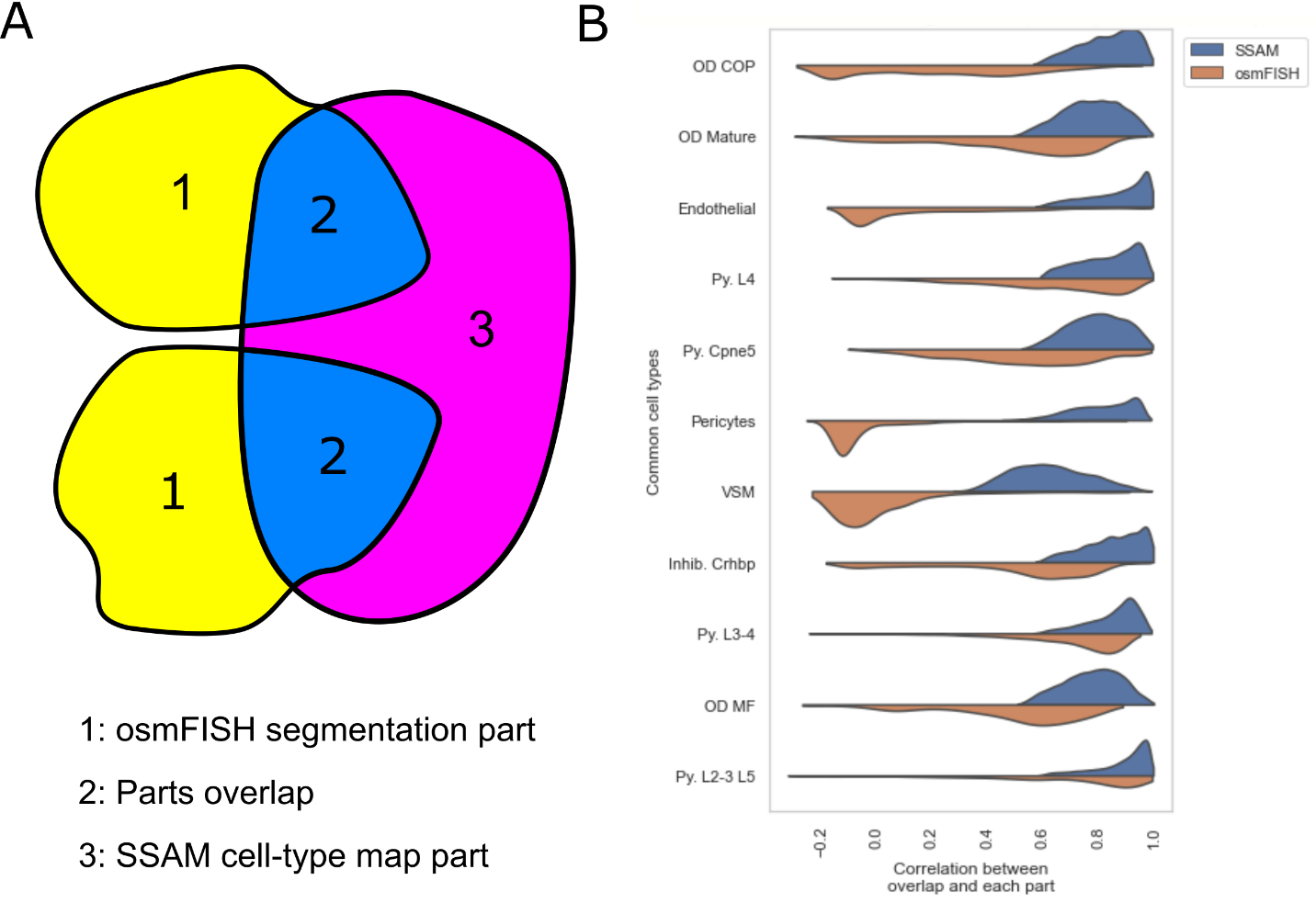

### Supplementary Figure 14. Overlap-based validation of the osmFISH cell-type map.

The osmFISH segmentation (1) is overlayed with the SSAM *de novo* cell-type map (3), and their overlap (2) (A); The correlations between the overlap and each part of the cell-type map unique to either SSAM (blue) or osmFISH segmentation (orange) reveals that the parts unique to the SSAM cell-type map better correlate with the gene expression in shared parts, compared to the parts unique to the osmFISH segmentation (B). The common cell-types can be found in Supplementary Table 3.

### Supplementary Figure 15. Laminar distribution of layer-specific pyramidal cell types in the mouse SSp.

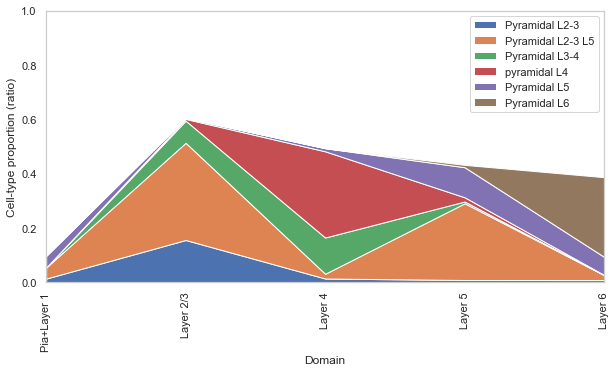

An established ground truth in neurobiology is the presence of various pyramidal cells in the cortical layers of the mouse brain. We validate our cell-type map and domain map by showing that: layer 2/3 primarily consists of Pyramidal L2-3/L5, L2-3, and L3-4 cell types; layer 4 consists of Pyramidal L4 and L3-4 cell types; layer 5 consists of Pyramidal L3-5 and L5 cell types; and layer 6 consists of Pyramidal L6 cell types. Other cell types, that are not show, contribute to the rest of the cell-type proportions (shown as white space).

### Supplementary Figure 16. MERFISH gene expression clustering heatmap with vertical cell type separations.

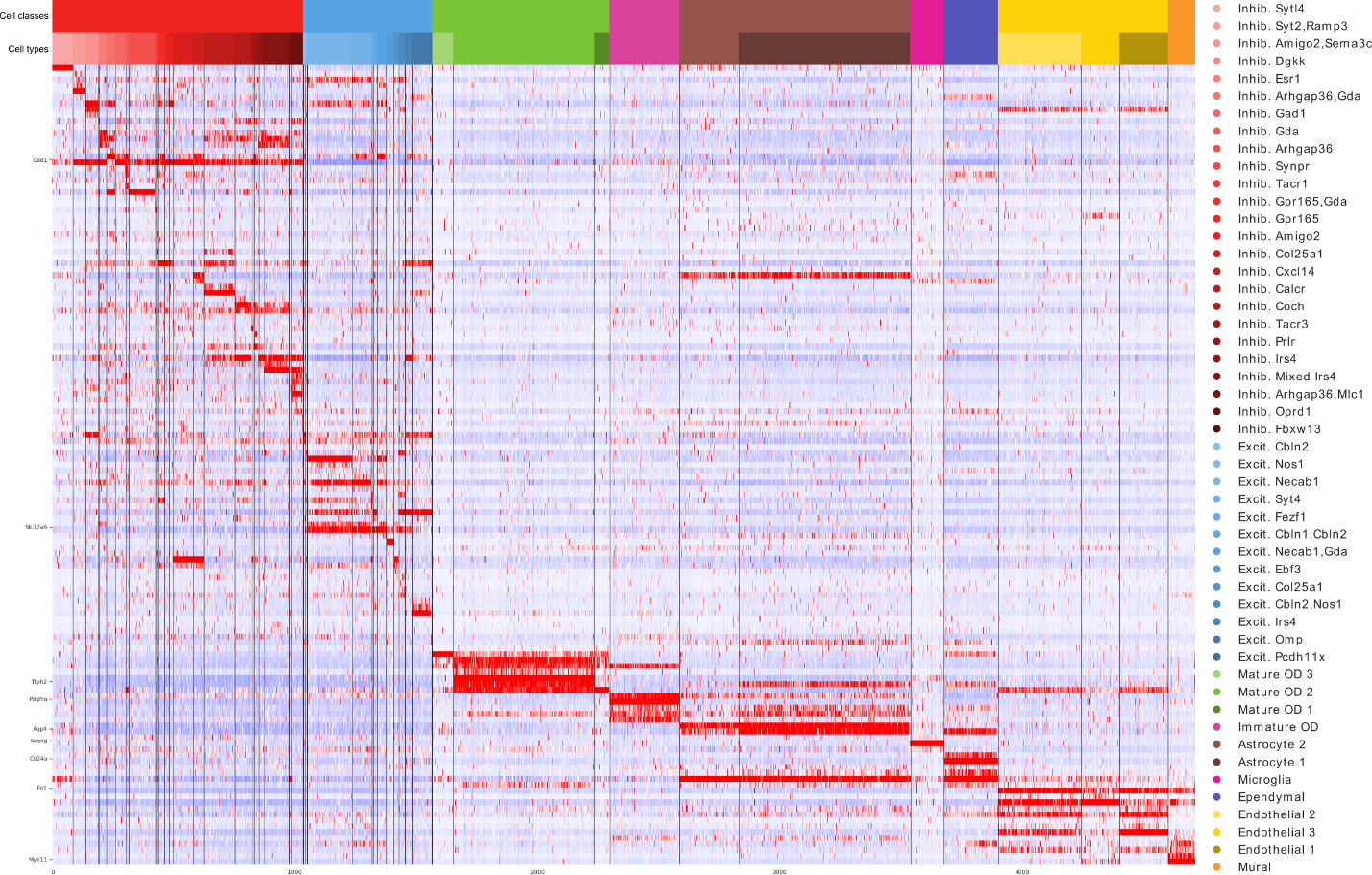

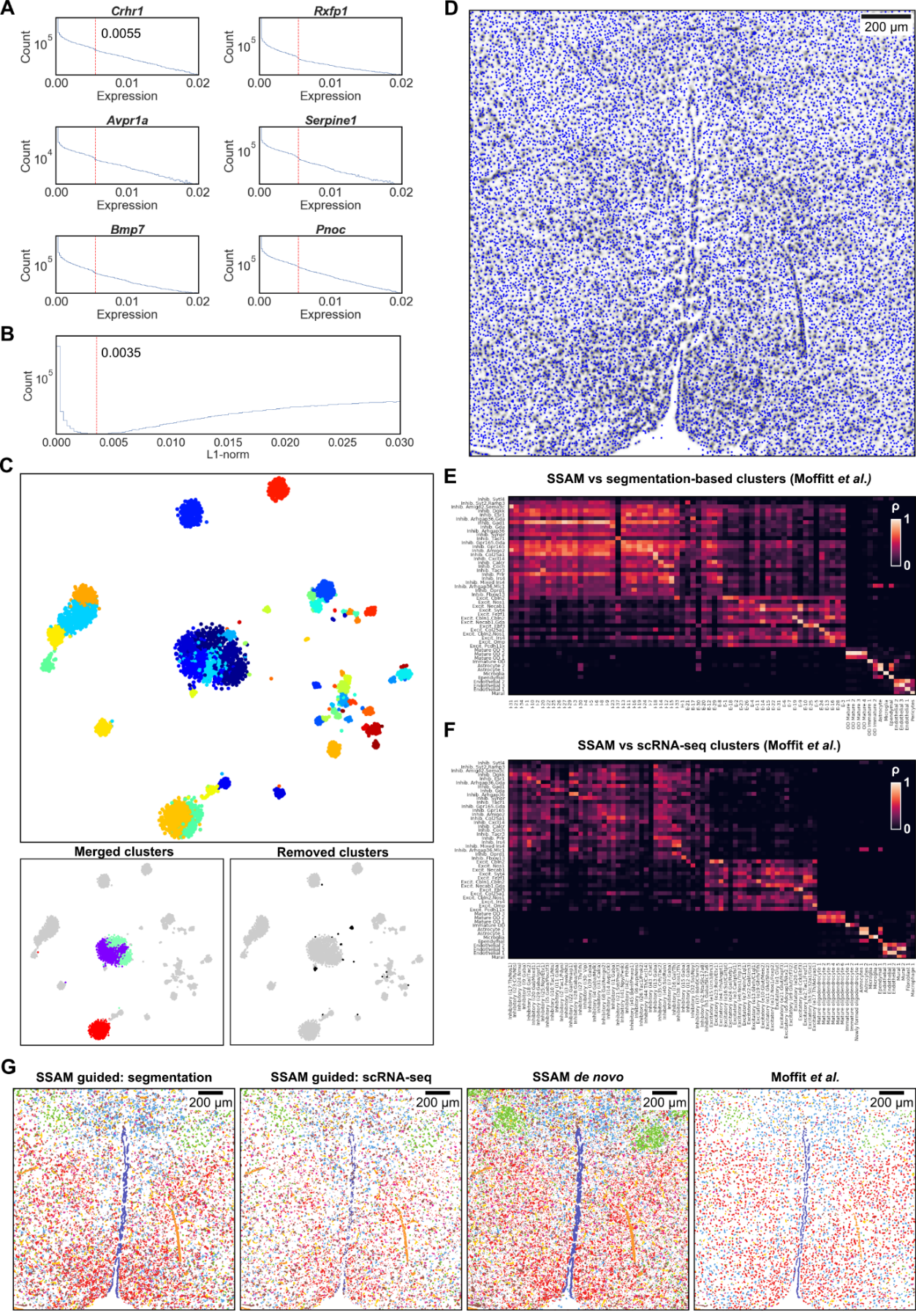

### Supplementary Figure 17. Local maxima selection, cell-type signature identification, and mapping in the mouse POA MERFISH dataset.

(A) The gene expression threshold (red vertical line) corresponds to an observable drop in expression signal, shown for 6 randomly selected genes from the dataset. Local maxima are only selected if they exceed the determined gene expression threshold for at least 1 gene, so that at least 1 gene is sufficiently expressed.

(B) The total gene expression threshold (red vertical line). Since this determined threshold is larger than the gene expression threshold determined in (A), this threshold was not applied.

(C) Clustering merging and filtering. A tSNE map of clustering all local maxima gene expression vectors (top). Clusters with similar expression profiles were merged (bottom left), and those mapping to artifact regions of the image, or containing highly variable gene signatures were removed (bottom right).

(D) Selected local maxima on the vector field.

(E) Comparison of the SSAM *de novo* cell-type signatures to the segmentation-based cell-type signatures from Moffitt *et al.*

(F) Comparison of the SSAM *de novo* cell-type signatures to the scRNA-seq derived cell-type signatures from Moffitt *et al.*

(G) Comparison of cell-type maps generated by SSAM guided and *de novo* mode. Cell- type maps generated using signatures derived from (left to right) segmentation and scRNAseq from Moffitt *et al.*, SSAM *de novo* signature identification, and the original cell map from Moffitt *et al.* Color of scRNA-seq clusters were determined by calculating Pearson’s correlation coefficient between scRNA-seq and segmentation-based cluster centroids. The colors of the cell types correspond to the cell-type legend in Fig. 4B.

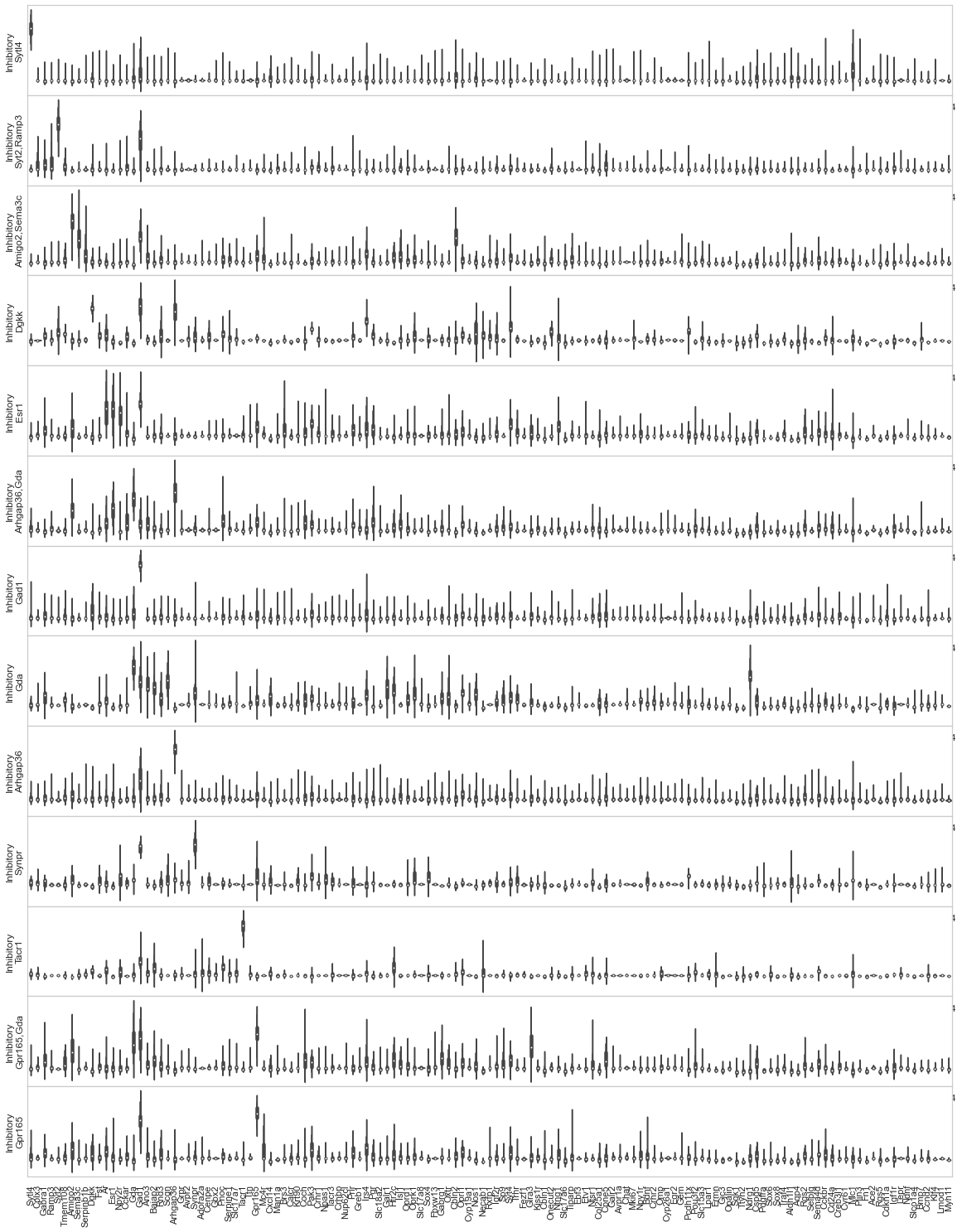

### Supplementary Figure 18. Gene expression violin plots of local maxima, MERFISH inhibitory neuronal cell types (part 1).

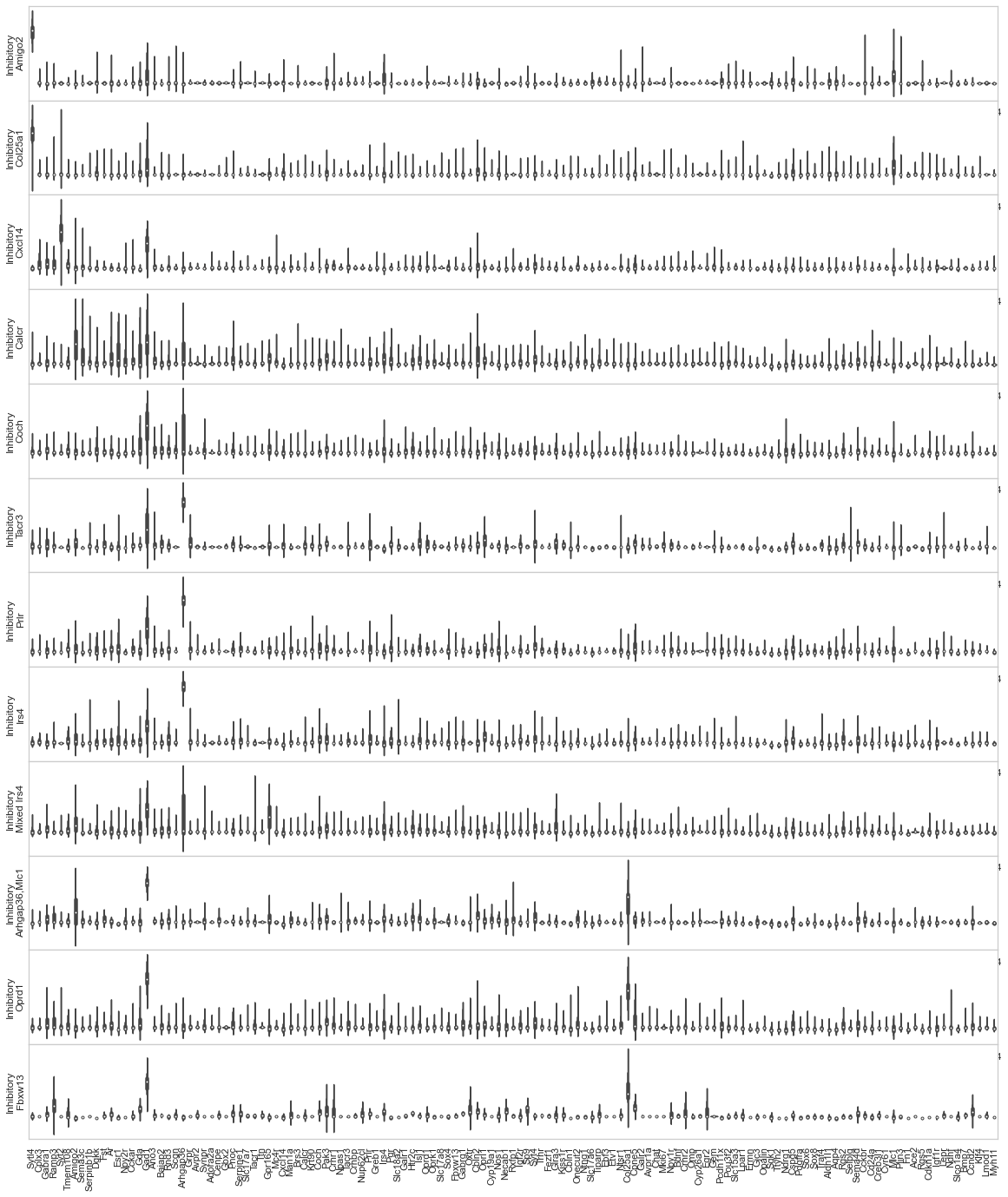

### Supplementary Figure 19. Gene expression violin plots of local maxima, MERFISH inhibitory neuronal cell types (part 2).

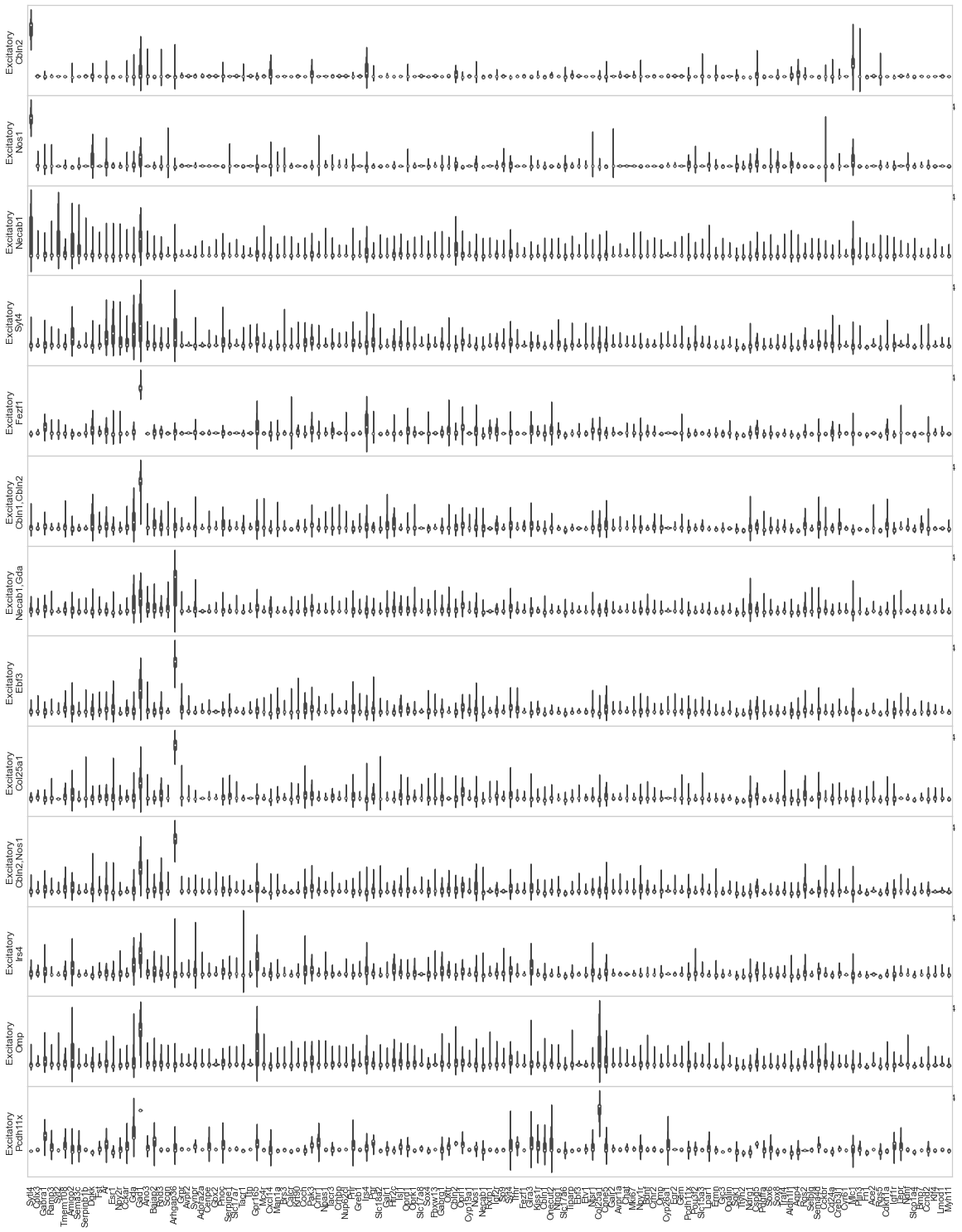

### Supplementary Figure 20. Gene expression violin plots of local maxima, MERFISH excitatory neuronal cell types.

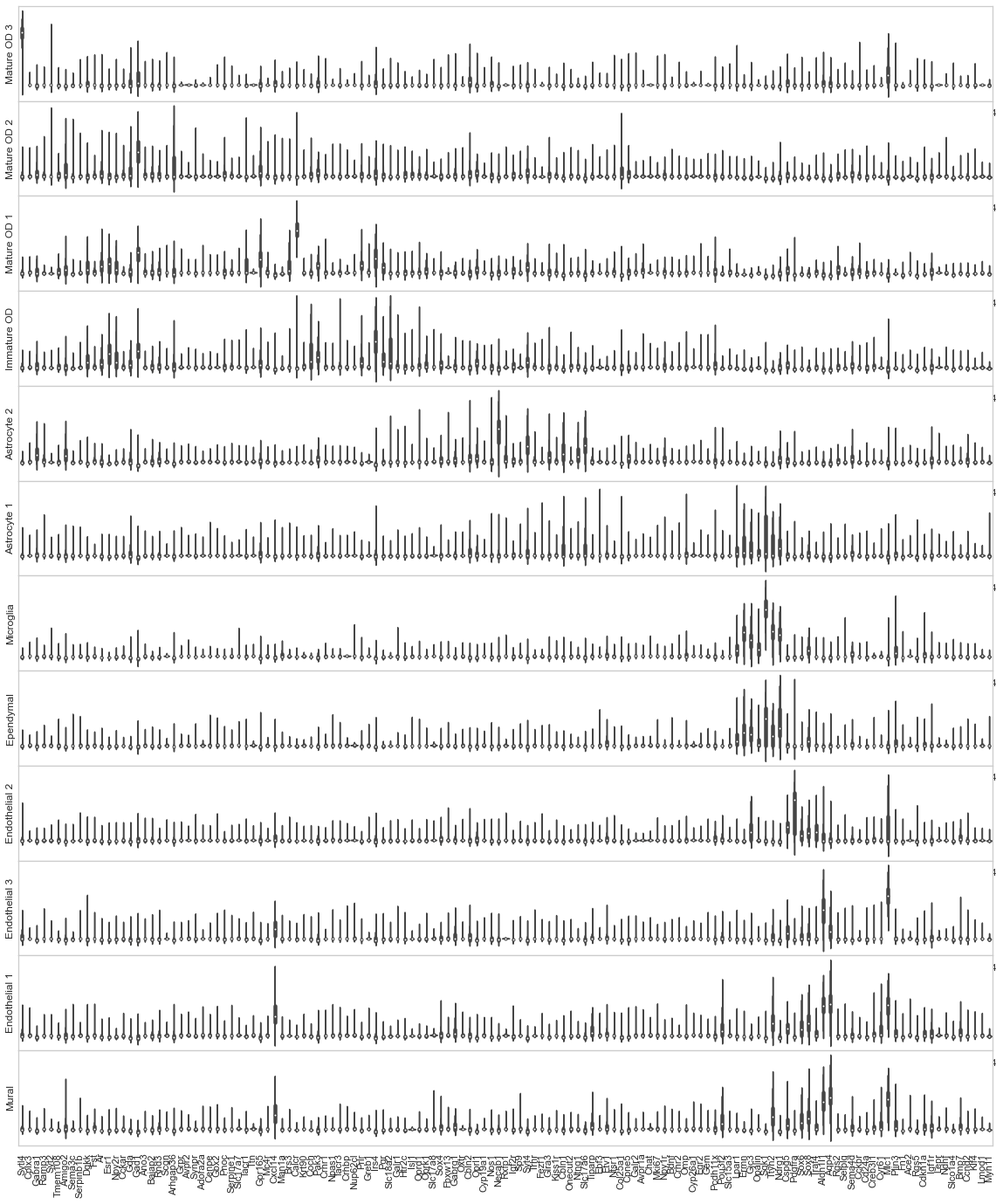

### Supplementary Figure 21. Gene expression violin plots of local maxima, MERFISH other cell types.

### Supplementary Figure 22. Comparison of spatial localization patterns of inhibitory and excitatory cell-types in the mouse POA MERFISH dataset.

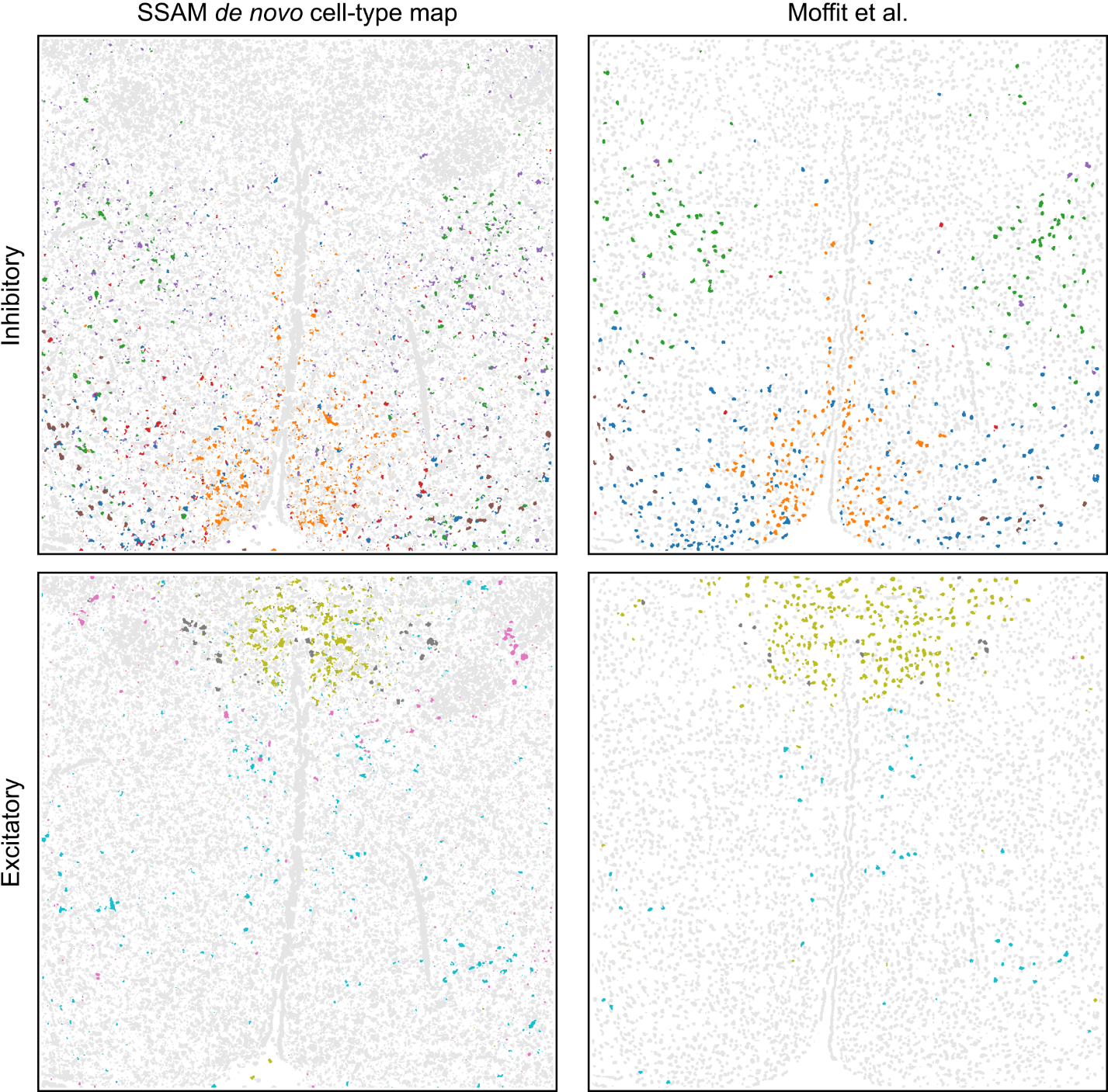

A number of inhibitory (top row) and excitatory (bottom row) cell types were found to have similar tissue localization patterns between the SSAM *de novo* cell-type map (left column) and the segmentation-based cell-type map (right column). The cell type legends can be found in Figure 4.

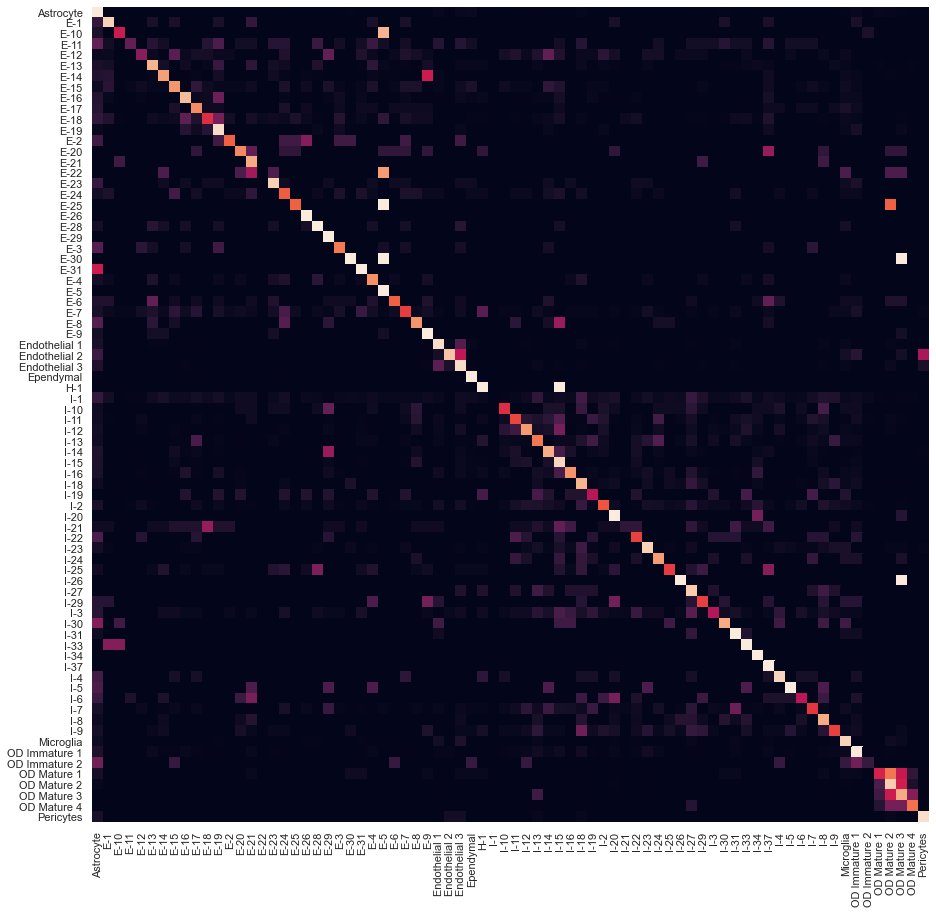

### Supplementary Figure 23. Pairwise matching scores heatmap between the guided mode cell type map (MERFISH segmentation based) and MERFISH segments.

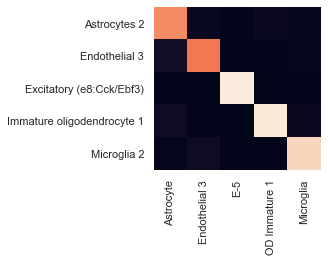

### Supplementary Figure 24. Pairwise matching scores heatmap between the guided mode cell type map (MERFISH scRNA-seq based) and MERFISH segments.

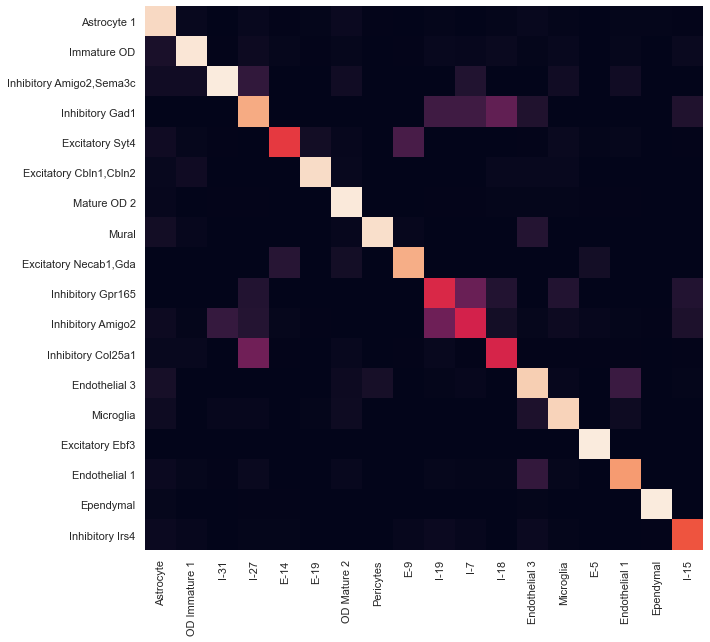

### Supplementary Figure 25. Pairwise matching scores heatmap between the *de novo* cell type map and MERFISH segments.

### Supplementary Figure 26. Comparison of Astrocytes identified by SSAM and segmentation vs Aldh1l1 expression in the mouse POA MERFISH dataset.

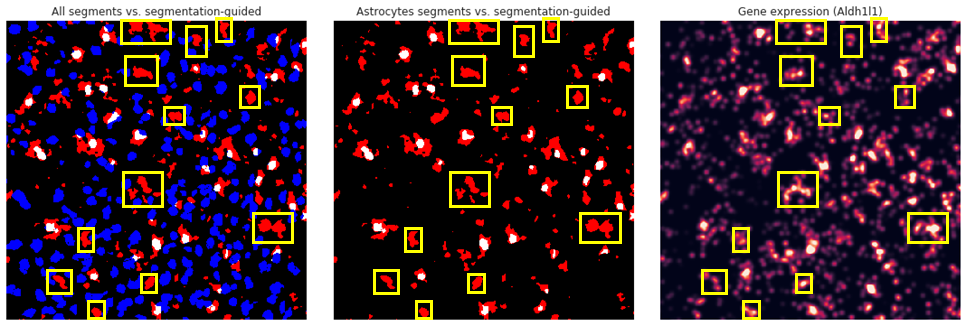

(Left) SSAM guided-mode cell type is colored in red, the segments of all cell types by Moffit et al. are colored in blue, and the overlap is colored in white. (Right) Gene expression of Aldh1l1. Although Aldh1l1 is not a specific marker for Astrocytes, it is one of the genes highly expressed in Astrocytes, which confirms the existence of Astrocytes uniquely identified by SSAM (highlighted in yellow).

### Supplementary Figure 27. SSAM identifies enriched inhibitory and excitatory tissue domains in the posterior hypothalamic POA.

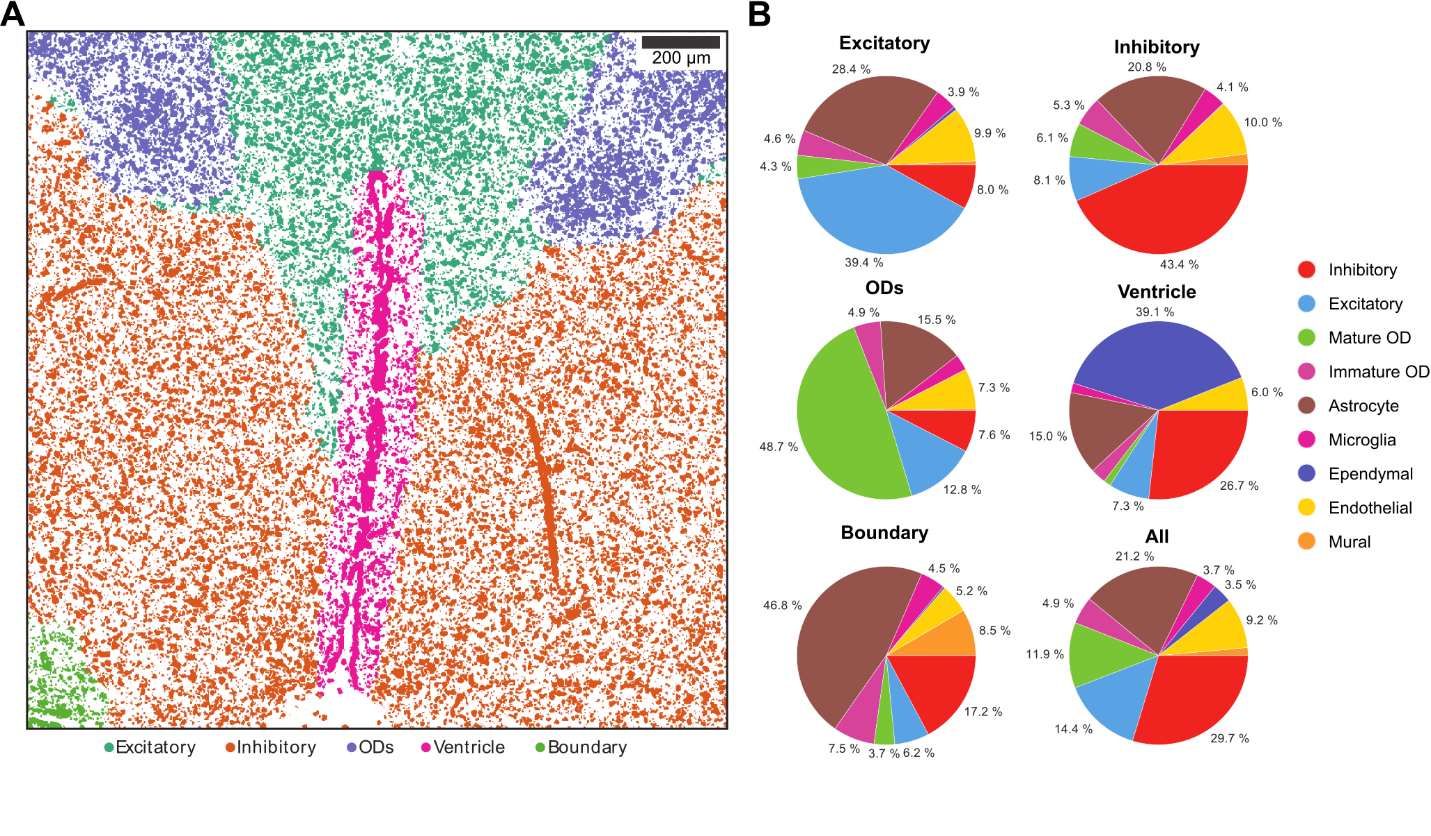

(A) Tissue domain map generated by SSAM. Tissue domain signatures were identified from clustering local cell-type composition over sliding 100 μm circular windows, and projected back onto the cell-type map. The ventricle was manually removed from the tissue domain reconstruction. The reconstruction shows while the distribution of inhibitory and excitatory regions is intermingled, there are domains with enrichment of either of these cell types.

(B) Cell-type composition within each tissue domain. The plots shows the composition ratio of approximately 5:1 of inhibitory to excitatory cell types in the inhibitory tissue domain and vice versa. The colors in the pie charts correspond to the cell-type legend in Fig. 4B.

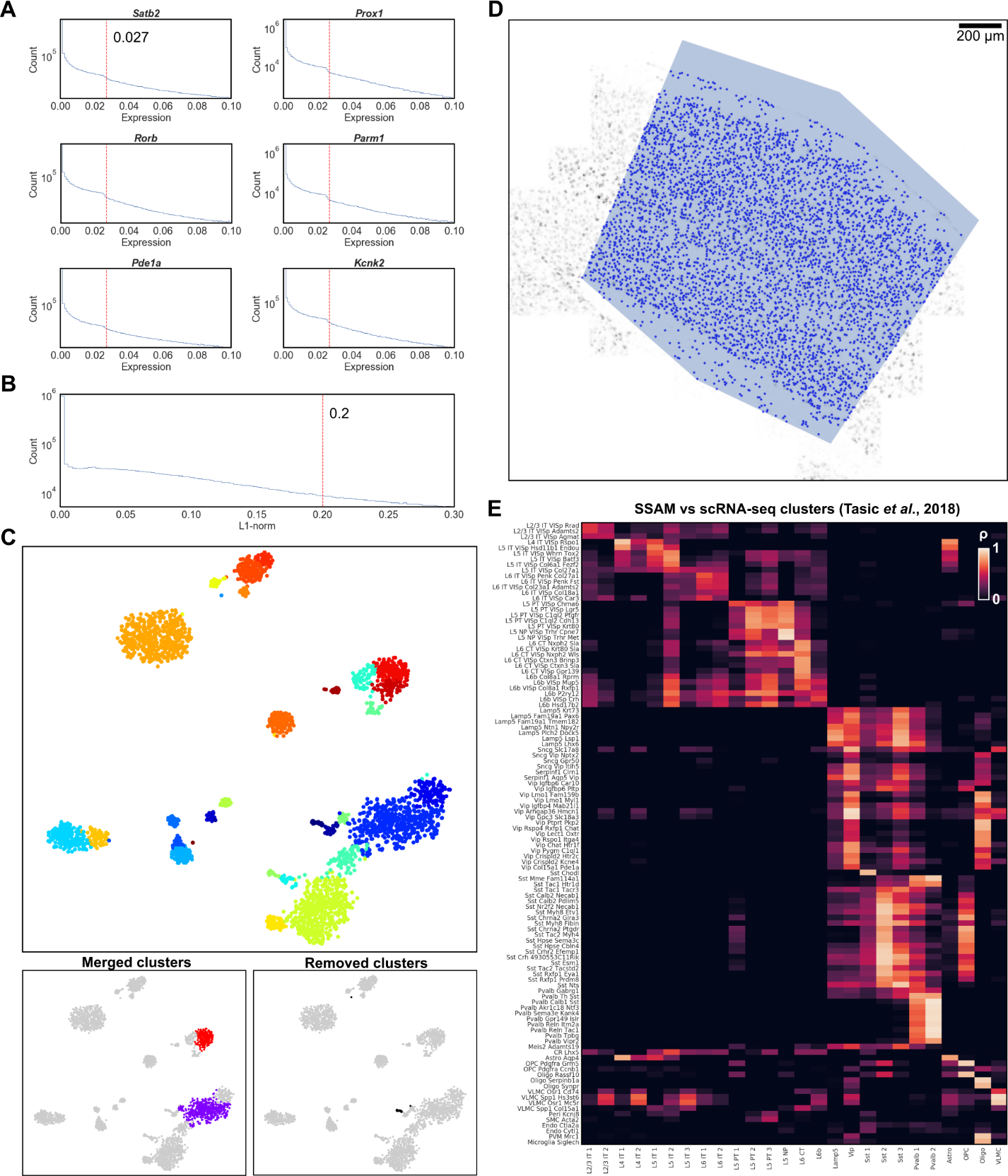

### Supplementary Figure 28. Local maxima selection and cell-type signature identification in the mouse VISp smFISH dataset.

(A) The gene expression threshold (red vertical line) corresponds to an observable drop in expression signal, shown for 6 randomly selected genes from the dataset. Local maxima are only selected if they exceed the determined gene expression threshold for at least 1 gene, so that at least 1 gene is sufficiently expressed.

(B) The total gene expression threshold (red vertical line). Local maxima are only selected if they exceed the determined total gene expression threshold, preventing selection of local maxima with a low total gene expression signal.

(C) Clustering merging and filtering. A tSNE map of clustering all local maxima gene expression vectors (top). Clusters with similar expression profiles were merged (bottom left), and those mapping to artifact regions of the image, or containing highly variable gene signatures were removed (bottom right).

(D) Selected local maxima on the vector field. The VISp region is supplied as in input mask, represented as a polygon on the image, so that SSAM only selects local maxima in the VISp region.

(E) Comparison of the SSAM *de novo* cell-type signatures to the scRNA-seq derived cell-type signatures from Tasic *et al* (2018).

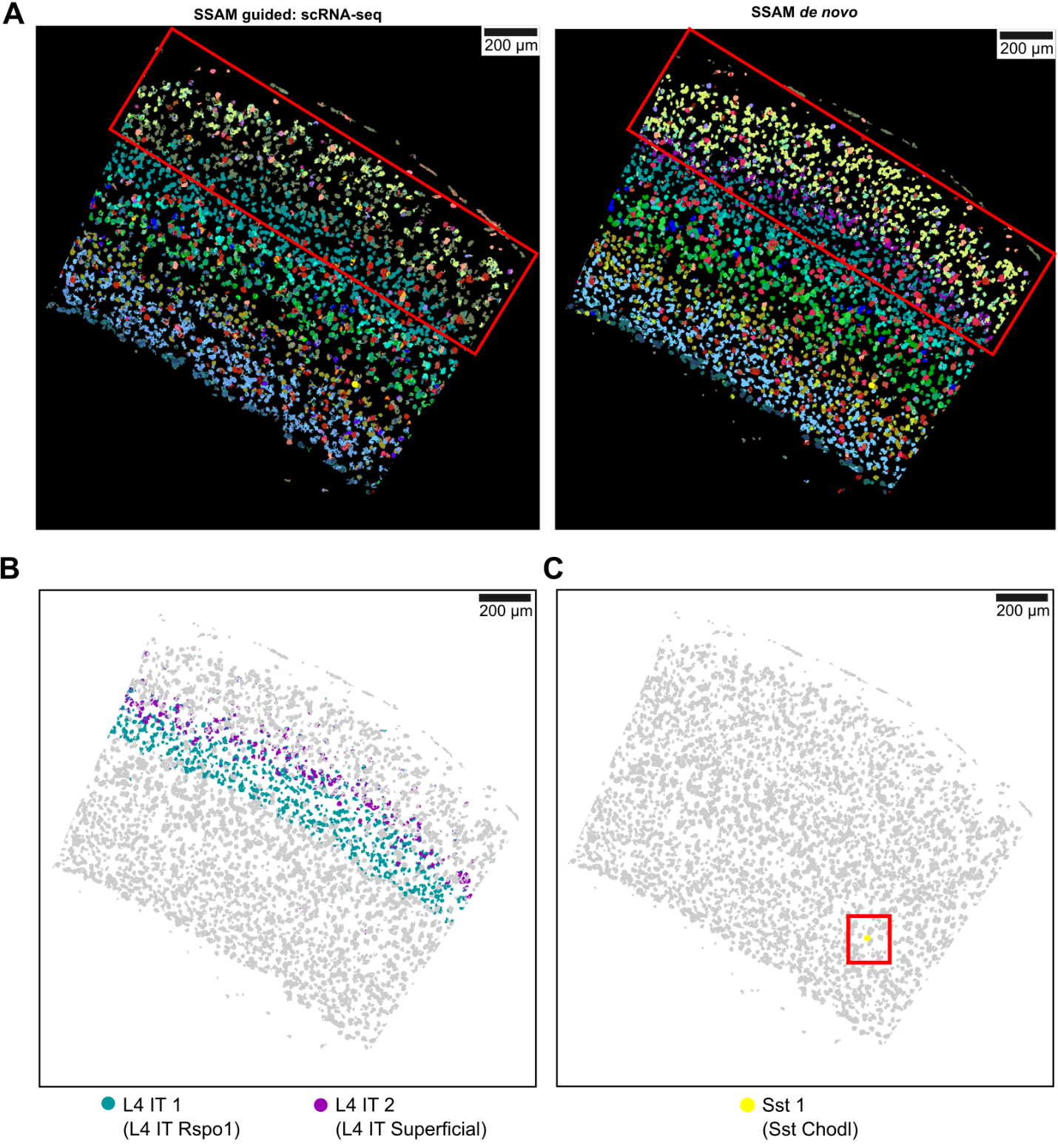

### Supplementary Figure 29. Identifying new sub-layering in the L4 cortical layer in the mouse VISp smFISH dataset.

(A) Comparison of SSAM guided mode and *de novo* mode results. The two results are visually similar apart from the L2/3 region (boxed in red), which shows wrongly mapped cell types in the guided mode result, indicating that the guided mode does not always guarantee the best results. Instead the *de novo* mode can be used to find accurate mapping of cell types on tissue when guided mode fails to reconstruct cell-type maps correctly.

(B) Localization of the two L4 IT sub-layering. The heterogeneity of L4 IT cell type is related with their localizations in layer 4 of the VISp, with cell type L4 IT1 localizing to the deep portion, and L4 IT2 localizing superficial.

(C) SSAM was able to find the rare Sst Chodl cell type (boxed in red), which shows the sensitivity of the cell type detection in *de novo* mode.

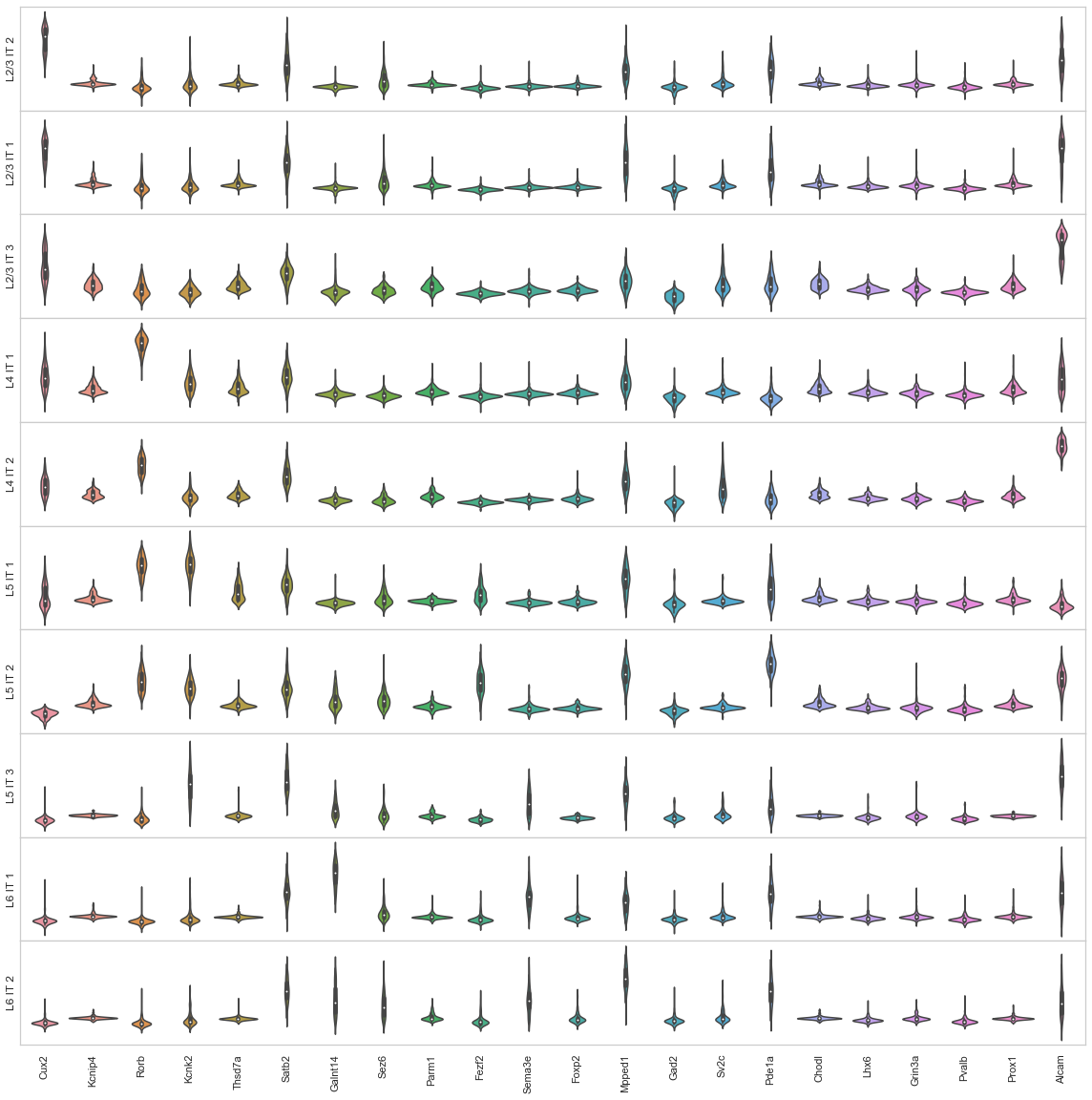

### Supplementary Figure 30. Gene expression violin plots of local maxima, multiplexed smFISH cell types (part 1).

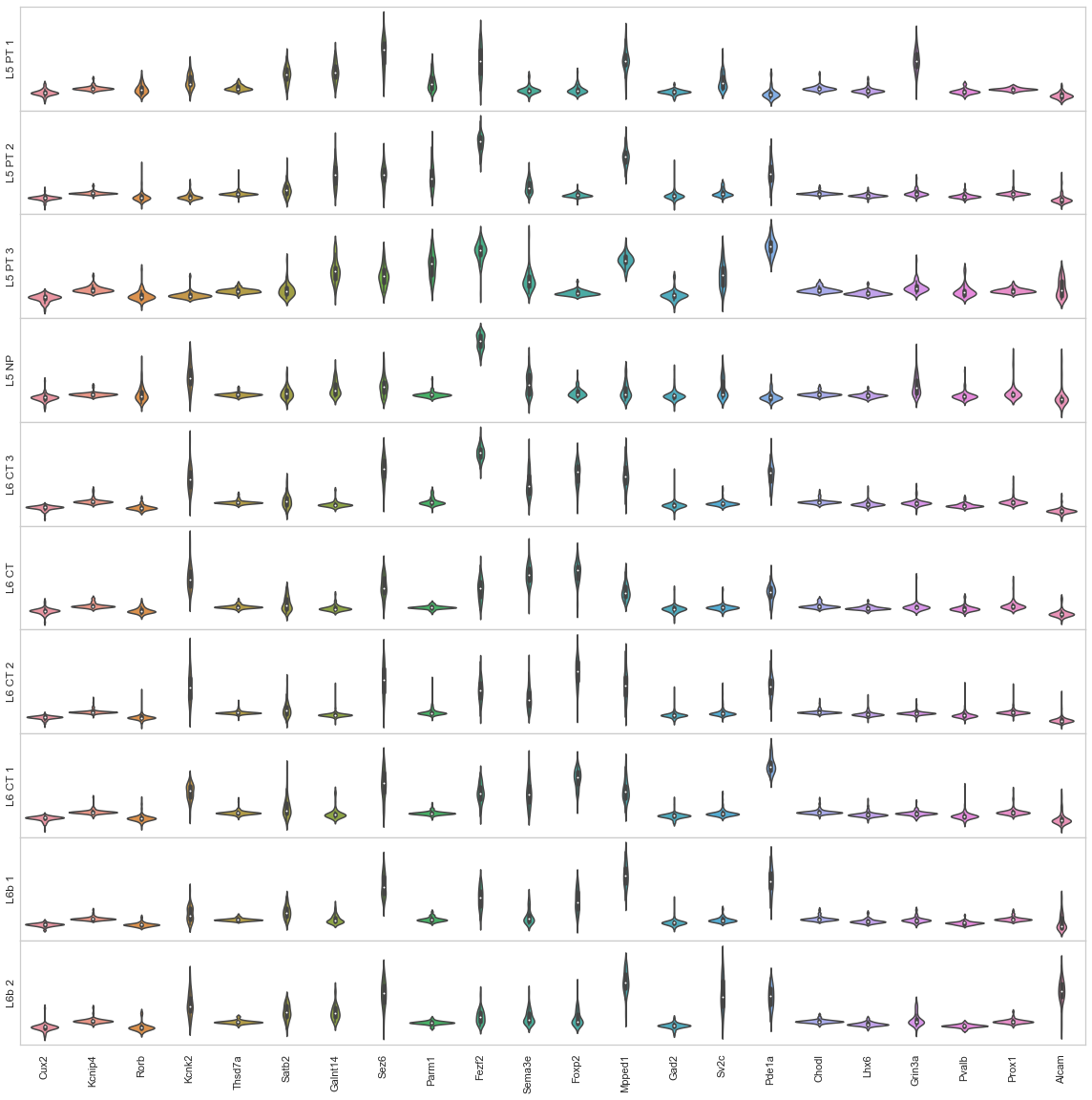

### Supplementary Figure 31. Gene expression violin plots of local maxima, multiplexed smFISH cell types (part 2).

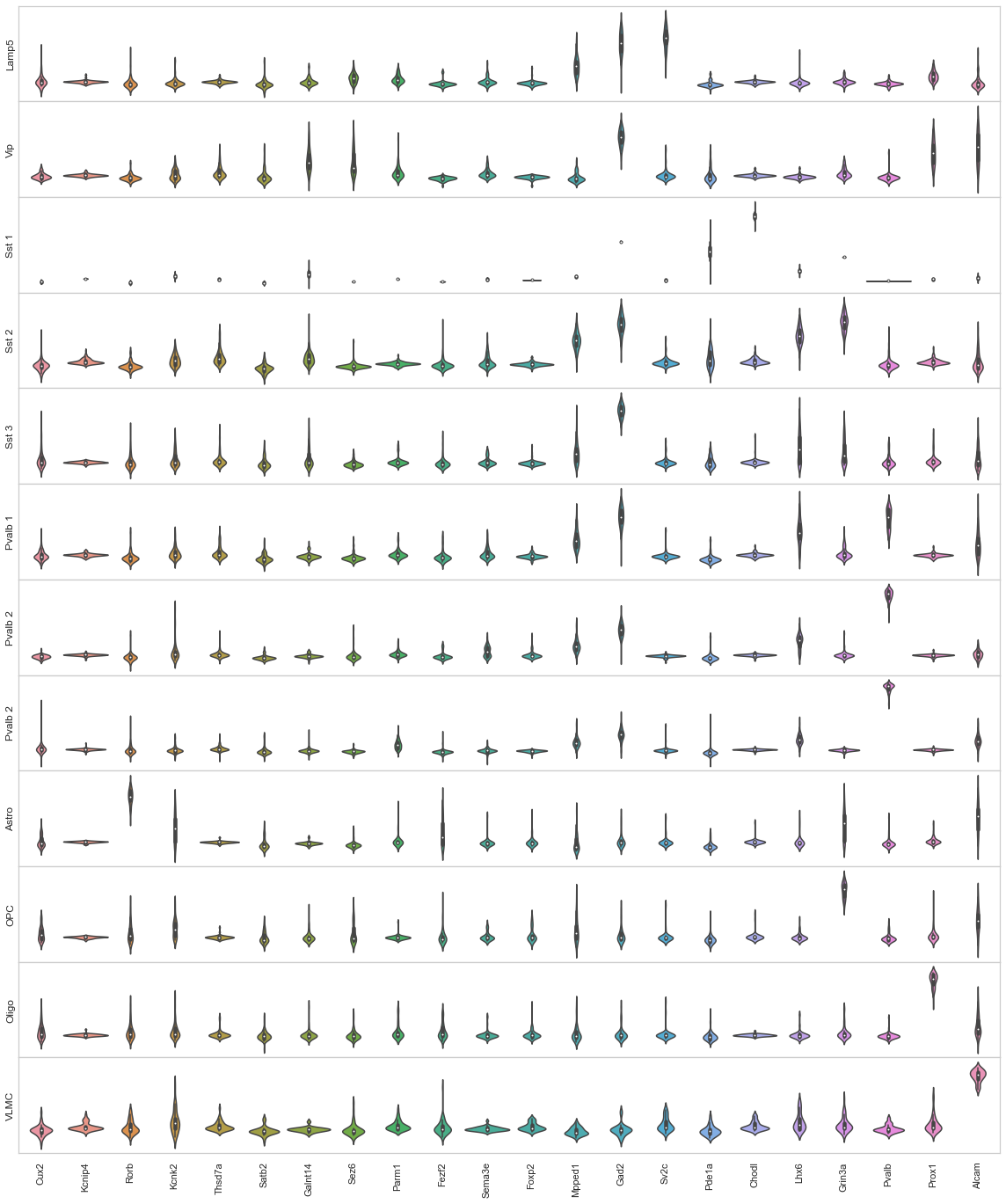

### Supplementary Figure 32. Gene expression violin plots of local maxima, multiplexed smFISH cell types (part 3).

### Supplementary Figures 33-36. Effect of KDE bandwidth and lattice spacing in 4 different regions.

The figures show the cell-type map of 4 different regions (highlighted in white boxes), generated with 5 different KDE bandwidth (0.5, 1, 2.5, 5, 10 um) and 3 different lattice spacing (0.5, 1, 2.5 um) in guided-mode SSAM (guided by segmentation-based signatures) using osmFISH dataset. The figure shows that only the bandwidth is related to the size of detected blobs cell-type map, and the lattice spacing is only related to its pixel size. For all cases, even including very extreme cases with bandwidths 0.5 and 10 um, the pixels in the cell-type map correlates well with the corresponding cell-type signatures and colored correctly. In other words, the signals estimated by KDE preserves most of the cell-type signatures even with extreme change of bandwidth, which implies that the cell-type identification will not be largely affected by a small change of bandwidth near 2.5 um.

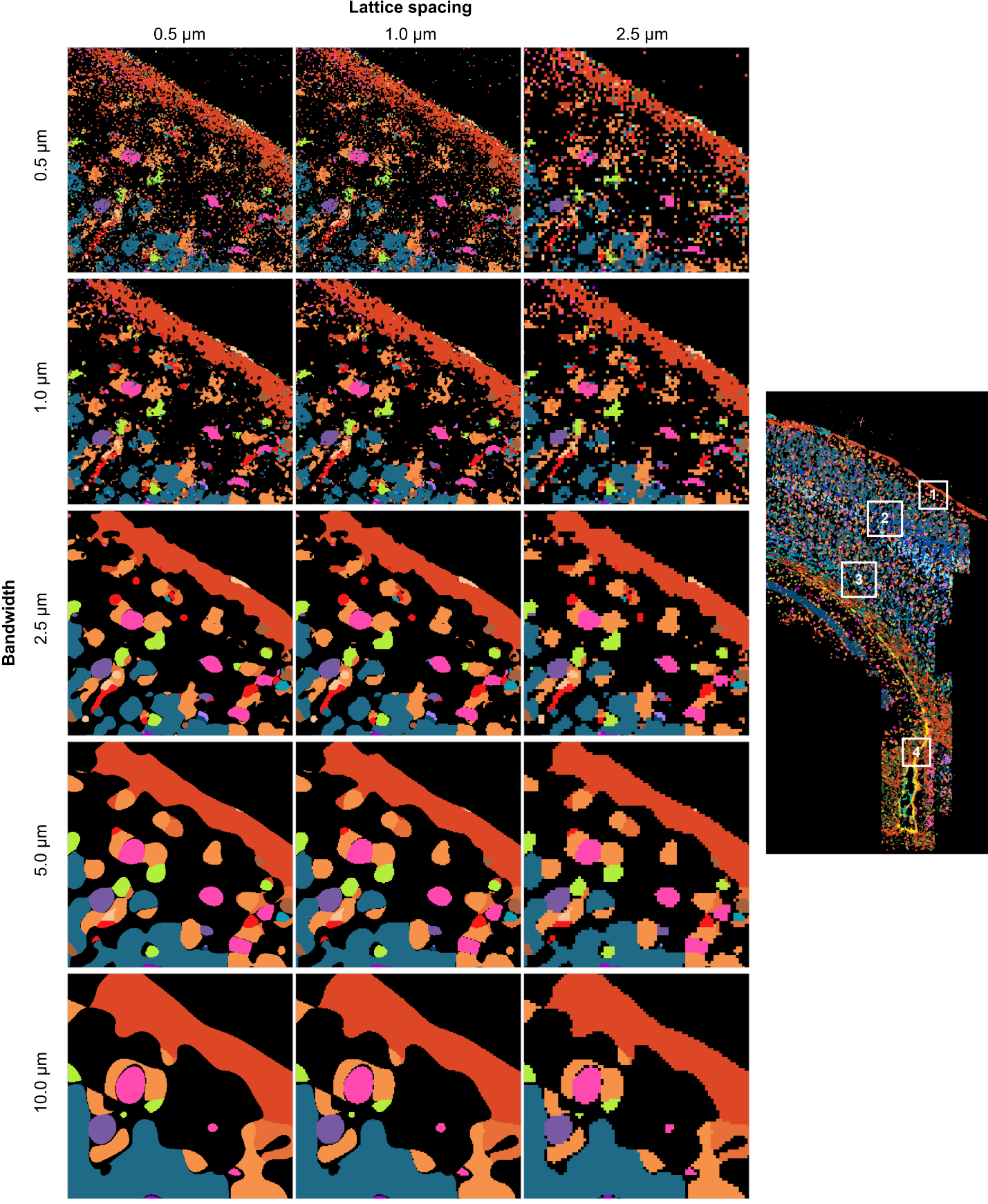

### Supplementary Figure 33. Effect of KDE bandwidth and lattice spacing in the first region.

### Supplementary Figure 34. Effect of KDE bandwidth and lattice spacing in the second region.

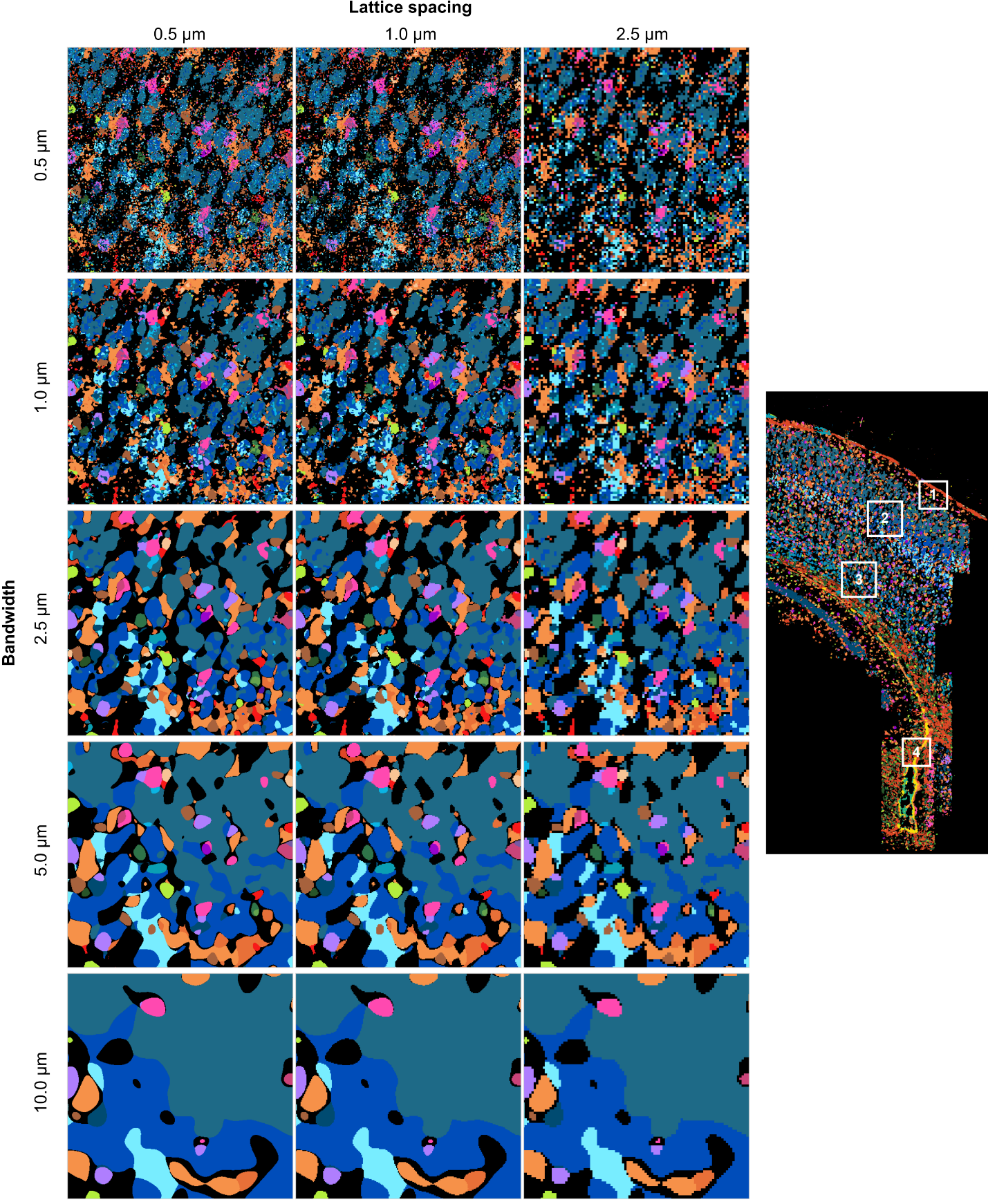

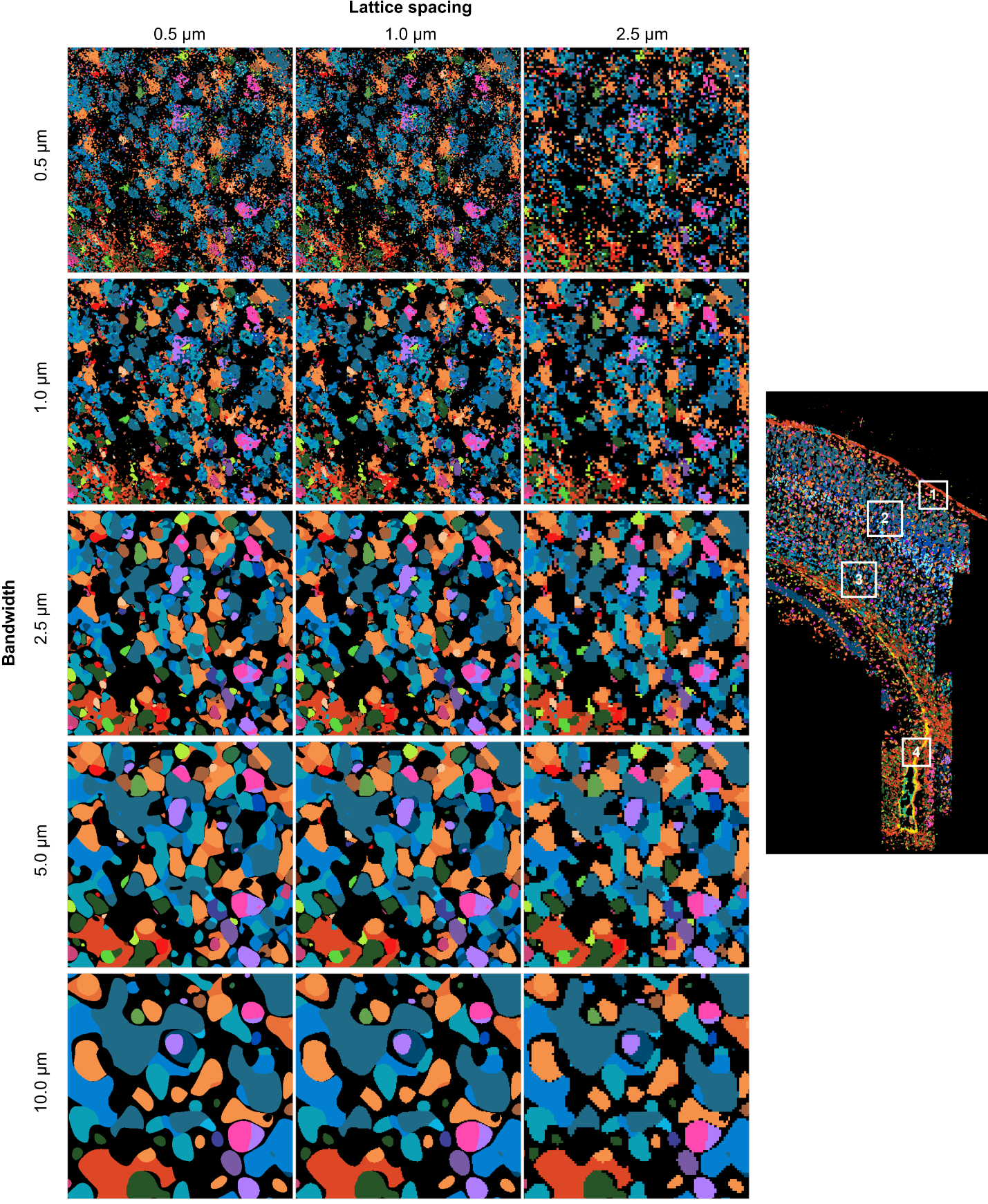

### Supplementary Figure 35. Effect of KDE bandwidth and lattice spacing in the third region.

### Supplementary Figure 36. Effect of KDE bandwidth and lattice spacing in the fourth region.

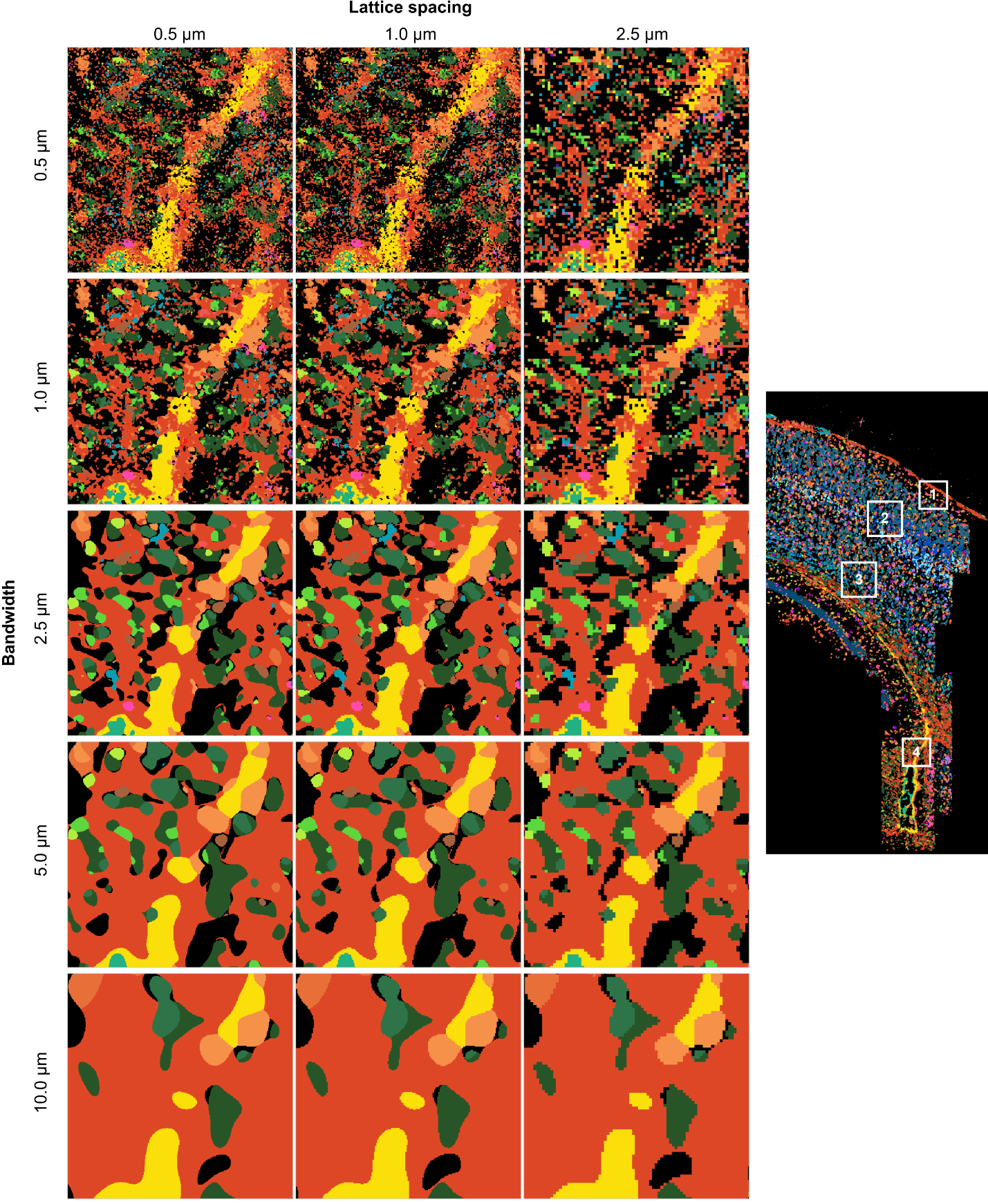

### Supplementary Figure 37. Correlations between the *de novo* clusters with bandwidth 2.5 and the unmerged clusters with different bandwidths.

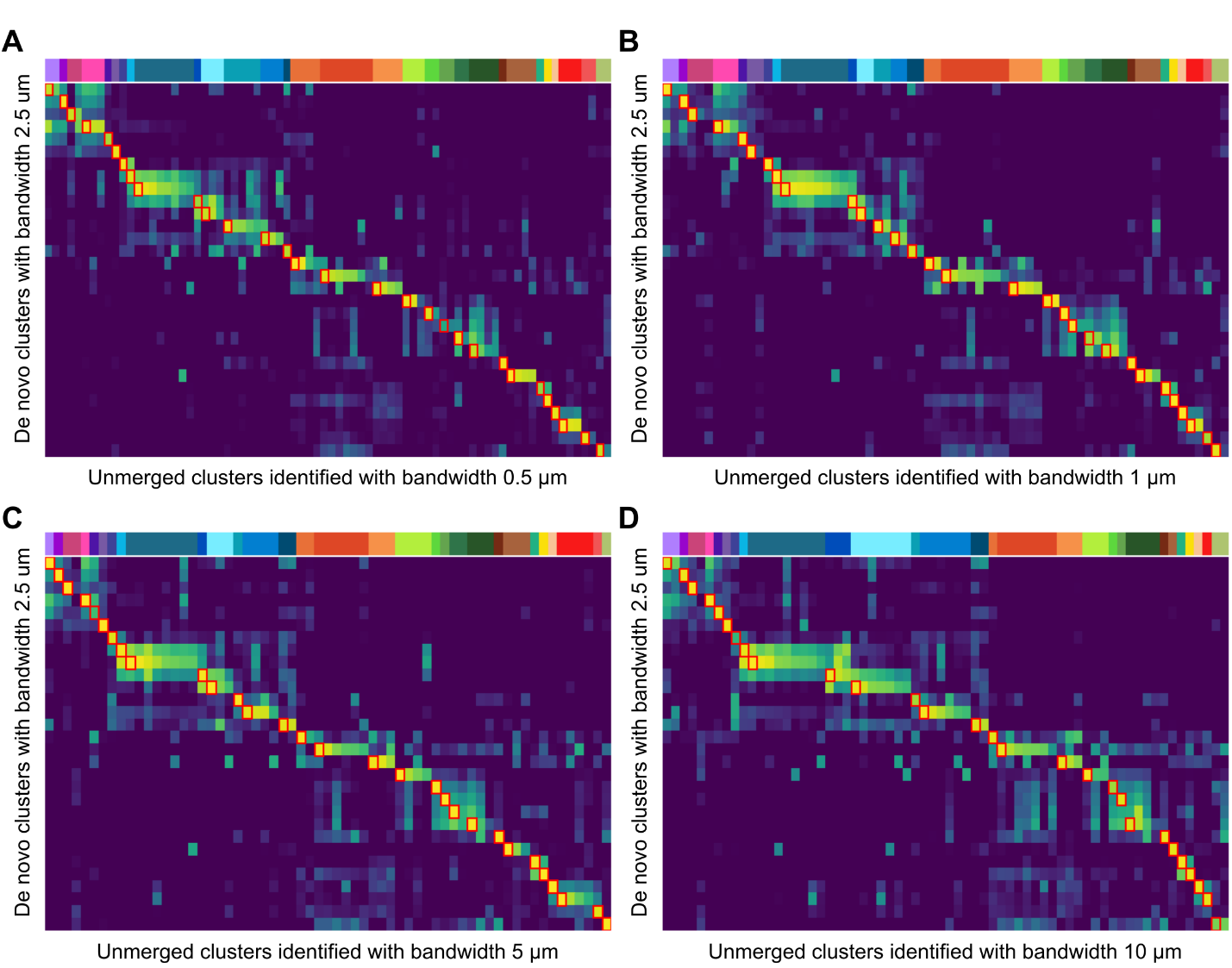

(All panels) Pearson’s correlation between the clusters with bandwidth 2.5 and the unmerged clusters identified with different bandwidths, 0.5, 1, 5, and 10. The representative vectors were selected at the same locations identified in the original *de novo* analysis. The highest correlated clusters for each cluster identified in the original *de novo* analysis are marked with red boxes. (A) Among the unmerged 76 clusters with bandwidth 0.5, all the *de novo* clusters were highest correlated with at least one of them. The highest correlated values in red boxes were with max 0.997, min 0.607, and median 0.961. (B) Among the unmerged 67 clusters with bandwidth 1, all the *de novo* clusters were highest correlated with at least one of them. The highest correlated values in red boxes were with max 0.998, min 0.861, and median 0.990. (C) Among the unmerged 63 clusters with bandwidth 5, all the *de novo* clusters were highest correlated with at least one of them. The highest correlated values in red boxes were with max 0.998, min 0.690, and median 0.993. (D) Among the unmerged 66 clusters with bandwidth 10, except 2 clusters, all the other *de novo* clusters were highest correlated with at least one of them. The highest correlated values in red boxes were with max 0.993, min 0.658, and median 0.968.

### Supplementary Figure 38. Comparison of representative vector selection based on local maxima and random selection.

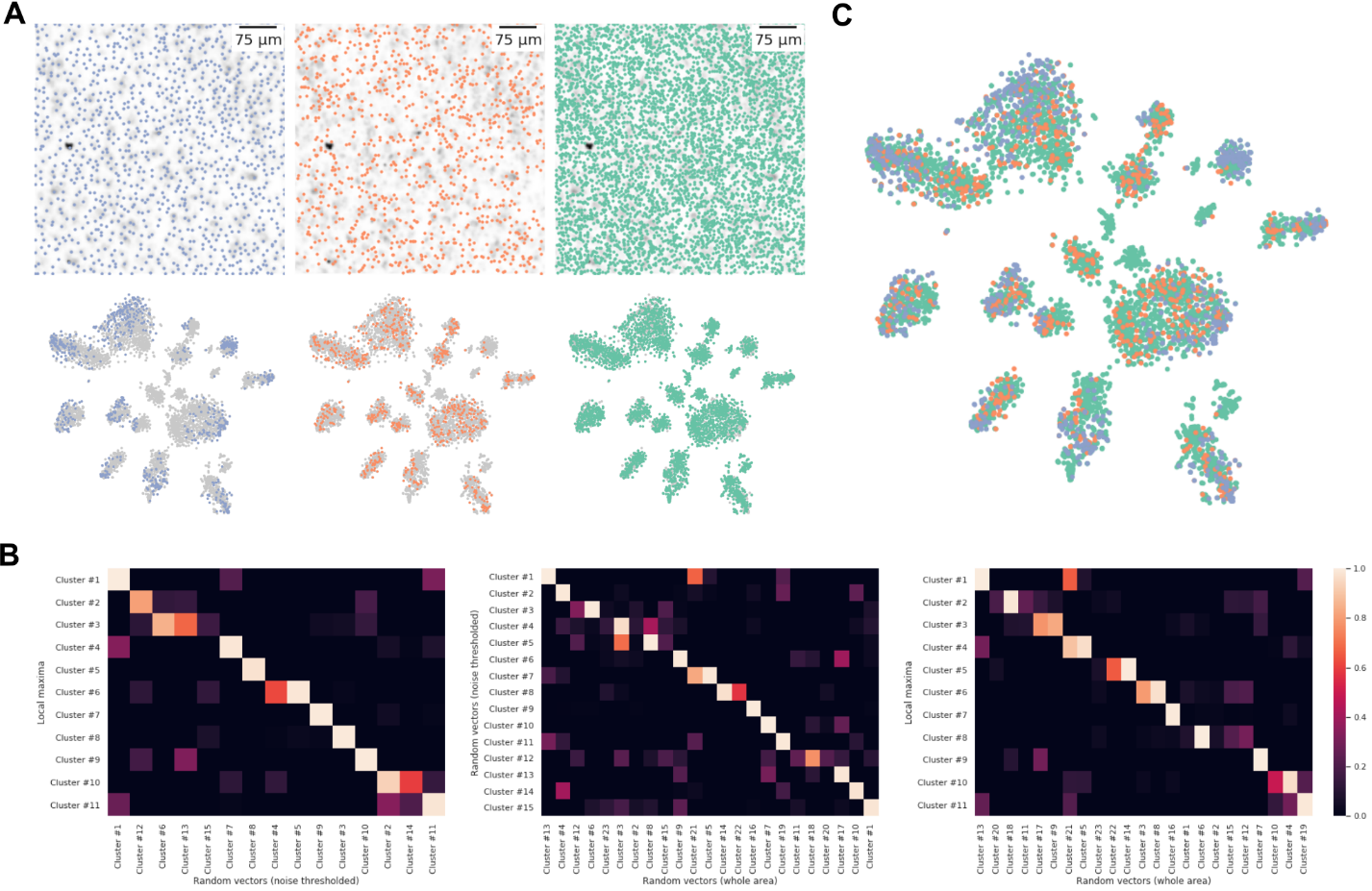

(A) Selected vectors of 1,056 local maxima (blue), 1,000 randomly selected locations above noise thresholds based on mRNA signal strength (orange), 5,000 randomly selected locations without thresholding (teal), and their corresponding 2-dimensional tSNE embeddings. (B) The pairwise correlation heatmaps between the cluster centroids of each case. Regardless of the selection strategy of vectors, still there is at least one highly correlated cluster between each other. (C) Combined tSNE embedding of the three different methods of vector selection.

### Supplementary Figure 39. Comparison of local maxima and random sampling in the osmFISH dataset shows local maxima better represent known gene expression profiles.

We compared the representation of cell-type gene expression signatures between local maxima selection and randomly sampled positions against the segmentation derived cell-type gene expression signatures. On average, the local maxima selection resulted in vectors having an average correlation of 0.971 (vertical red line). This was markedly higher when compared against 100 permutations of sampling random pixels equal to the number of local maxima (distribution shown as a blue histogram).

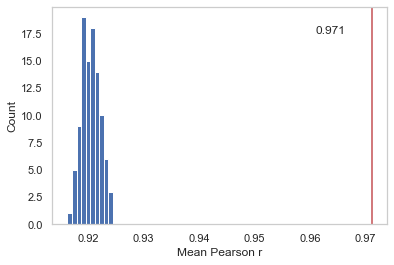

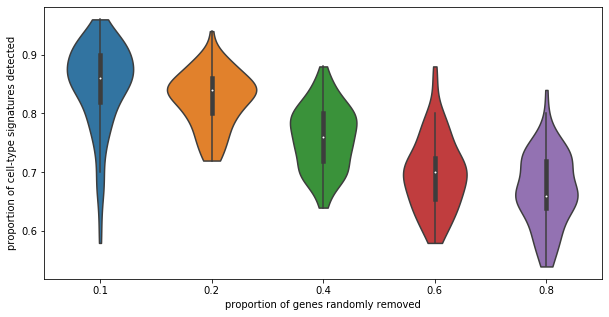

### Supplementary Figure 40. Cell-type detection rate as a function of number of genes using simulated downsampling of MERFISH data.

Benchmarking the cell-type detection rate whereby a percentage of genes were randomly removed from MERFISH data (100 iterations). X axis shows the proportion of gene removed, 10%, 20%, 40%, 60%, and 80%; and Y axis shows the proportion of cell types detected by *de novo* analysis. A cell type was determined to be identifiable if any cluster exceeded a cell-type specific correlation threshold (based on the highest observed non-identity correlation among all cell types identified in the full dataset).

### Supplementary Video Legends

**Supplementary Video 1. MERFISH 3D cell-type map, turntable rotating**

**Supplementary Video 2. MERFISH 3D cell-type map, sweeping along z axis by 1 μm**

**Supplementary Video 3. MERFISH neuronal cells, sweeping along z axis by 1 μm**

**Supplementary Video 4. MERFISH astrocytes, sweeping along z axis by 1 μm**
